# Paired CRISPR screens identify mitochondrial metabolism and UBE2H as aneuploid-specific dependencies in human cancer cell lines

**DOI:** 10.64898/2026.04.26.720636

**Authors:** Klaske M. Schukken, Saron M. Akalu, Charles Zou, Pranav K. Kandikuppa, Ryan A. Hagenson, Jessica L. Keane, Mason P. Lynch, Toyoki Yoshimoto, Olaf Klingbeil, Erin L. Sausville, Sanat Mishra, Christopher R. Vakoc, Zuzana Storchova, Sarah J. Aitken, Jason M. Sheltzer

## Abstract

Aneuploidy is a hallmark of cancer and imposes widespread cellular stress, including proteotoxicity, transcriptional dysregulation, and increased metabolic demand. Although these stresses are predicted to create therapeutic vulnerabilities, the genetic dependencies of aneuploid cells remain incompletely characterized. Here, we performed paired CRISPR loss-of-function screens in isogenic aneuploid and near-euploid cancer cell line models to systematically identify aneuploidy-specific dependencies. Seven genome-wide paired screens identified ribosomes, rRNA processing, spliceosome-mediated RNA processing, proteasome subunits, and mitochondrial metabolism as top aneuploid-specific dependency gene groups. To identify therapeutically targetable aneuploid dependencies, we performed 18 additional paired CRISPR screens using a focused druggable genome library. This analysis identified the ubiquitin-conjugating enzyme UBE2H as a top aneuploid-selective dependency. Functional validation confirmed aneuploid cell dependency on UBE2H, and mechanistic analyses linked UBE2H to mitochondrial protein abundance, suggesting a role in maintaining mitochondrial proteostasis under aneuploid stress. Together, these findings define core cellular systems that support the viability of aneuploid cells and identify UBE2H as a potential therapeutic vulnerability connecting ubiquitin signaling to mitochondrial homeostasis.

## Introduction

Aneuploidy – whole chromosome or arm-level chromosome DNA copy number alterations – occurs in approximately 90% of solid tumors^1^ and is a hallmark of cancer ^2,3^. Aneuploidy has been shown to promote drug resistance^4,5^, and is associated with tumor progression and poor clinical outcome^6–10^. Paradoxically, although aneuploidy is generally detrimental to cell growth^11,12^, cancer cells frequently select for specific chromosome gains and losses containing oncogenes or tumor suppressor genes^13^, respectively, which are ultimately beneficial to cell growth and survival despite global fitness costs^14–16^. While aneuploid cells experience protein-level dosage compensation, chromosome gains and losses can simultaneously alter the abundance of hundreds or thousands of proteins^17,18^. As a consequence, aneuploid cells have been shown to experience gene dosage-associated stresses, including transcriptomic stress^19–21^, proteotoxic stress^20,22–24^, and reduced ribosomal abundance^17,21,25,26^. In addition, aneuploid cells exhibit increased energy expenditure and metabolic stress^11,12,21,24,26,27^.

The frequent occurrence of aneuploidy in cancer, coupled with its low prevalence in normal tissues^28,29^, creates a window of opportunity for therapeutic strategies that selectively target aneuploid tumors. As proof-of-principle that changes in cellular ploidy are sufficient to induce therapeutically targetable dependencies, previous studies have shown that polyploid cells have increased dependency upon the spindle kinesin KIF18A^30,31^, for proper chromosome alignment in mitosis. This has led to the development of KIF18A inhibitors^32^, some of which are currently undergoing clinical trials in patients with advanced solid tumors (Trial IDs: NCT06084416, NCT07226427, NCT05902988, NCT06772415, NCT07260513).

Aneuploid stressors may impose vulnerabilities that can be therapeutically exploited to selectively target aneuploid cancer cells. Two recent studies using CRISPR screens in paired aneuploid and near-euploid controls have begun to uncover aneuploidy-associated genetic dependencies^27,33^. Magesh et al. (2025)^27^ found an increased dependency on pyrimidine biosynthesis and mitochondrial metabolism genes while Zerbib et al. (2024)^33^ found aneuploid clones were less dependent upon DNA damage response genes and the tumor suppressor TP53 pathway. However, these studies each focused on a single untransformed cell type and its aneuploid subclones. Additionally, these screens were limited to a single method of inducing aneuploidy, and only examined aneuploid subclones with net chromosome gains, potentially limiting the generalizability of their findings.

In this study, we generate over 60 novel aneuploid subclones from near-euploid human cancer cell lines and perform 24 CRISPR screens on aneuploid clones along with their isogenic near-euploid controls. These screens span eight cell types, four cancer types, two CRISPR libraries, include aneuploid subclones generated using three distinct methods of inducing aneuploidy, and include subclones with net chromosome losses and those with net gains. Through this approach, we identify proteasome, spliceosome, ribosome, rRNA processing, and mitochondrial metabolism–associated genes as conserved aneuploid-specific dependencies. We further identify and experimentally validate the E2 ubiquitin-conjugating enzyme UBE2H as a key aneuploid dependency and demonstrate that its loss significantly affects mitochondrial electron transport chain protein abundance.

## Results

### Generating novel aneuploid subclones

To identify aneuploid-specific genetic dependencies in human cancers, we set out to compare aneuploid subclones relative to isogenic near-euploid controls in human cancer cell lines. We screened eight different cell types, seven of which are cancer cell lines, and used three different methods to generate aneuploid subclones. By generating aneuploidy via three independent methods, we sought to uncover potentially therapeutic aneuploid vulnerabilities that are independent of the method by which karyotypes were manipulated.

The first method involved Inducing Chromosome INstability (ICIN) in near-euploid cell lines using the Monopolar Spindle 1 (MPS1) inhibitor AZ3146, followed by single-cell sorting of the resulting heterogeneous aneuploid population to isolate aneuploid subclones (Figure 1A). Inhibiting MPS1 via AZ3146 inhibits the spindle assembly checkpoint, induces chromosome missegregation, and leads to aneuploidy^34,35^. All subclones were karyotyped using low-coverage whole-genome sequencing to identify aneuploid clones. Using this approach, 63 aneuploid clones were generated across six cell lines (A2780, DLD1, RKO, SNU1, SW48, and VACO432). Most (49/63) aneuploid clones that we generated contained one or more chromosome gains. The SNU1 near-tetraploid cell line generated subclones with both gains and losses, as well as subclones with only chromosome losses. As SNU1 is a tetraploid cell line, trisomies represent chromosome losses relative to the basal ploidy.

**Figure 1:**
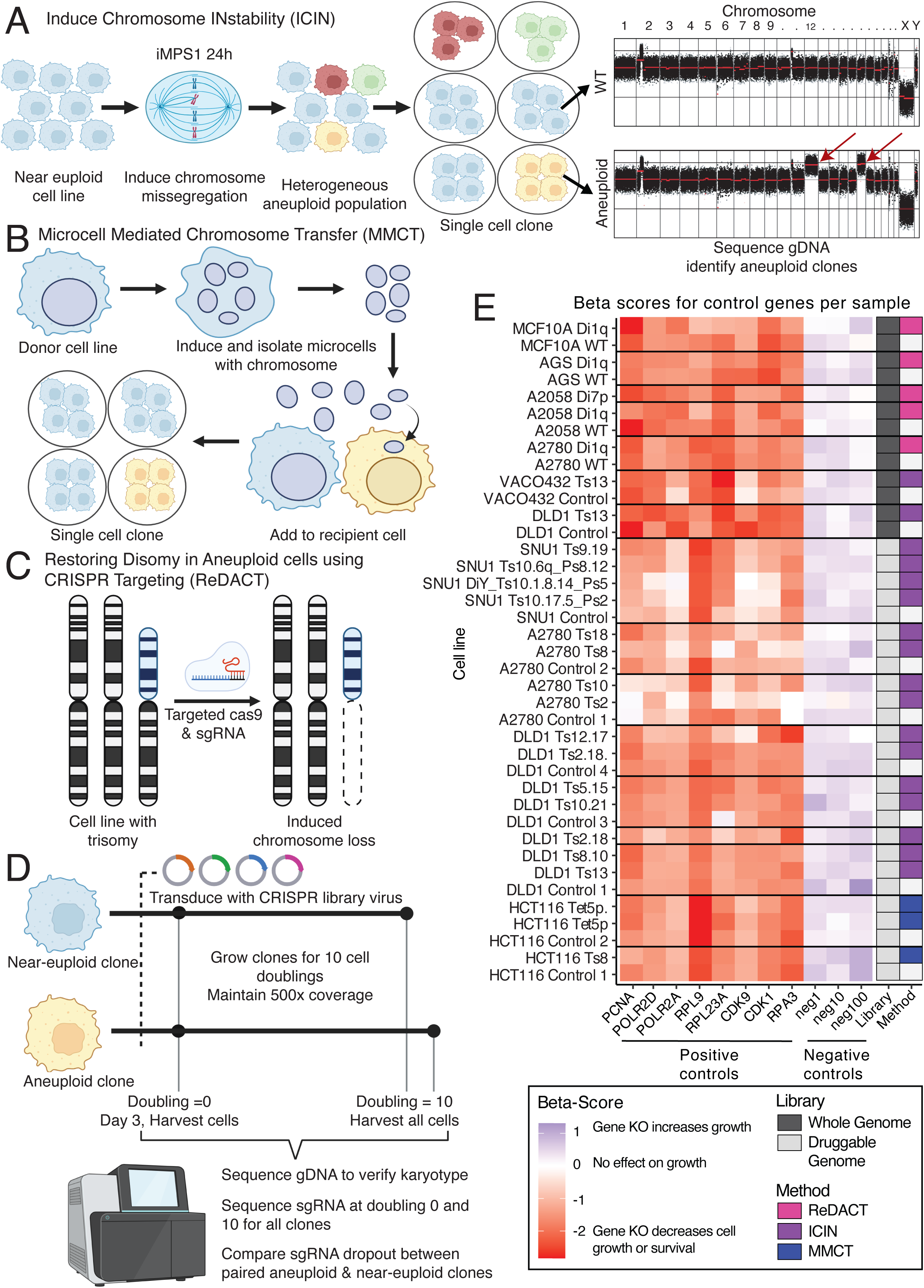
Paired aneuploid CRISPR screening strategy A–C) Schematics illustrating methods used to generate aneuploid cell lines: A) Induce Chromosome INstability (ICIN) followed by isolation of aneuploid clones; B) Microcell Mediated Chromosome Transfer (MMCT); C) Restoration of Disomy in Aneuploid cells using CRISPR Targeting (ReDACT). D) Overview of the paired CRISPR screening workflow, in which aneuploid clones are screened alongside their matched near-euploid control. E) Heatmap of quantile-normalized CRISPR beta scores for positive control genes and negative control sgRNAs across all screened cell lines. The CRISPR library (whole-genome library [WGL] or druggable genome [DG]) and the method used to generate each aneuploid clone are indicated. Black lines denote screening batches. SNU1 cells are near-tetraploid, whereas all other cell lines are near-diploid. Abbreviations: Ts, trisomy; Di, disomy; Tet, tetrasomy; Ps, pentasomy.

The second method used to generate aneuploid clones was Microcell Mediated Chromosome Transfer (MMCT). Briefly, microcells were induced in donor cells, and a microcell containing the chromosome of interest was isolated and injected into a recipient cell, as previously described^20,36,37^ (Figure 1B). These clones contained the gain of one chromosome or one chromosome arm.

Finally, the Restoring Disomy in Aneuploid Cells using CRISPR Targeting (ReDACT) methods were used to generate disomic subclones from cell lines harboring trisomies of 1q and/or 7p as previously described^16^ (Figure 1C). Newly generated disomy subclones were set as the near-euploid control relative to the parental trisomy cell line.

In total, 79 aneuploid subclones were generated or obtained from other laboratories (Supplementary Data 1). Paired subclones were generated across 11 cell lines, including five colon cancer cell lines (DLD1, HCT116, RKO, SW48, and VACO432), one ovarian cancer cell line (A2780), two stomach cancer cell lines (SNU1 and AGS), one melanoma cell line (A2058), and two untransformed epithelial cell lines (MCF10A and RPE1).

### Paired CRISPR screening to identify aneuploid-specific dependencies

Paired aneuploid and near-euploid control clones were screened using CRISPR libraries targeting either the whole genome (13 screens across 6 cell types) or the druggable genome (26 screens across 4 cell types). The Brunello Whole Genome Library (WGL) contains single guide RNAs (sgRNAs) targeting all 19,112 genes in the human genome, while the Druggable Genome (DG) library contains sgRNAs targeting 3,377 genes including the kinases, proteases, ubiquitin ligases, epigenetic modulators and transcription factors (Supplementary Data 1). The DG library was included to enrich for potentially therapeutic aneuploid dependencies. Since aneuploidy has been reported to reduce cell growth^11,12,33^, and differences in the number of cell divisions could influence gene dependency measurements, we monitored cell doubling during the screens and harvested each clone after 10 population doublings rather than after a fixed number of days (Figure 1D, Supplementary Data 1).

As aneuploidy can cause chromosome instability^12,36^, we sequenced nearly all cell lines at both the start (38/39 clones) and end (39/39 clones) of each screen to confirm that their karyotypes were maintained throughout the assay (Supplementary Data 2). 37 aneuploid clones retained their starting karyotypes while two aneuploid clones (HCT116 cells with a trisomy of chromosome 13 (Ts13) and DLD1 Ts4) exhibited evidence of chromosome loss by passage 10 and were removed from further analysis. HCT116 cells with tetrasomy 5p and trisomy 5q (Tet5p and Ts5q) lost a copy of chromosome 5q but maintained the 5p tetrasomy and so were relabeled Tet5p and were included in our analysis. DLD1 control 2 was removed due to low read coverage.

In total, 24 screens of 21 unique aneuploid clones and 15 near-euploid controls across eight human cell line types were screened and passed quality control. Two cell lines harbor loss-of-function mutations in (DLD1 and AGS) while six cell lines are TP53 wild-type. We screened three colorectal cancer (DLD1, HCT116 and VACO432), two stomach cancer (AGS and SNU1), one melanoma (A2058), one ovarian cancer (A2780), and one untransformed epithelial (MCF10A) cell line.

We screened six clones with chromosome arm level gains, eight clones with whole chromosome gains, six clones with double chromosome gains, three clones with both gains and losses, and one clone with only chromosome losses. The most commonly altered chromosomes, or chromosome arm, in our screens were 1q, 2, 5p, 8, and 10. These were each gained in three or more clones that we assayed. Across all screens, we screened subclones containing aneuploid copy numbers for 16 of the 22 autosomal chromosomes in human cells. All WGL pairs screened a cell line with one additional chromosome arm level trisomy relative to its isogenic disomy control.

Gene dropout beta scores were calculated for each cell line screened using the MAGeCK method^38^ (Supplementary Data 3, see methods), which enables robust identification of gene dependency from genome-scale CRISPR/Cas9 knockout screens. CRISPR screen results of samples that passed quality control and retained their aneuploid karyotype were merged and normalized (Supplementary Data 3). Normalized beta scores for pan-essential positive control genes and non-targeting negative control guide RNAs are shown for each cell line (Figure 1E).

To identify factors influencing gene dependency, CRISPR screens were clustered based on gene dropout profiles (Supplementary Figure 1A). Clone beta scores clustered primarily by CRISPR library, cell line type, and screening batch. Three aneuploid clones were screened twice. DLD1 Ts13 was screened with the WGL and DG library. WGL DLD1 Ts13 clustered with WGL DLD1 control more than DG DLD1 Ts13. The DLD1 cell line with trisomy 2 and 18 (Ts2.18) was screened twice in two different screening batches and clustered with the control clone of that batch more than with the other Ts2.18. Finally, HCT116 Tet5p was screened twice in the same screening batch and clustered together (Supplementary Figure 1A). Principal component analysis of beta scores, performed separately for the two CRISPR libraries showed clustering by cell line type rather than by aneuploidy status (Supplementary Figure 1B, 1C). Collectively, this demonstrates the significant effect libraries, cell type, and even screening batches can have on gene dropout scores. This underlies the importance of having batch-specific, paired cell line controls in our screens in order for us to identify biologically meaningful and clinically actionable dependencies.

### Identifying aneuploidy-specific dependency gene groups

To identify aneuploid-specific dependencies, we calculated the average difference in CRISPR beta score between aneuploid clones and their paired near-euploid controls in the WGL screen (Figure 2A). We also calculated the average difference in beta score for the merged DG and WGL datasets, restricting the analysis to genes present in both libraries (Figure 2B; Supplementary Data 3).

**Figure 2:**
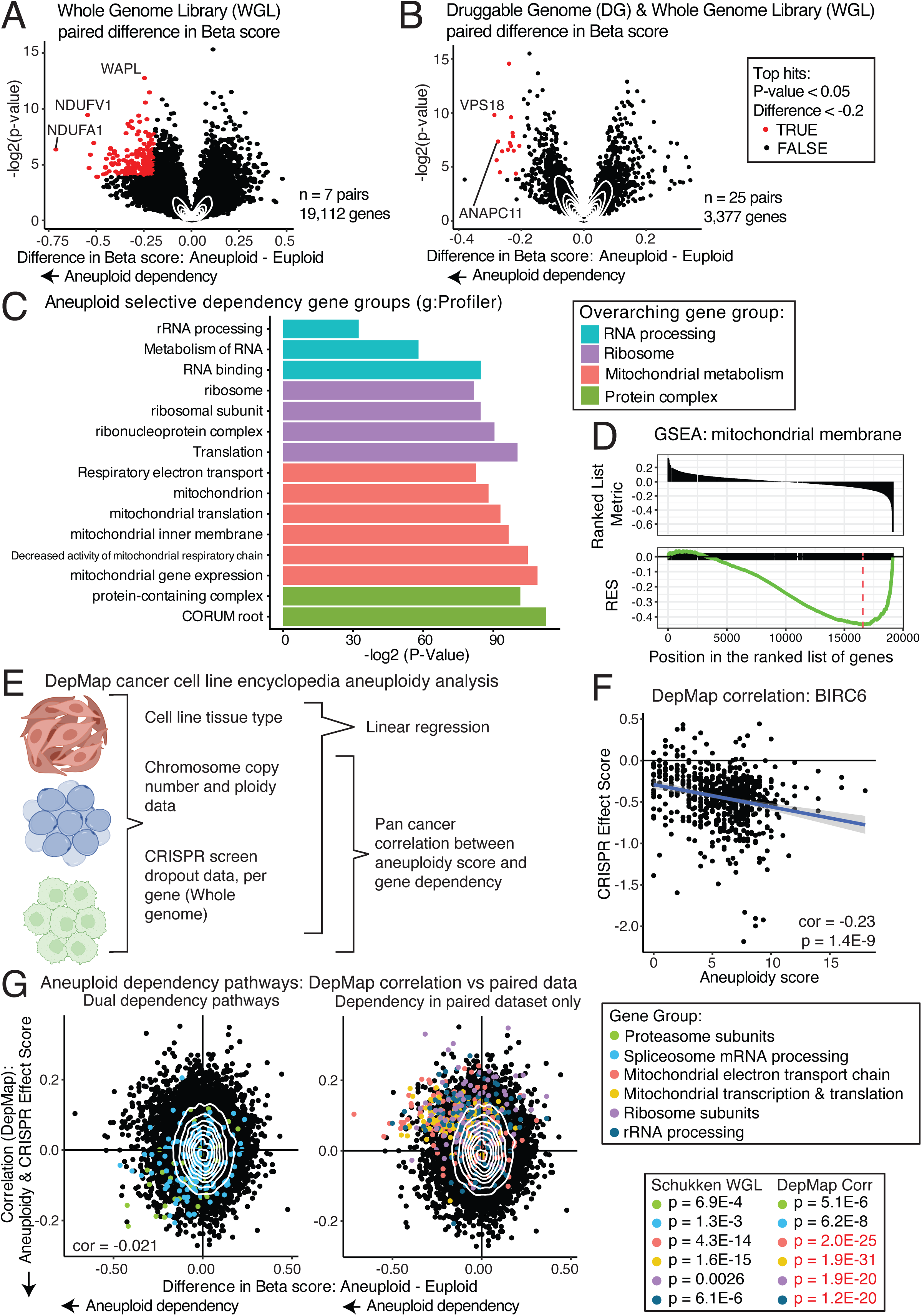
Identification of aneuploidy dependency gene groups A–B) Normalized paired CRISPR screen results. Each point represents a gene. The x-axis shows the average paired difference in CRISPR beta score between aneuploid clones and their matched near-euploid controls; the y-axis shows the –log₂(p-value) from a two-sided paired t-test. Negative beta score differences indicate increased dependency in aneuploid cells. A) All genes screened in the WGL, and B) screen samples from both WGL and DG libraries, with only the genes present in the DG library. Genes with p < 0.05 and difference in beta score < –0.2 are highlighted in red. C) Gene sets significantly enriched among aneuploid dependencies in the WGL paired screen, identified using g:Profiler and colored by overarching gene group. D) Gene set enrichment analysis (GSEA) for Mitochondrial Membrane gene set. E) Schematic of DepMap Cancer Cell Line Encyclopedia (CCLE) aneuploid dependency analysis. F) Example correlation between aneuploidy score (number of aneuploid chromosome arms divided by ploidy) and CRISPR Effect Score for BIRC6 across DepMap cell lines. Pearson correlation coefficient and p-value are shown. G) Comparison of aneuploid dependency gene groups in DepMap correlation analysis and WGL paired screens. Left: gene groups showing concordant aneuploid dependency in both datasets. Right: gene groups identified only in paired screens. Two-sided t-test p-values for each gene group per dataset. Black text denotes aneuploid dependency; red text denotes aneuploid resistance.

To identify cellular gene groups upon which aneuploid cells are preferentially dependent, we analyzed aneuploid-specific dependencies from the WGL screen using both g:Profiler and Gene Set Enrichment Analysis (GSEA) (Figure 2C, 2D; Supplementary Figure 2A, 2B; Supplementary Data 4). The WGL screens only analyze cell lines with single chromosome arm gains relative to isogenic near-diploid controls, so aneuploid dependencies found in this analysis may be specific to chromosome gains. This analysis revealed significant enrichment of genes involved in RNA processing, ribosomal genes, protein complexes, and mitochondrial metabolism including both components of the electron transport chain and mitochondrial transcription and translation.

Next, we compared results from our paired screens with cancer cell line gene dependency data from the Cancer Dependency Map (DepMap) project^39^. DepMap contains CRISPR dropout data^39–43^, transcriptomic data^44^, proteomic data^45^ and clinical metadata pertaining to hundreds of human cancer cell lines across multiple tumor types. Analyzing DepMap data allowed us to compare our data to broad pan-cancer trends. For each gene, we calculated the Pearson correlation coefficient between gene dependency (CRISPR Effect Score) and cell line aneuploidy score, which was calculated as the number of aneuploid chromosome arms^30^ normalized by ploidy (Figure 2E; Supplementary Data 5). For example, there is a significant negative correlation (cor=-0.23, p=1.4E-9) between aneuploidy score and BIRC6 CRISPR Effect Score, showing that highly aneuploid cell lines tend to be more dependent upon the anti-apoptotic gene BIRC6 (Figure 2F).

Several aneuploid-specific dependency gene sets were shared between the DepMap correlation analysis and the paired beta score differences observed in our screens (Supplementary Data 6). Shared gene sets included proteasome subunits and mRNA processing via the spliceosome (Figure 2G, left). In contrast, genes involved in mitochondrial electron transport chain (including both the subunits and genes involved with assembly), mitochondrial transcription and translation genes (including mitochondrial ribosome genes and mitochondrial transcription factors), ribosomal subunits (cytosolic), and rRNA processing genes were identified as aneuploid dependencies in our paired screens but not in the DepMap correlation analysis (Figure 2G, right).

To control for the effect of cell lineage in DepMap aneuploidy-specific dependencies, we performed multivariate linear regression between CRISPR effect scores and aneuploidy score while setting cancer lineage as an independent variable (Supplementary Figure 3A; Supplementary Data 5). The aneuploidy score coefficient in the linear model showed that proteasome and spliceosome genes continued to exhibit a greater dependency in aneuploid cells, consistent with our results (Supplementary Figure 3B). The linear model showed mitochondrial electron transport chain, mitochondrial transcription and translation, ribosomal subunits and rRNA processing genes had significantly greater dependency in near-euploid cells (Supplementary Figure 3C). The linear regression model and the aneuploidy score correlation both show similar trends in DepMap aneuploidy-specific dependencies.

DepMap primarily profiles established cell lines that may have adapted to chronic aneuploid stresses, whereas our paired screens analyze low passage number aneuploid or low passage number near diploid subclones. We considered whether the discrepancies between the DepMap and paired screen aneuploid dependencies might be due to adaptation over time. Bökenkamp et al. (2025)^25^ recently investigated protein and transcriptional changes over time in newly induced aneuploid clones. We reanalyzed their protein data and found that newly generated aneuploid clones (passage 0) have significantly lower ribosome and rRNA processing protein abundance relative to isogenically matched WT cells (Supplementary Figure 3D). Although ribosomal proteins do show significant upregulation in aged clones (passage 50) relative to passage 0 (Supplementary Figure 3E), this upregulation is not a full rescue, as late aneuploid clones still show reduced ribosome proteins relative to wild-type (Supplementary Figure 3F). This shows an adaptation to aneuploid induced ribosomal stress over time and may contribute to the reduced aneuploid-specific dependency on ribosomal and rRNA processing genes in DepMap compared to our lower passage aneuploid clones.

Similar to ribosomes, electron transport chain associated proteins and mitochondrial transcription and translation proteins show a significantly reduced abundance in low passage aneuploid clones relative to wild-type (Supplementary Figure 3G). Unlike ribosomal proteins, mitochondrial metabolism proteins show further decreased abundance at passage 50 in aneuploid cell lines (Supplementary Figure 3H, 3I). While this finding does not rule out alternative metabolic adaptations to aneuploidy, it shows a consistent downregulation of mitochondrial metabolic proteins in stable aneuploid clones.

### Identifying UBE2H and FOSL1 as top aneuploid dependency genes

To identify high-confidence aneuploid-specific dependencies, we selected genes that showed significant aneuploid dependency both in our paired screen (difference< -0.2, p<0.05) and in DepMap correlation analysis (correlation score< -0.1, p<0.005). Analysis of the WGL dataset (7 pairs; 19,112 genes) using these thresholds identified 15 top aneuploid-specific dependencies (Figure 3A). This list includes an ESCRT-III subunit (CHMP6), a signalosome component that regulates E3 ubiquitin ligases (COPS6), a translation associated mRNA gene (LARP4), a mitochondrial biogenesis regulator (PPRC1), two proteasome subunits (PSMA3, PSMB7), and an AP-1 transcription factor subunit (FOSL1).

**Figure 3:**
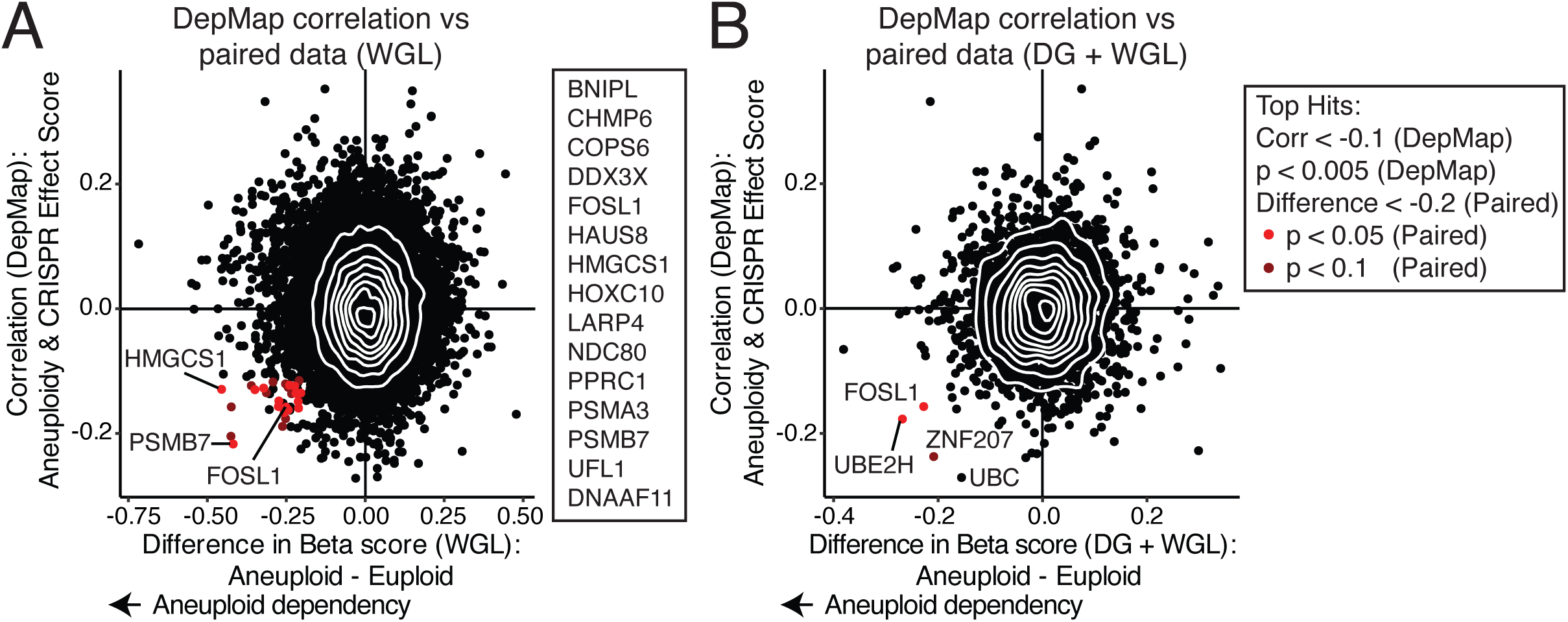
Identification of top aneuploid-specific dependencies A–B) Paired CRISPR beta score differences from A) WGL-only screens or B) merged DG and WGL screens, plotted against DepMap Pearson correlation scores between aneuploidy burden and CRISPR Effect Score. Aneuploidy burden is defined as the number of aneuploid chromosome arms divided by ploidy. Genes meeting the following criteria are highlighted: DepMap correlation < –0.1, DepMap p < 0.005, paired beta score difference < –0.2, and paired p < 0.05 (red). Genes meeting all criteria except paired p-value < 0.1 are shown in dark red.

Next, paired samples from the DG and WGL screens (25 pairs) were combined; analysis was restricted to the genes present in the DG library (3,377 genes). Ubiquitin C (UBC) showed an aneuploid trend but did not meet our significance cutoffs. Two genes did meet our significance criteria for aneuploidy-specific dependency: UBE2H and FOSL1 (Figure 3B).

To identify dependencies associated with total aneuploidy burden, we calculated an aneuploidy dependency score based on the number of aneuploid chromosomes present in each cell line, normalized to their basal ploidy (Supplementary Figure 4A, Supplementary Data 7). Comparison of our total aneuploidy burden correlations with DepMap aneuploidy correlations identified 76 shared dependencies (Supplementary Figure 4B, Supplementary Data 8). UBE2H showed an aneuploid trend (correlation=-0.28, p=0.085). Top hits included PPP1R12A, the aneuploidy adaptation-associated transcription factor FOXM1^25^, and WDR26, a member of the CTLH E3 ubiquitin ligase complex that exclusively uses UBE2H as its E2 cognate^46,47^, among other hits. We repeated this analysis, this time including data from the DG library and we found that nine genes had dual significance in both our aneuploidy burden correlation and DepMap analyses (Supplementary Figure 4C), including ubiquitin (UBC).

### Independent paired screens show similar aneuploidy-specific dependencies

To identify true and consistent aneuploid dependencies shared across laboratories and screen designs, we compared our data with two recently published paired CRISPR screens of non-cancer aneuploid and near-euploid cell lines. Magesh et al. (2025)^27^ screened four aneuploid clones derived from a human mammary epithelial cell line (hMEC), while Zerbib et al. (2024)^33^ screened three aneuploid clones of the RPE1 cell line (Figure 4A). Both screens focus primarily on chromosome gains relative to paired diploid controls.

**Figure 4:**
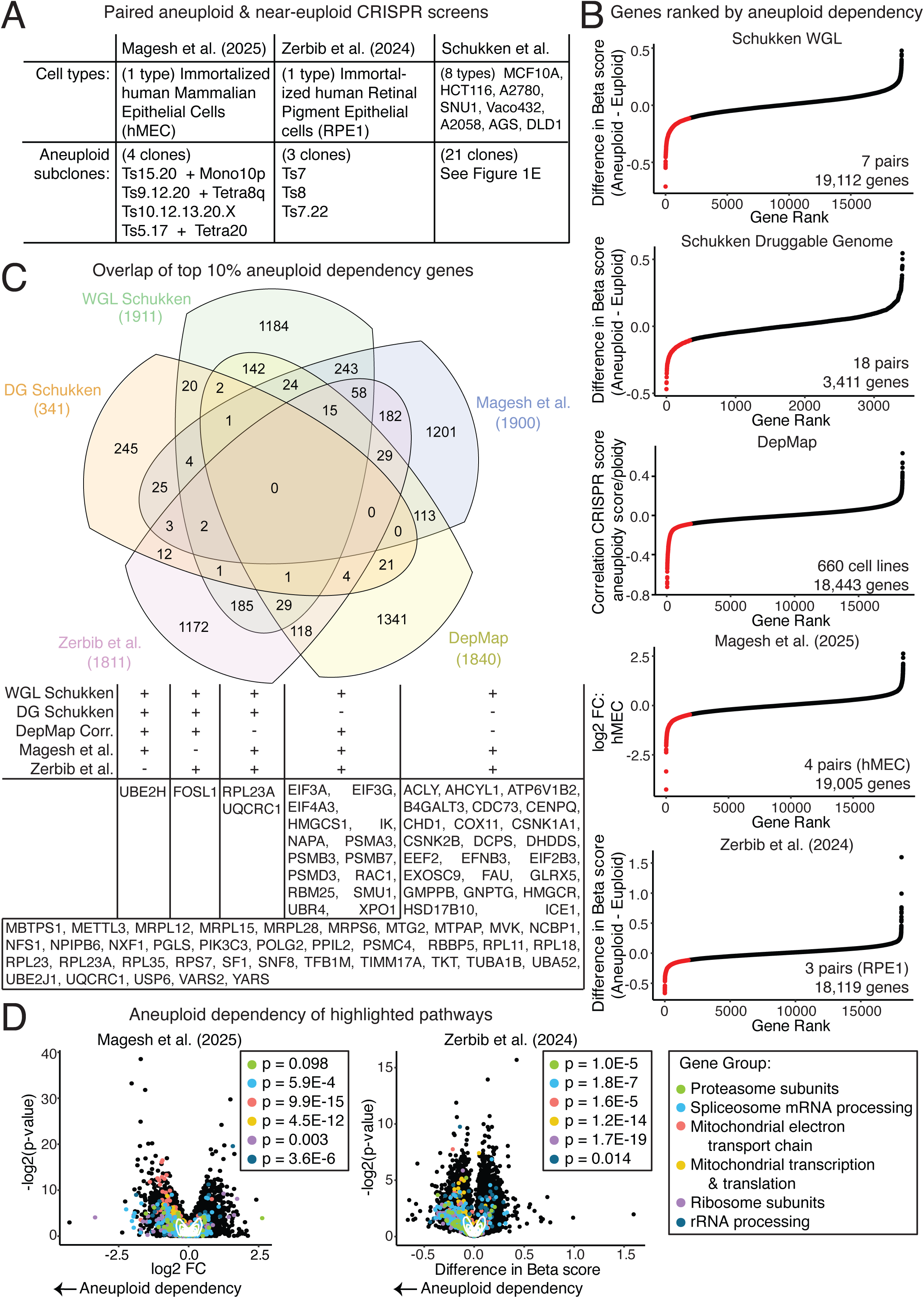
Comparison of aneuploid dependencies to other paired aneuploid CRISPR screens A) Summary table of cell line types and numbers of aneuploid subclones analyzed in each published paired screen. B) Gene rankings by aneuploid dependency measurement for each dataset. C) Venn Diagram of overlapping genes within the top 10% of aneuploid dependency rankings across datasets. Table of gene lists within top 10% of indicated datasets. D) Comparison of paired aneuploid dependency scores and –log₂(p-values) per gene from Zerbib et al. and Magesh et al. Highlighted gene groups are indicated. Two-sided t-test p-values are shown; black text denotes aneuploid dependency and red text denotes aneuploid resistance.

To compare these datasets, we ranked all genes by their respective aneuploid dependency scores and identified the top 10% of genes in each dataset with the largest aneuploid dependency (Figure 4B; Supplementary Data 8). Several genes overlapped as aneuploid-specific vulnerabilities in four of the five datasets (our WGL and DG screens, Magesh et al., Zerbib et al., and DepMap correlation). FOSL1 was ranked as a top aneuploid dependency in all except Magesh et al., while UBE2H was a top aneuploid dependency in all except Zerbib et al.

When we excluded the DG library screen and only considered the larger whole genome libraries, we identified 15 genes shared across the remaining datasets (Figure 4C). These genes include proteasome subunits (PSMA3, PSMB3, PSMB7, PSMD3), translation initiation factors (EIF3A, EIF3G, EIF4A3), an E3 ubiquitin ligase (UBR4), and spliceosome components (IK, SMU1), among others. When we identified aneuploid dependency genes shared between the three whole genome paired screens (our WGL Schukken, Magesh et al. and Zerbib et al.) we identified 58 additional overlapping genes that were not top hits in DepMap (Figure 4C). These included ribosomal subunits, rRNA processing genes, mitochondrial translation genes, and electron transport chain associated genes, among others.

Consistent with the findings in our own paired aneuploidy screen, mitochondrial electron transport chain genes, mitochondrial transcription and translation genes, ribosomal genes, and rRNA processing genes showed greater dependency in aneuploid clones relative to their wild-type controls in both Magesh et al. and Zerbib et al. (Figure 4D). All but one of these gene groups showed statistically significant aneuploid dependency relative to other genes in the dataset, and the proteasome subunits showed a trend toward aneuploid dependency in Magesh et al. (p-value=0.098).

### Chromosome specific & cell line specific analyses highlight heterogeneity in aneuploid dependencies

We investigated chromosome-specific aneuploid dependencies. For the WGL screens, we analyzed clones with trisomy 1q, 7p, and 13 independently. All three trisomy groups had significantly increased dependency upon the mitochondrial metabolism gene group (Supplementary Figure 5A-C). However, the wildtype Ts1q and Ts7p cell lines showed little to no dependency on spliceosome genes, proteasome subunits, or ribosome subunits relative to their ReDACT generated disomy subclones. The Ts13 clones, both generated via ICIN, did show significant dependencies on these gene groups. These differences may indicate chromosome specific dependencies or adaptation to aneuploidy over time.

Next, we split DepMap cell lines into those with gains for a specific chromosome arm and those without and calculated the average difference in CRISPR Effect Score between these two groups. Chromosome arm specific hits were identified per chromosome arm by comparing the paired WGL and DepMap arm level dependencies (Figure 5A, Supplementary Data 3). Trisomy 1q increases dependency on the nuclear RNA export regulator NXT1. Trisomy 7p made A2058 cells more dependent on GRB2, an adaptor protein that plays a role in multiple signaling cascades, including linking EGFR to its downstream pathways^48^; the gene for EGFR is located on chromosome 7p. Finally, the gain of chromosome 13 increases dependency upon CTNNB1 and IRS2, two key components of the Wnt pathway, as well as upon TCF7L2, a key transcriptional effector of the Wnt pathway. The gene for IRS2 is located on chromosome 13.

**Figure 5:**
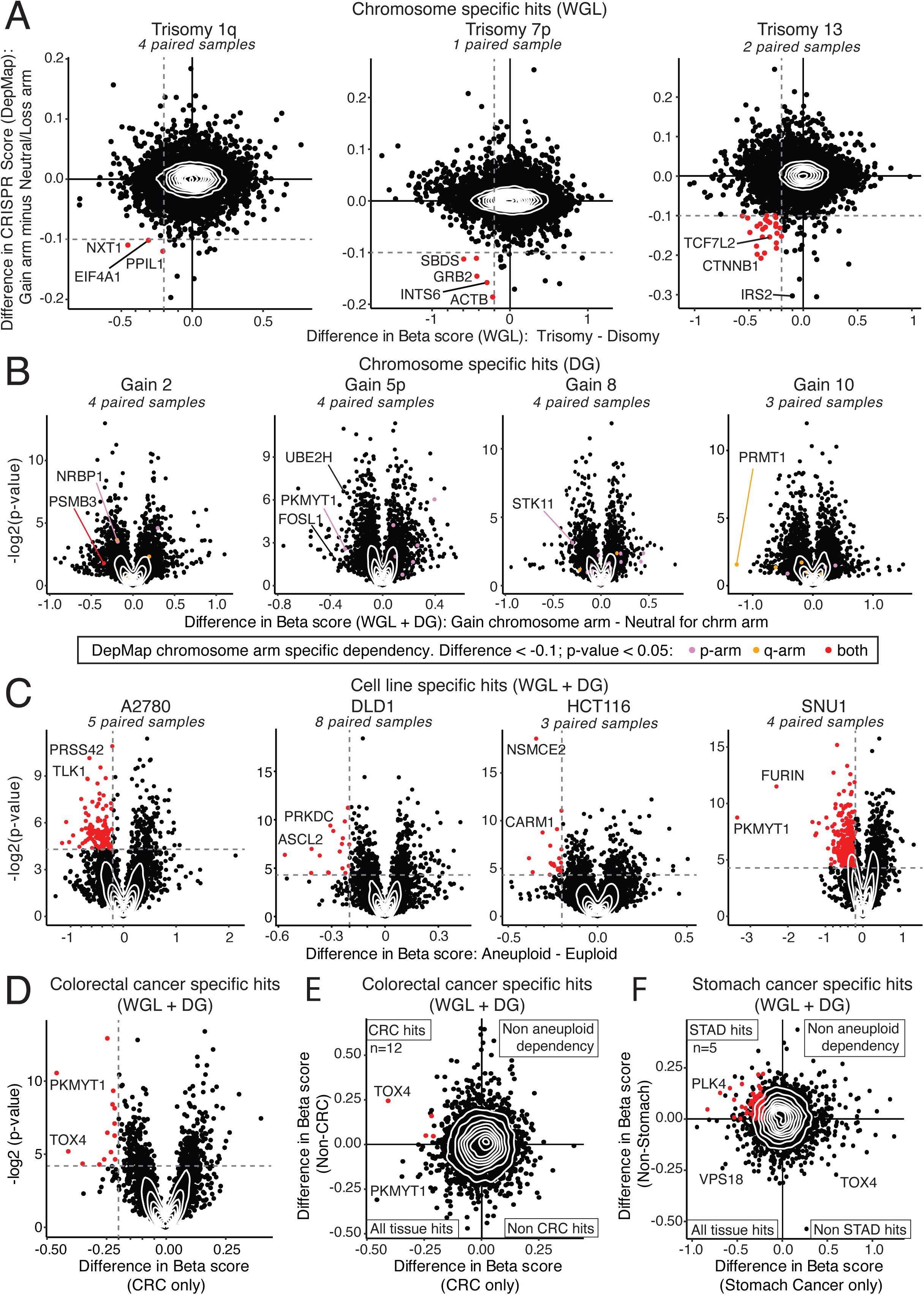
Chromosome and tissue specific gene dependency analysis highlights aneuploid heterogeneity A) WGL paired screens grouped by shared trisomic chromosome arm. The x-axis shows the average paired beta score difference between trisomy and disomy pairs for the indicated chromosome arms; the y-axis shows the average difference in DepMap CRISPR Effect Scores between cell lines with gains versus neutral or loss copy number for the same arm. Cutoffs were set at –0.2 for WGL paired data and –0.1 for DepMap data. B) DG paired screens grouped by shared chromosome gains. The x-axis shows the paired beta score difference per gained chromosome arm; the y-axis shows -log₂(p-value) from paired t-tests. For each chromosome arm, average differences in CRISPR Effect Score were calculated for DepMap data between cell lines with a gain for that arm versus a neutral or loss copy number. Significant arm-level dependencies in DepMap are highlighted (DepMap difference < –0.1 and paired p < 0.05). C) Paired DG and WGL screens grouped by cell type. The x-axis shows the paired beta score difference for each cell line (A2780, DLD1, HCT116 and SNU1); the y-axis shows –log₂(p-value). Significance cutoffs: difference < –0.2 and p < 0.05. D) Average paired beta score difference across colorectal cancer cell lines (HCT116, DLD1, VACO432). Y axis displays the -log₂(p-value) of paired t-test. Significance cutoff at difference < -0.2 and p-values < 0.05. E) Comparison of average paired beta score differences in colorectal cancer cell lines versus all non-colorectal cell lines. Significance cutoff at difference < -0.2 in colorectal and > 0 in non-colorectal cell lines. F) Comparison of average paired beta score differences in stomach cancer cell lines (SNU1, AGS) versus all non-stomach cell lines. Significance cutoff at difference < -0.2 in stomach and > 0 in non-stomach cell lines.

We further investigated chromosome specific hits by expanding our analysis to include the cell lines screened with the DG library. Four additional chromosomes (arms) had at least three different paired trisomy and disomy clones: 5p, 2, 8, and 10. Our paired dependencies were compared to DepMap arm level dependencies to identify chromosome specific hits (Figure 5B). Trisomy 5p, 8, and 10 increase dependence upon PKMYT1, STK11, and PRMT1, respectively. These genes were also found to be aneuploid dependencies in our total paired analysis. Gaining chromosome 2 increases dependency on NRBP1, a proposed WNK pathway regulator^49^, whose gene is located on chromosome arm 2p.

Since CRISPR beta scores cluster by cell type (Supplementary Figure 1A), we considered if each cell type had unique aneuploid-specific dependencies, akin to tissue-specific dependencies in tumors. Grouping paired samples by cell line type, we analyzed HCT116, DLD1, A2780, and SNU1 cells independently, as we had a minimum of three aneuploid clones for these cell lines (Figure 5C). We found that no single gene was a significant (difference< -0.1, p<0.05) dependency in all four cell types. This underscores the substantial effect cell line variance has on aneuploid dependency. However, several genes had consistent aneuploid trends in all cell types. Six genes were within the top 10% of aneuploid dependencies in all four cell lines, and 23 genes were within the top 20% of all four cell lines (Supplementary Data 8). Randomly sampling ten percent of the DG four times would have led to an estimated 1.1 overlapping genes, and randomly sampling twenty percent would have only led to an estimated 5.4 genes. Shared dependencies include proteasome subunit PSMD14, heat shock transcription factor HSF1, cell cycle regulators PLK3 and PKMYT1, AP-1 transcription factor FOSL1, and ubiquitin E2 ligase UBE2H.

Next, we investigated tissue-specific aneuploid dependencies. Since the largest tumor type represented in our dataset was colorectal cancer (HCT116, DLD1, and VACO432), we combined all paired cell line clones and identified the top colorectal cancer aneuploid dependencies (Figure 5D). We then grouped all non-colorectal pairs and identified top aneuploid dependencies. Comparing colorectal and non-colorectal aneuploid dependencies, we identified TOX4 as a potential colorectal-specific aneuploid dependency (Figure 5E). A similar analysis of stomach cancer cell lines (AGS and SNU1) identified PLK4 as a potential stomach cancer-specific aneuploid dependency (Figure 5F).

Based on the consistency of aneuploid-specific dependency across multiple analytical approaches, we selected UBE2H for experimental validation. Unlike FOSL1, UBE2H was a top hit in both the paired analysis and the total aneuploidy burden analysis. Additionally, UBE2H’s protein structure was predicted to be druggable, while FOSL1 was not^50^. This makes UBE2H a potential therapeutic target for highly aneuploid cancers. For these reasons, we focused primarily on UBE2H for experimental validation.

### Validating UBE2H as an aneuploid dependency

To validate UBE2H as an aneuploid-specific dependency, we performed competition assays in a SNU1 near-tetraploid control clone and three aneuploid SNU1 subclones. Our CRISPR screens primarily assayed aneuploid clones with chromosome gains. To validate UBE2H as a broader aneuploid dependency, we selected aneuploid clones with primarily chromosome losses. Two of the SNU1 aneuploid clones harbor both chromosome gains and losses, and one clone contains exclusively losses. For each cell line, we transduced cells with a GFP, Cas9, sgRNA vector, where the sgRNA targeted either a control gene or our gene of interest. We then measured the fold change in the proportion of GFP-positive cells over time for each transduction (Supplementary Figure 6A). In the SNU1 control cell line, UBE2H sgRNAs showed an 18% dropout relative to pan-essential genes, whereas relative sgRNA dropout was markedly increased in aneuploid clones: 49% in clone 114 (Ts9.19), 79% in clone 12 (Ts10.5.17 and Pentasomy 2 (Ps2)), and 92% in clone 24 (Ts1.8.10.14.Y Ps5) (Figure 6A-D). When analyzed collectively, aneuploid clones exhibited significantly greater dependency on UBE2H than control cells (p=8.6E-4).

**Figure 6:**
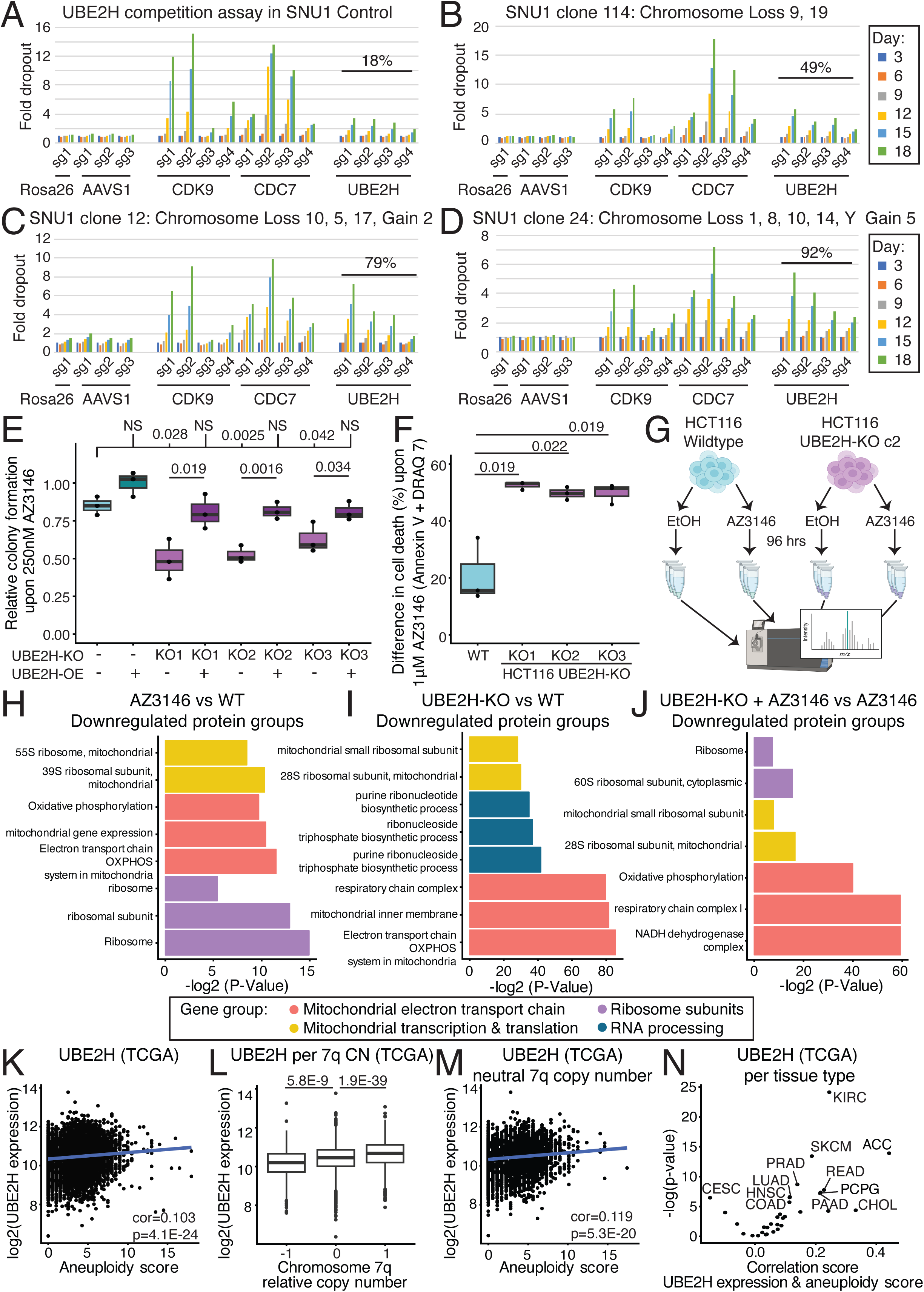
Validation and mechanistic analysis of UBE2H A-D) Competition assay results showing fold dropout of sgRNAs targeting negative controls (ROSA26, AAVS1), positive controls (CDK9, CDC7), and UBE2H in A) SNU1 near-tetraploid control cells, B) SNU1 aneuploid subclone 114 (trisomy 9, 19), C) SNU1 aneuploid subclone 12 (trisomy 5, 10,17; pentasomy 2), D) SNU1 aneuploid subclone 24 (trisomy 1,8,10,14; DiY; pentasomy 5). Percent dropout is normalized relative to positive and negative controls. E) Colony formation following AZ3146 treatment (250 nM) relative to ethanol vehicle control in HCT116 wild-type cells, three UBE2H knockout clones, and UBE2H cDNA rescue clones. P-values from two-sided t-test relative to wild-type are shown. One sided t-test for overexpression clones rescue relative to knockout. F) Change in percentage of Annexin V and/or DRAQ7-positive cells following 96 h treatment with AZ3146 (1 µM), relative to ethanol-treated controls in HCT116 wild-type and UBE2H-KO clones. Two-sided t-tests. G) Diagram of sample preparation for Mass Spectrometry analysis. H-J) g:Profiler gene group enrichment analysis of mass spectrometry data. Protein abundance downregulated in HCT116 in H) AZ3146 treated versus wild-type, I) UBE2H-KO versus wild-type and J) AZ3146 treated UBE2H-KO versus AZ3146 treated wild-type cells. K) UBE2H RNA expression correlated to aneuploidy score in TCGA human tumor data. Pearson correlation and corresponding p-value. L) UBE2H RNA expression stratified by chromosome arm 7q copy number status; with 7q arm loss (-1), neutral (0) and gain (1) in TCGA data. Two-sided t-test p-values are shown. M) UBE2H RNA expression correlated to aneuploidy score for samples with a neutral copy of chromosome arm 7q in TCGA data. N) UBE2H RNA expression and aneuploidy score Pearson correlation coefficient and -log₂(p-value) per tumor types (TCGA). All significant correlations labeled with tumor abbreviation.

To further verify UBE2H aneuploid dependency, we generated UBE2H knockout clones in the near-diploid HCT116 and Cal51 cell lines by transfecting cells with two independent UBE2H-targeting sgRNAs followed by single-cell cloning^51^. Loss of UBE2H protein was confirmed by Western blotting (Supplementary Figure 6B and 6C). To further assess the effects of losing UBE2H on the viability of aneuploid cells, we treated UBE2H-wildtype and UBE2H-knockout cells with the MPS1 inhibitor AZ3146 and then quantified colony formation. In both Cal51 and HCT116, UBE2H-knockout clones showed significantly reduced colony formation relative to control cells upon AZ3146 treatment (Figure 6E; Supplementary Figure 6D). To exclude off-target effects of UBE2H sgRNAs, we performed rescue experiments using an UBE2H cDNA construct in HCT116 clones (Supplementary Figure 6B). Colony formation assays demonstrated that UBE2H cDNA overexpression fully rescued colony formation in all three knockout clones relative to wild-type (Figure 6E).

To determine whether the reduced colony formation in aneuploid UBE2H-knockout cells reflected increased cell death, rather than slower growth, HCT116 and Cal51 cells were treated with AZ3146 for 96 hours and then stained with the apoptosis marker Annexin V and the viability dye DRAQ7. All HCT116 UBE2H knockout clones exhibited significantly increased cell death upon AZ3146 treatment compared to HCT116 wild-type treated with AZ3146 (Figure 6F), and one of the two Cal51 UBE2H-KO also showed increased cell death (Supplementary Figure 6E).

To investigate the molecular mechanism underlying UBE2H dependency, we performed mass spectrometry-based proteomic analysis on HCT116 wild-type cells and UBE2H knockout clone 2 treated with a vehicle control or with AZ3146 to induce aneuploidy (Figure 6G; Supplementary Data 9). As expected, UBE2H knockout samples showed significantly reduced UBE2H protein abundance relative to wild-type cells (Supplementary Figure 6F).

We used gene set profiling to identify significantly down and upregulated proteins (Figure 6H-J, Supplementary Figure 6G-I). Ribosomal proteins were the most significantly downregulated proteins in AZ3146 treated HCT116 cells (Figure 6H). AZ3146 treatment also led to a small but significant downregulation of mitochondrial metabolism associated proteins (Figure 6H, Supplementary Figure 6J). UBE2H knockout resulted in a greater significant reduction in mitochondrial metabolism proteins than AZ3146 treatment (Figure 6I, Supplementary Figure 6K). AZ3146 treatment of UBE2H-KO cells led to a significantly greater reduction in mitochondrial metabolism proteins compared to AZ3146 treatment of wild-type cells (Figure 6J, Supplementary Figure 6L). In summary, these results indicate that AZ3146 induces a modest reduction in mitochondrial metabolism proteins and UBE2H loss amplifies these effects.

UBE2H localizes to mitochondria and is the only E2 ubiquitin-conjugating enzyme reported to localize to this organelle in the Human Protein Atlas^52–54^. RNA expression clustering in human tumors places UBE2H within cluster 76, classified as non–tissue-specific metabolism with high confidence (0.99)^53,55^. This cluster was enriched for mitochondrial matrix and oxidoreductive associated genes, including 31 genes involved in the mitochondrial electron transport chain or mitochondrial transcription and translation.

We hypothesized that UBE2H expression promotes cell survival in the presence of aneuploidy. To test this hypothesis, we analyzed previously published gene expression data from near-diploid and aneuploid HCT116 cell lines^20,21^, and we found that UBE2H was in the top 10% of RNAs and proteins upregulated by aneuploidy (Supplementary Figure 7A-B). Next, we analyzed isogenic strains of aneuploid budding yeast^56^, and found that the UBE2H homolog UBC8 was similarly upregulated in disomic aneuploid yeast strains (Supplementary Figure 7C-D), indicating that its role in promoting the growth of aneuploid cells may be evolutionarily conserved.

To investigate the role of UBE2H in aneuploid human cancers, we analyzed DepMap cancer cell line and The Cancer Genome Atlas (TCGA) human tumor data^39–45,57–60^. We found a positive correlation between UBE2H RNA expression and aneuploidy score in human tumors (TCGA: Pearson correlation score=0.103, p=4.1E-24; Figure 6K) and in cancer cell lines (DepMap: corr=0.135, p=6.7E-4; Supplementary Figure 7E). Next, we investigated whether tumor cells might select for a gain of UBE2H, which is located on chromosome arm 7q. Chromosome arm 7q was the fifth most frequently gained chromosome arm in TCGA, with 24% of tumors gaining this chromosome arm^58–60^. UBE2H was not reported to undergo significant dosage compensation upon chromosome gain, but does show compensation upon chromosome loss^17^. Consistent with this, human tumors with a gain of chromosome 7q also show increased UBE2H expression (p=1.9E-39; Figure 6L). Similarly, gaining a copy of chromosome 7q was associated with a significant increase in UBE2H RNA expression (p=1.7E-12) and protein abundance (p=8.0E-7) in cancer cell lines (Supplementary Figure 7F-G), while losing 7q did not result in a significant decrease in UBE2H protein. For tumors without a gain of chromosome 7q, UBE2H expression still positively correlates with aneuploidy scores in human tumors (corr=0.119; p=5.3E-20; Figure 6M), and in cancer cell lines (corr=0.123, p=0.017; Supplementary Figure 7H). Finally, to account for tumor type differences, we correlated aneuploid scores and UBE2H expression per tumor type and found that most tumor types had a positive correlation. Eleven tumor types had a significant positive correlation between UBE2H expression and aneuploidy score, including renal clear cell carcinoma, adrenocortical carcinoma, pancreatic adenocarcinoma, and skin cutaneous melanoma (Figure 6N; Supplementary Figure 7I-L).

## Discussion

Aneuploidy is a hallmark of cancer, yet the cellular dependencies created by large-scale copy number imbalance are not fully understood. Here, by systematically profiling genetic dependencies across a diverse panel of isogenic aneuploid and near-euploid human cancer cell lines, we identify a core set of cellular pathways that are consistently required for aneuploid cell fitness. These dependencies span multiple biological processes—including the proteasome, the spliceosome, ribosomes, rRNA processing, mitochondrial transcription and translation, and the mitochondrial electron transport chain—and are conserved across distinct cancer types, karyotypic configurations, and methods of inducing aneuploidy. Together, our results suggest that aneuploidy imposes a conserved set of physiological stresses that create exploitable vulnerabilities in cancer cells.

A major strength of this study is the breadth of aneuploid contexts examined. Using ICIN, we generated 63 novel aneuploid clones across 6 cell types. We then performed 39 CRISPR screens with two libraries, across eight cell types, and on aneuploid subclones generated with three independent approaches (ICIN, MMCT and ReDACT). We screened both newly generated aneuploid subclones and adapted aneuploid cell lines relative to their newly generated more-euploid controls. These broad screens demonstrate that many aneuploid dependencies are not artifacts of a specific cell line, karyotype, or aneuploid induction technique. Instead, they reflect fundamental challenges faced by cells harboring chromosome-scale copy number alterations.

In particular, the consistent identification of proteasome and spliceosome-associated genes aligns with prior work demonstrating that aneuploidy increases the burden of aberrant transcripts and misfolded or excess proteins, thereby heightening reliance on protein quality control and RNA processing machinery^22,24,61,62^. Consistent with these results, proteasome and spliceosome components are also pan-cancer aneuploid dependencies in the DepMap dataset.

Similarly, dependencies on ribosomal genes and rRNA processing genes in our isogenic screens reinforce previous studies showing that aneuploid cells experience ribosome imbalance and translational stress as a direct consequence of gene dosage perturbations^17,20,21,25,26^. However, DepMap shows the opposite effect, with highly aneuploid cell lines being less dependent on ribosomal genes. Ribosomal dependency may represent a temporary aneuploid dependency of newly generated aneuploid cells. Consistent with this, the gain of chromosome 13 induced a strong ribosomal dependency in our subclones, while adapted trisomic cells (Ts1q or Ts7p) with disomy subclones showed little to no change in ribosomal dependence. Additionally, Bökenkamp et al.^25^ show an initial strong downregulation of ribosomal abundance that is partially rescued at a later passage of the same aneuploid cells.

Notably, our analyses also reveal mitochondrial processes as a central and recurrent vulnerability of aneuploid cells. Genes involved in mitochondrial transcription and translation, as well as components of the mitochondrial electron transport chain, emerged as top aneuploid-specific dependencies across multiple datasets. These findings extend previous observations that aneuploidy disrupts cellular metabolism and increases energetic demand^26,27^ and suggest that maintaining mitochondrial function is particularly critical for the survival of cells with imbalanced genomes. The convergence of mitochondrial dependencies across independent isogenic paired screens highlights mitochondrial metabolism as a core axis of aneuploid cell fitness. Unlike ribosomal protein abundance, Bökenkamp et al. show both newly induced aneuploid clones and late passage adapted aneuploid clones have reduced mitochondrial protein abundance and fail to show protein upregulation over time. We also found that AZ3146-treated cells downregulate mitochondrial protein abundance. Consistent with this, both cell lines with the gain of chromosome 13 and adapted trisomic cell lines (Ts1q and Ts7p) showed significant aneuploid-specific dependency on mitochondrial metabolism.

Despite these shared dependencies, we also observe substantial heterogeneity in aneuploid vulnerabilities across cell types, karyotypic states, and cancer lineages. Certain dependencies are more pronounced in specific tissues or in cells harboring particular chromosome gains or losses, indicating that the precise manifestation of aneuploid stress is modulated by cellular context. This heterogeneity likely reflects differences in baseline metabolic programs, protein turnover, aneuploid karyotype, tissue specific expression, genetic background and compensatory mechanisms across cancer cell lines.

Additionally, our work has several limitations. While we have several net chromosome loss aneuploid clones in the DG screen, the WGL screens exclusively cover chromosome arm level gains. Our aneuploid gene group analysis may be limited to chromosome gain effects. Additionally, our Druggable Genome library is not a comprehensive library of all druggable genes, as several gene groups including phosphatases and transferases were not included. The DG screen did not identify any gene that was a significant hit across all cell lines (A2780, DLD1, HCT116 and SNU1) screened, highlighting heterogeneity in aneuploid cell lines.

We identify the ubiquitin E2–conjugating enzyme UBE2H as a top, and recurrent aneuploid-specific dependency. UBE2H dependency correlates with aneuploidy burden across our cancer cell lines and is a pan-cancer aneuploid dependency in DepMap. Additionally, UBE2H is upregulated by aneuploidy in both yeast and human cells, and its expression is associated with increased aneuploidy in human cancers. Aneuploid dependency on UBE2H was validated in isogenic competition assays and knockout-rescue experiments. Proteomic analyses further demonstrated that loss of UBE2H significantly reduced the abundance of mitochondrial transcription, translation, and electron transport chain proteins. These effects are exacerbated by treatment with chromosome instability-inducing drug AZ3146. These data suggest that UBE2H plays an important role in supporting mitochondrial protein homeostasis in aneuploid cells, potentially by facilitating the turnover or quality control of mitochondrial proteins under conditions of gene dosage imbalance.

The localization of UBE2H to mitochondria, together with its association with mitochondrial metabolic gene clusters in human tumors, further supports a model in which ubiquitin-mediated regulation is required to maintain mitochondrial function in aneuploid cells. A recent study also found that aneuploidy-induced proteostatic changes mediate the formation of mitochondrial precursor protein aggregates, reduce abundance of mitochondrial proteins, and impairs mitochondrial function^24^. While the precise substrates of UBE2H remain to be defined, our findings raise the possibility that aneuploid cells depend on enhanced mitochondrial protein degradation to cope with increased metabolic demand and imbalanced protein expression. More broadly, these results highlight a previously underappreciated connection between ubiquitin signaling and mitochondrial adaptation to aneuploidy.

In summary, our study provides a comprehensive and comparative view of aneuploid-specific genetic dependencies across diverse cancer contexts. By integrating paired CRISPR screens, large-scale dependency datasets, and functional validation, we identify a conserved network of gene groups—including proteostasis, RNA processing, ribosome biogenesis, and mitochondrial metabolism—that underpin aneuploid cell survival. These findings advance our understanding of how cancer cells tolerate chromosome-scale genomic imbalance and suggest that targeting mitochondrial metabolic gene groups may offer therapeutic opportunities for selectively exploiting aneuploidy in cancer.

## Methods

### Cell line culture

The identity of all cell lines was confirmed using short tandem repeat profiling. Cells were regularly tested for mycoplasma contamination. Unless otherwise stated, all cells were cultured at 5% CO_2_ at 37°C, with 1x Pen/Strep (Thermo Scientific, 15140-122), 1% L-Glutamine (Thermo Scientific, 25030-081) and 10% FBS (Gibco, A5256701) in associated cell culture media. DLD1, VACO432, A2058, and Cal51 were cultured in DMEM media (Thermo Scientific, 11995065). SNU1, and A2780 were cultured in RPMI 1640 (Thermo Scientific, 11875093) media. HCT116 and RKO were cultured in McCoy-5A (ATCC, 30-2007) media. AGS were grown in F-12K (ATCC, 30-2004) media. MCF10A cells were grown in DMEM/F12 (Invitrogen, 11330-032) media with 5% Horse Serum (Gibco, 16050-122), 1x Pen/Strep (Thermo Scientific, 15070-063), EGF (20 ng/ml, PeproTech, AF-100-15-500UG), Hydrocortisone (0.5 mg/ml, Millipore Sigma, H0888-5G), Cholera toxin (100 ng/ml, Millipore Sigma, C8052-1MG), Insulin (10 ug/ml, Millipore Sigma, I1882-100MG), and transferrin (5 ug/ml, Millipore Sigma, T8158-100MG).

Inducing aneuploidy in near-euploid cell lines:

### Induced Chromosome INstability (ICIN)

Near-euploid clones were grown with 1 or 2 µM AZ3146 for 24 hours. The drug was washed away, cells were allowed to recover for 24 hours in drug-free media and were then single cell sorted. Clones were grown and karyotyped to identify aneuploid chromosomes. For cell lines screened with the druggable genome, cells were transduced with a Cas9 vector (Addgene, 65655) before ICIN was performed.

### Microcell Mediated Chromosome Transfer (MMCT)

All Microcell Mediated Chromosome Transfer cell lines were received from the Storchova Lab, RPTU, Germany. The MMCT method is described in Kneissig M., et al. (2019)^37^. The generation of HCT116 Tetrasomy 5p was described in Stingele et al. (2012)^20^, and HCT116 Trisomy 8 was described in Donnelly et al. (2014)^63^.

### Restoring Disomy in Aneuploid Cells using CRISPR Targeting (ReDACT)

All ReDACT induced chromosome loss clones were generated in Girish et al. (2023)^16^. All three of the ReDACT methods were used to generate the four induced disomy subclones (positive-negative selection, artificial telomere, and CRISPR only; Supplementary Data 1). In AGS, positive-negative selection was used to generate disomy 1q, and the parental clone with the integrated selection cassette was used as the paired aneuploid clone. In all other cases, the wild-type parental cell lines were used.

### Karyotyping

To infer copy number alterations, samples were sequenced with whole genome sequencing at 0.5-1.0% coverage. A custom Nextflow wrapper (v1.0) for CNVKit (v0.9.11) was used to generate estimated copy number data^64–66^. Batches of whole genome sequencing were performed with “batch--method wgs” with a bin size of 100,000. The log_2_ copy number ratios of DNA reads were aligned with the human genome. Karyotyping plots were generated using a custom Python script (v3.12.4)^67^ with Matplotlib (v3.8.4)^68^ and pandas (v2.2.2)^69^. The script was adapted from CNVkit’s scatter.py with specific modifications to enhance visual representation. The workflow is available on GitHub: https://github.com/sheltzer-lab/Karyotyping.

### Generating the Druggable Genome library

The Druggable Genome library was generated by combining domain-focused gRNA sub-libraries targeting 482 kinases, 563 ubiquitin ligases, 598 epigenetic regulators, 1,427 transcription factors, and 528 proteases into a single pooled library. Among these, the kinase, transcription factors, and ubiquitin ligase sub-libraries were described in previous studies^70–72^. To allow blasticidin selection, GFP in the combined library was replaced with a GFP–P2A–BlastR cassette. After assembly, the library was sequenced using a NextSeq 500 to confirm that sgRNA abundance and distribution were maintained relative to the original library.

### CRISPR screens

For both the WGL and DG screens, cell lines were transduced at an estimated 0.3 Multiplicity Of Infection (MOI) to minimize dual guide transduction. Cells were screened in batches of 1-6 cell lines at the same time. A minimum of 500x coverage of the library was maintained throughout the screens for proper coverage. Cells were harvested three days after transduction (D=0) and again after they reached ten population doublings (D=10). Screens lasted between 12 and 37 days. We performed 14 batches of screens. Most (23/25) subclones were set relative to their batch specific controls. AGS 7p and DLD1 Ts2.18 (CRISPR screen batch 2) were set relative to AGS parental and DLD1 control 3, respectively.

The Whole Genome Library (WGL) was purchased from Addgene (73179-LV): the Human CRISPR Knockout Pooled Library (Brunello) virus lentiCRISPRv2^73^. The library covered 19,112 genes, with 4 guides per gene. The WGL contains a Puromycin resistance marker, sgRNAs, and Cas9. Puromycin resistance was used to estimate viral transduction efficiency in WGL screens. Before the start of the screen, cells were transduced with 5 different volumes of virus to estimate MOI in the cell line. Transduction efficiency was estimated based on puromycin resistance. Cell lines were Puromycin selected for 3 days after the first time point to enrich for sgRNA containing cells. If the cell line already contained Puromycin resistance, transduction efficiency was estimated by doing a test transduction on the wild-type version of that cell line and no selection took place.

The Druggable Genome (DG) library, also called the “Mega-A” multi-domain library, was expanded at the Yale Center for Molecular Discovery and the virus was generated using polyethylenimine (PEI) transfection in six 15 cm plates of 293FT cells. The DG library contained GFP, but did not contain Cas9 or a selection marker. All cell lines screened with the DG library were transduced with a Cas9 vector and single cell cloned before the start of the screen. To estimate the MOI in the cell line, control cells were transduced with 5 different volumes of virus, and transduction efficiency was estimated based on percent GFP expression. During the screen, GFP was measured in each cell line three days after transduction and measured again every time the cell line was passaged using the MACSQuant flow cytometer to maintain 500x coverage.

Genomic DNA was extracted from cell line pellets with JETQUICK Blood & Cell Culture DNA MAXI Spin Kit (A30706). For most screens, DNA was sent to Cellecta for sgRNA amplification and sequencing. For screen batches 1, 2 and 3, sgRNAs were amplified with barcoded Illumina adaptor sgRNA primers (P5 and P7) and purified via gel extraction. These PCR amplicons were sequenced with Illumina sequencing at the Yale Center for Genome Analysis (YCGA). The following Illumina sequencing primers were used:

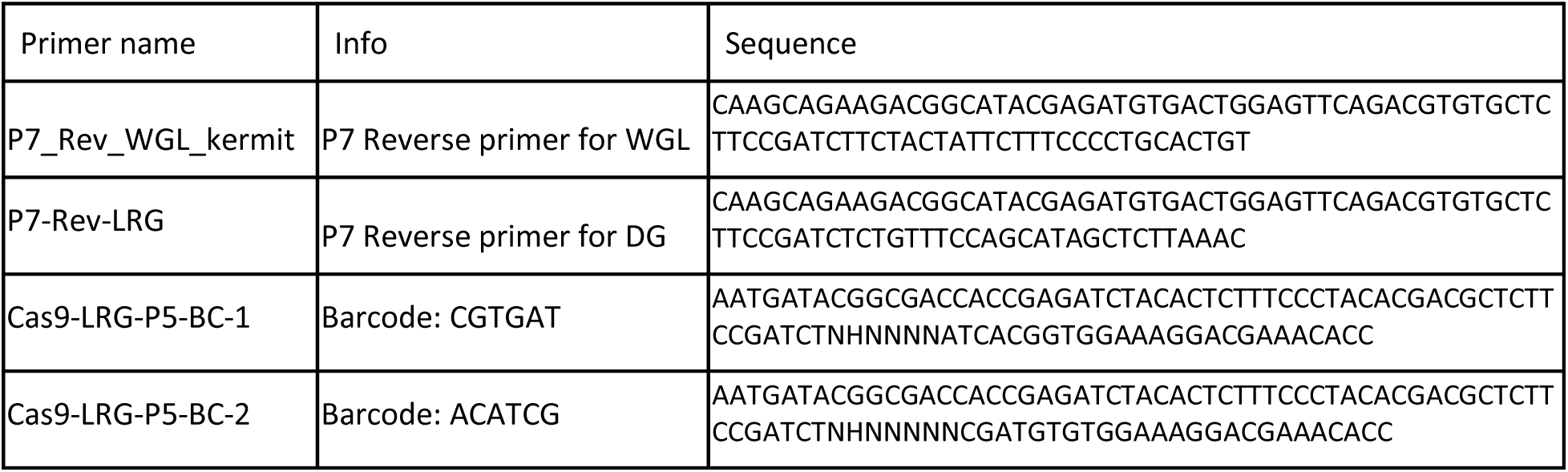
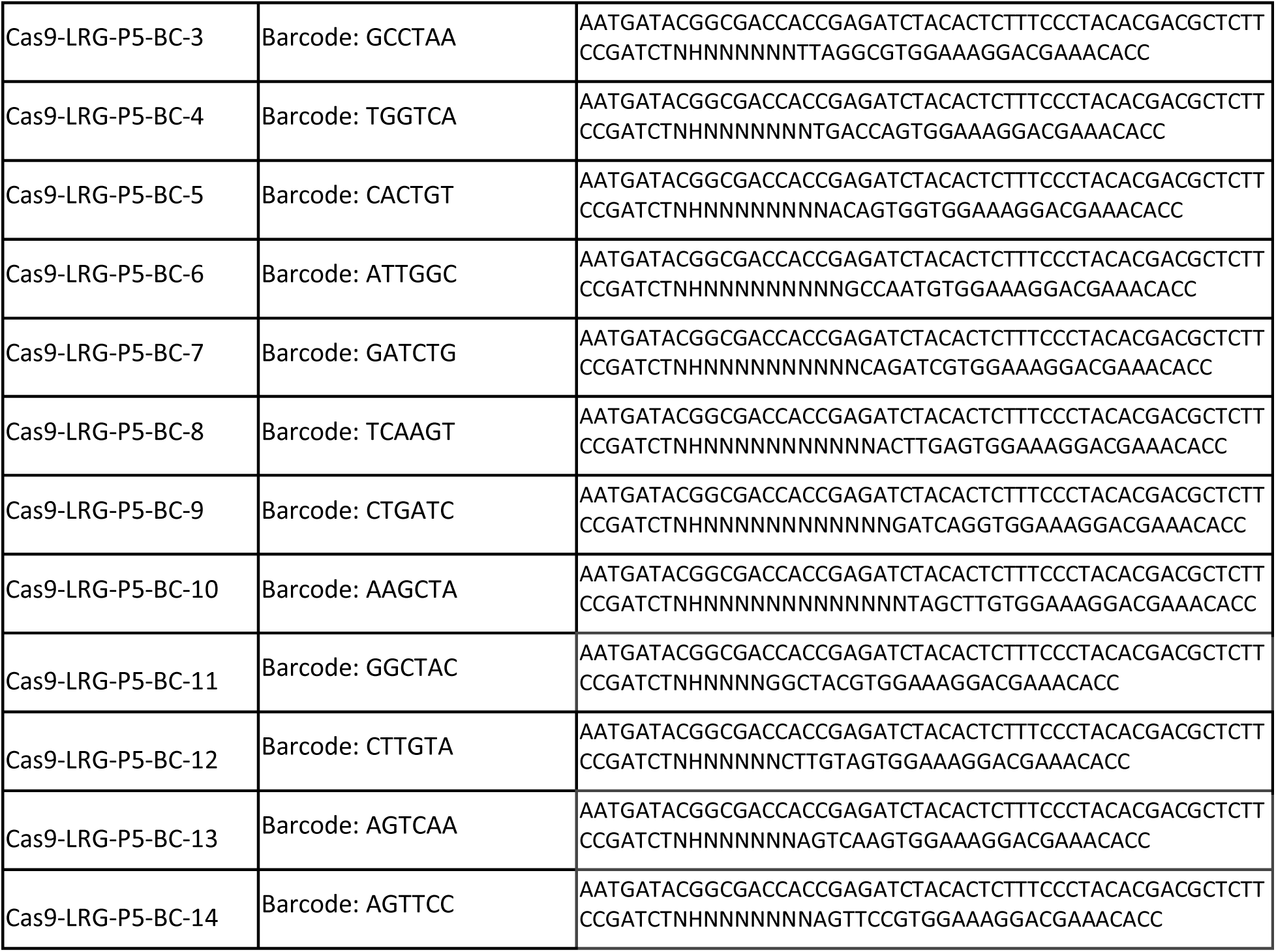

### CRISPR gene dropout analysis

Reads were extracted based on the sgRNA primary sequence and adapters. MAGeCK was used on read count data to analyze gene dropout and generate beta scores for each screen^38^. Quantile normalization was used to normalize beta scores across screens.

### Gene Set Enrichment Analysis and g:Profiler

G:Profiler values were calculated with R package “gprofiler2” v0.2.4^74^ and plotted using “ggplot2” v4.0.2.^75^ Enrichment cutoff for WGL paired gene difference was set at -0.2. Gene Set Enrichment Analysis was performed and visualized using R packages “DESeq2” v1.44.0^76^, “AnnotationDbi” v1.66.0^77^, “enrichplot” v1.24.4^78^, and “clusterProfiler” v4.12.6^79,80^. Samples were submitted as an ordered list.

### Cell gene dropout clustering analysis

The heatmap() function in R stats package v4.4.1^81^ was used to calculate and generate a dendrogram of cell clustering based on CRISPR screen beta scores. The princomp() function from R stats package v4.4.1 was used to generate the principal component analysis.

### Gene groups

The gene groups used in this paper were derived from published gene groups^82,83^ (downloaded 2025). Proteome genes were taken from HUGO group 690. Ribosome genes from HUGO group 1054, subset to include only cytoplasmic L and S ribosomal proteins. RNA processing genes originated from GO term GO:0006396 and subset into spliceosome genes with annotated terms "mRNA splicing, via spliceosome" or "regulation of mRNA splicing, via spliceosome"; RNA processing genes were also subset into rRNA processing genes with annotated terms "rRNA processing" or "regulation of rRNA processing". Mitochondrial translation and transcription genes are from combined HUGO groups 646, 1534, 1538, 2044, and 2289. Mitochondrial electron transport chain associated genes were from HUGO groups 639, 645, 1387, and 1853. The gene lists used are listed in Supplementary Data 6.

### Venn diagram

Top 10% of aneuploid-dependency gene overlap were identified and visualized with interactive InteractiVenn^84^ (Figure 4C).

### Western blots antibodies

Cell pellets of approximately 5 million cells were harvested and lysed in RIPA buffer with protease and phosphatase inhibitors. Lysates were shaken for an hour and spun down, and protein concentrations were determined by BCA protein assay kit (Thermo Scientific #A55865). Equal amounts (25 µg) of protein were loaded into mini-PROTEAN TGX 4-15% gels (BioRad #4568083). After running gels, protein was transferred to a blot, blocked in 5% milk W/V for 1 hour, and incubated overnight at 4°C with primary antibody. Secondary antibodies were incubated for one hour before washing and imaging. The following antibodies were used in this paper: 1:100 UBE2H antibody (Santa Cruz sc-515567), 1:5000 GAPDH-HRP antibody (Santa Cruz sc-365062 HRP), and 1:5000 secondary goat anti-mouse IgG HRP (Invitrogen 31430).

### Dropout assay

Dropout assays were performed as in Girish and Sheltzer (2020)^85^. As SNU1 cells are grown in suspension, 1µl of propidium iodide (PI; Thermo Scientific P3566) was added to samples before running samples through Cytoflex S and PI+ cells were excluded from further analysis. The dropout of the test gene sgRNAs were set relative to the controls, where the average fold dropout of the positive controls (CDK9 and CDC7) was set to 100% dropout, and the fold dropout of the negative controls (Rosa26 and AAVS1) set to 0%. sgRNA sequences were taken from previous publications^16,71,73,85^.

### Guide RNAs

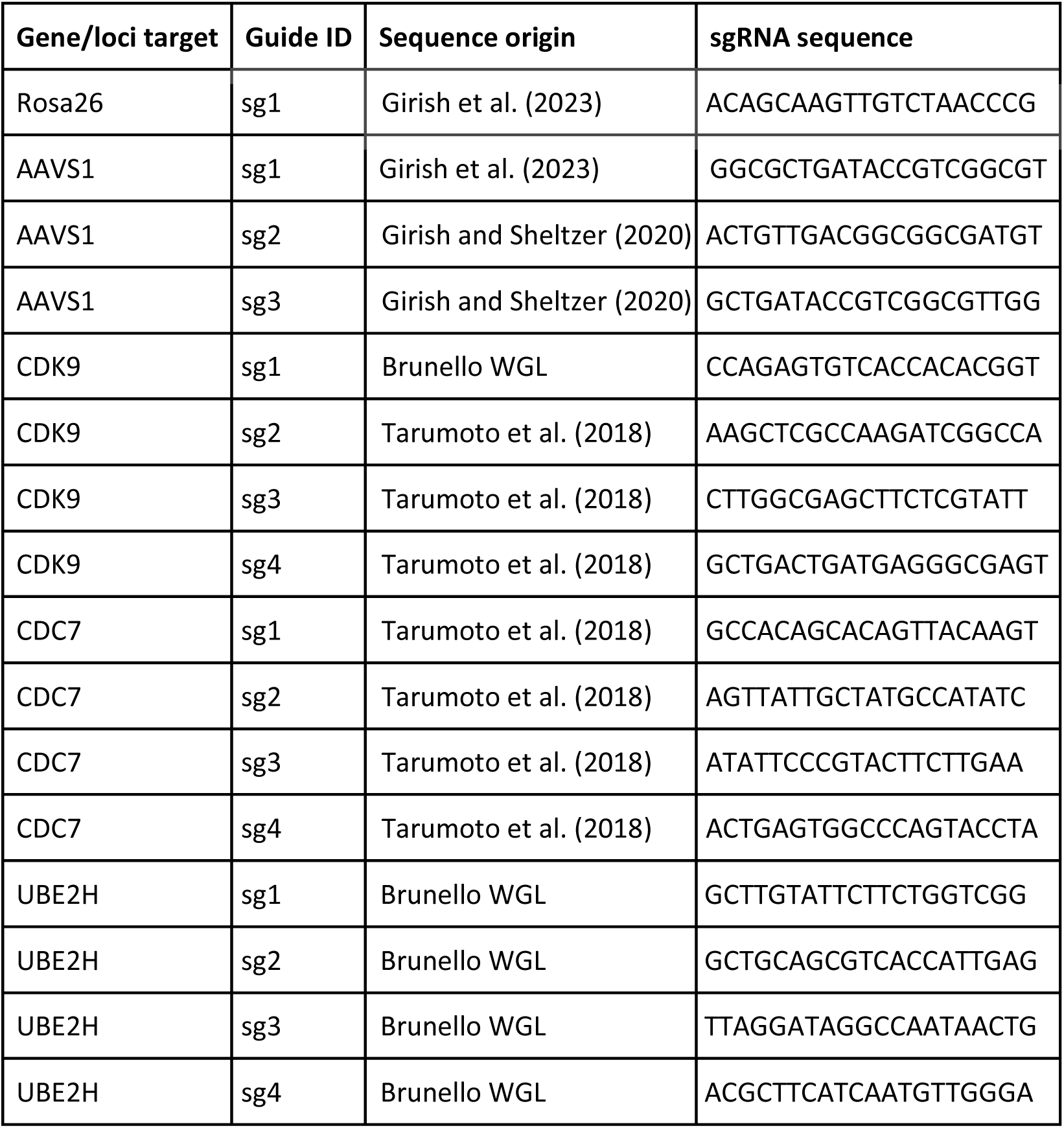

### Plasmids

pLenti Cas9 sgRNA vectors used in the dropout experiment and used to generate knockout clones were created by inserting target guide sequences into Lenti-spCas9-gRNA-GFP (Addgene #124770). The UBE2H cDNA overexpression vector was purchased from Genecopoeia (NM_003344.4). Lenti spCas9 PuromycinR vector (Addgene #65655) was used to integrate Cas9 into cell lines. Standard cloning techniques were used to generate Cas9-Hydro, by cloning a hygromycin resistance (from Addgene #80922) marker into the Cas9-Puro vector, replacing Puromycin resistance marker. Standard cloning techniques were used to clone UBE2H sg4 into pLenti Cas9 sgRNA mCherry (Addgene #108099).

### Cell line generation

To establish UBE2H knockout clones, HCT116 and Cal51 WT cells were transiently transfected with spCas9-gRNA-GFP UBE2H sg1 and spCas9-gRNA-mCherry UBE2H sg4 simultaneously^51^. Double-positive GFP+ mCherry+ cells were single cell sorted into 96-well plates to generate clones. Western blots were used to validate knockouts. Overexpression clones were generated by transducing cells with UBE2H cDNA vector and selecting for Puromycin (1 µg/ml) for 4 days. Western blot was used to confirm overexpression. PEI and polybrene were used for plasmid transfections. For transductions, helper plasmids pMD2.G VSVG (Addgene #12259) and psPAX2 (Addgene #12260) were co-transfected with plasmids.

### Colony formation assay

1,000 cells of each cell line were passaged into 6-well plates with 20% FBS and treated with 250 nM AZ3146 (Selleck Chemicals #S2731) or ethanol vehicle control. Cells were grown for 10 days until colonies were visible. Cells were washed with cold PBS, fixed with methanol, and stained with 0.5% crystal violet for 10 minutes. Colonies were washed, dried, and counted.

### Cell death staining

Cell lines were grown for 96 hours with 1 µM AZ3146 (Selleck Chemicals S2731) or ethanol vehicle control, in triplicate. Cells and media were harvested, resuspended in 100 µl annexin buffer and stained with AnnexinV FITC conjugate (Fisher Scientific, A13199) and then with 1 µl of DRAQ7 dead cell dye (Fisher Scientific, D15106). Cells were incubated for 20 minutes and 5 minutes, respectively. Samples were run through the Cytoflex S flow cytometer. Percent total death was measured by taking a sum of AnnexinV positive, DRAQ7 positive and dual positive cells. The difference in percent total death was measured by subtracting the mean total death of the ethanol vehicle control from the AZ3146 treated cells.

### Mass spectrometry analysis

Label-free mass spectrometry was performed by the Yale Keck Mass Spectrometry and Proteomics resource. Cells were treated with 1 µM AZ3146 or ethanol vehicle control for 96 hours before harvest. All conditions were analyzed in biological triplicate. Samples were processed by the Yale Keck Mass Spectrometry and Proteomics Resource. A background-based t-test was used to generate p-values and corrected using the Benjamini-Hochberg method.

## Material Availability

All cell lines associated with the paper are available upon request to the lead contact, Jason M. Sheltzer. Plasmids generated in this paper will be made available on Addgene.

## Data Availability

Raw MAGeCK beta scores for each CRISPR screen are available in Supplementary Data 2. sgRNA read counts and MAGeCK result files are available on GitHub. Proteomics data is available in Supplementary Data 9. Additional information required to reanalyze the data reported in this paper is available from the lead contact upon request. DepMap data were downloaded from the DepMap repository^39–44^, accession number 24Q2. DepMap aneuploidy scores were downloaded from Cohen-Sharir et al. (2021)^30^ and DepMap associated protein abundance data was downloaded from Nusinow et al. (2020)^45^. TCGA UBE2H expression data was downloaded December 2025 from cBioportal^58–60^, and TCGA aneuploidy data from Taylor et al. (2018)^57^.

## Code Availability

All code used to analyze CRISPR results and to plot figures are available on GitHub: https://github.com/kschukken/Aneuploid_CRISPR_Screens. The complete workflow used to generate karyotype images are available in a public repository on GitHub: https://github.com/sheltzer-lab/Karyotyping.

## Author Contributions

KMS: conceptualization, generated resources, investigation, validation, formal analysis and writing original draft. SMA, CZ, and PKK: investigation. RAH and SM: generated software and data curation. JK and ML: validation. TY, OK, ZS, and ELS: provided resources. CRV: supervision, provided resources. SJA: supervision, funding and manuscript review. JMS: conceptualization, methodology, supervision, funding and manuscript review.

## Competing Interest Statement

J.M.S. is an inventor on a patent related to chromosome engineering and the construction of aneuploid genomes. J.M.S. is a co-founder of and equity holder in Meliora Therapeutics and KaryoVerse Therapeutics. J.M.S. has received consulting fees from Merck, Pfizer, Ono Pharmaceuticals, Highside Capital Management, and Meliora Therapeutics and is a member of the advisory boards of BioIO, Permanence Bio, KaryoVerse Therapeutics, and the Chemical Probes Portal. C.R.V. has received consulting fees from Flare Therapeutics, Roivant Sciences, and C4 Therapeutics. C.R.V. has served on the advisory boards of KSQ Therapeutics, Syros Pharmaceuticals, and Treeline Biosciences. C.R.V. has received research funding from Boehringer Ingelheim and Treeline Biosciences; and owns stock in Treeline Biosciences.

## Acknowledgements

We thank the Mass Spectrometry & Proteomics Resource at Yale University for providing the necessary mass spectrometers and the accompanying biotechnology tools, funded in part by the Yale School of Medicine and by the Office of the Director, National Institutes of Health (S10OD02365101A1, S10OD019967, and S10OD018034). We would like to thank the Yale Center for Genome Analysis for sequencing part of the CRISPR screens; Research reported in this publication was supported by the National Institute of General Medical Sciences of the National Institutes of Health under Award Number 1S10OD030363-01A1. We thank the Yale Center for Molecular Discovery for their assistance with expanding the Druggable Genome CRISPR library. The Core is supported in part by an NCI Cancer Center Support Grant # NIH P30 CA016359. This work was supported by the USA National Institutes of Health (NIH) F32 fellowship 1F32CA265041 (to K.M.S), a Cancer Research UK Clinician Scientist Fellowship RCCCSF-May23/100001 (to S.J.A.), and by a grant from ONO pharmaceuticals Co. Ltd. (J.M.S.).

## Supplementary Data Files

Supplementary Data 1- Aneuploid clones and sgRNA libraries

Supplementary Data 2- Karyotypes for each clone at start and end of screen

Supplementary Data 3- CRISPR Screen dropout results

Supplementary Data 4- WGL Aneuploid dependency gene set enrichment

Supplementary Data 5- DepMap aneuploidy score correlation

Supplementary Data 6- Gene lists

Supplementary Data 7- Total aneuploidy burden correlation

Supplementary Data 8- Ranked top hit genes per dataset

Supplementary Data 9- Proteomics

## Notes

https://github.com/kschukken/Aneuploid_CRISPR_Screens

https://github.com/sheltzer-lab/Karyotyping

