## Supplementary Figure 1 for "Paired CRISPR screens identify mitochondrial metabolism and UBE2H as aneuploid-specific dependencies in human cancer cell lines"

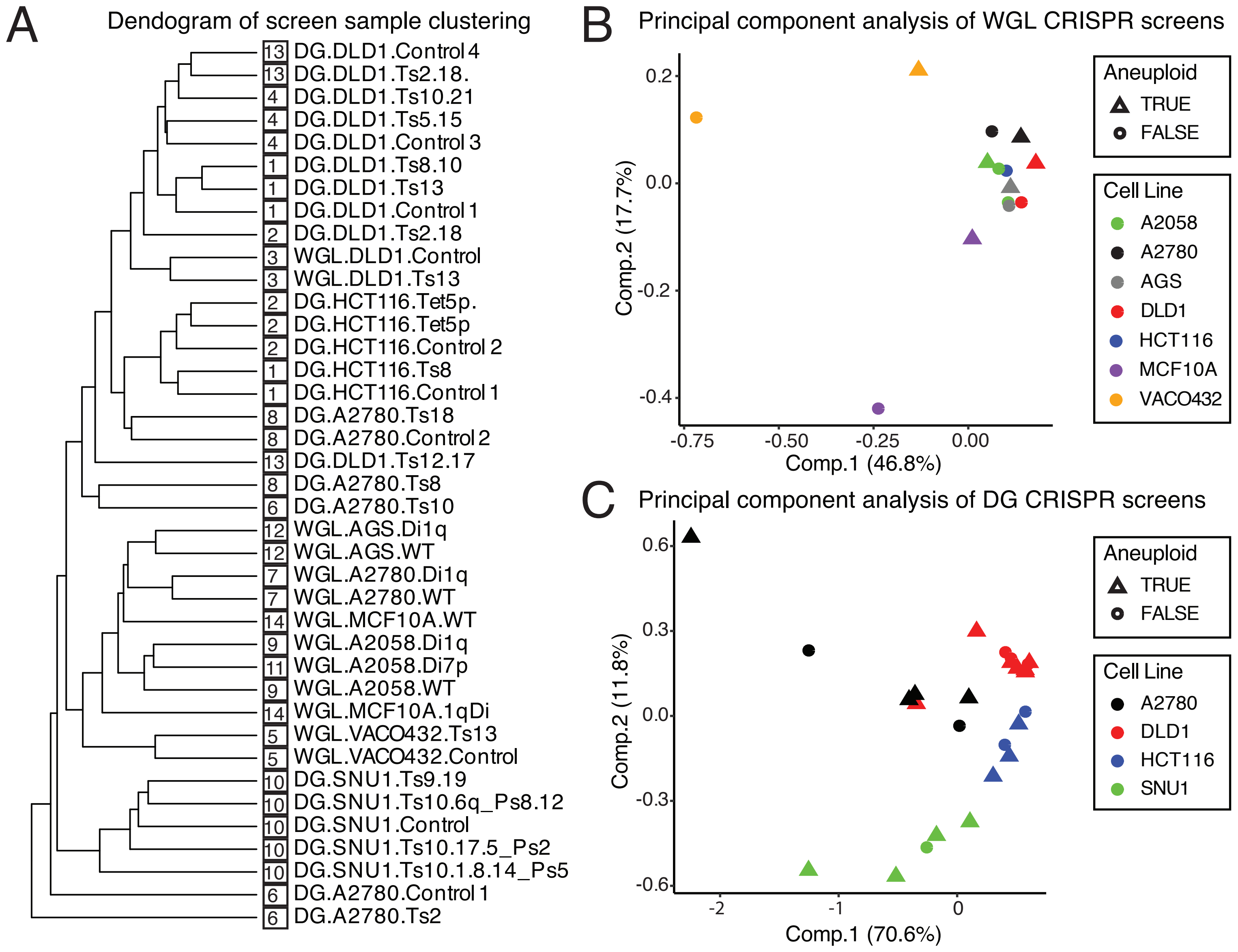


Figure S1: CRISPR screen cell cluster
A) Hierarchical clustering dendrogram of cell lines based on normalized CRISPR beta scores. CRISPR screen batch indicated by boxed number. B–C) Principal component analysis of normalized CRISPR beta scores for B) WGL and C) DG screens. Samples are labeled by cell line and relative aneuploidy status.
