## Supplementary Figure 2 for "Paired CRISPR screens identify mitochondrial metabolism and UBE2H as aneuploid-specific dependencies in human cancer cell lines"

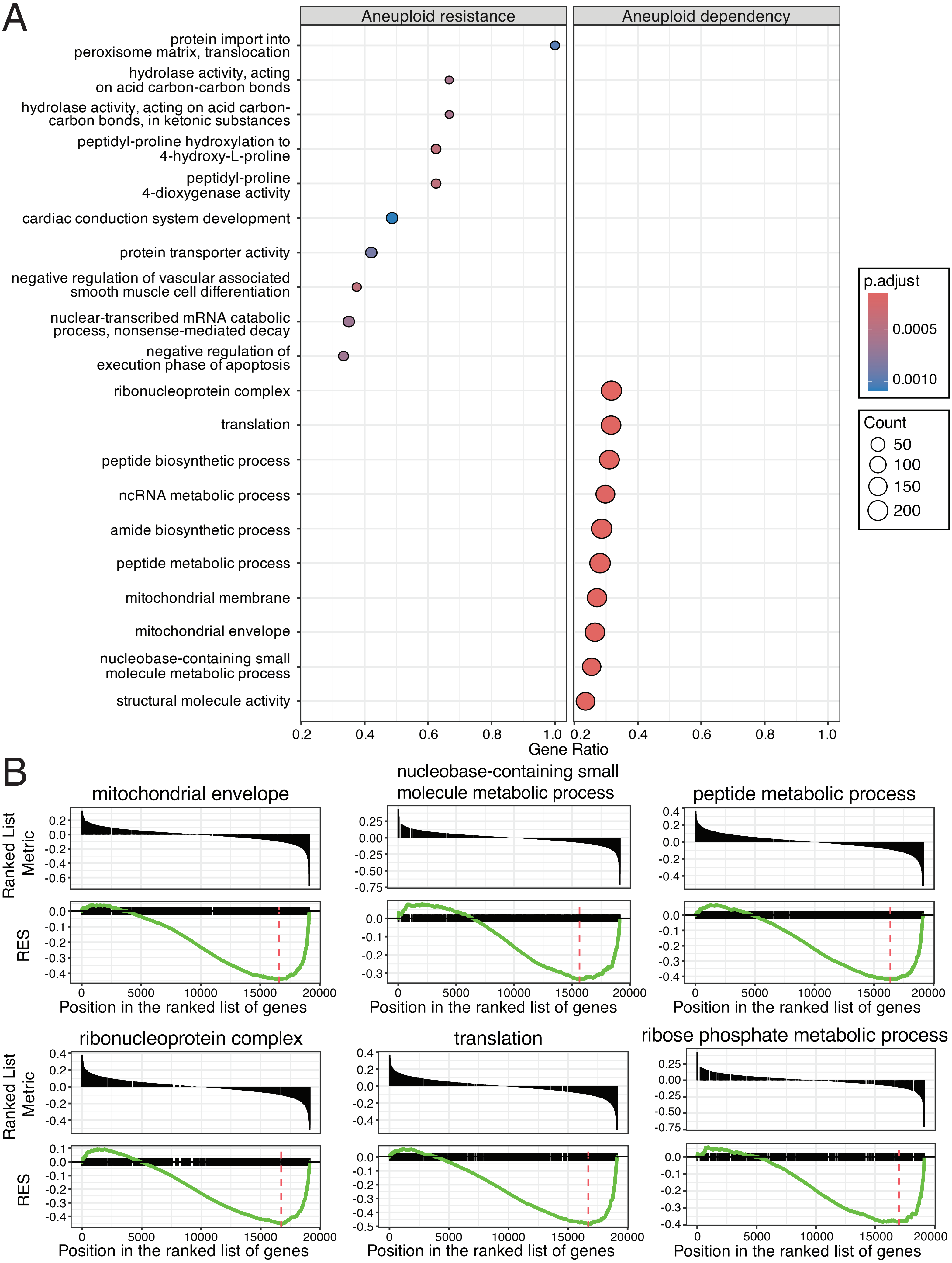


Figure S2: Aneuploid dependency gene groups in WGL screen

A) Top ten gene sets enriched for aneuploid dependency or resistance based on GSEA of paired WGL data. Dot size indicates gene set size; color denotes adjusted p-value. B) GSEA enrichment plots for six significant aneuploid dependency gene groups.
