## Supplementary Figure 3 for "Paired CRISPR screens identify mitochondrial metabolism and UBE2H as aneuploid-specific dependencies in human cancer cell lines"

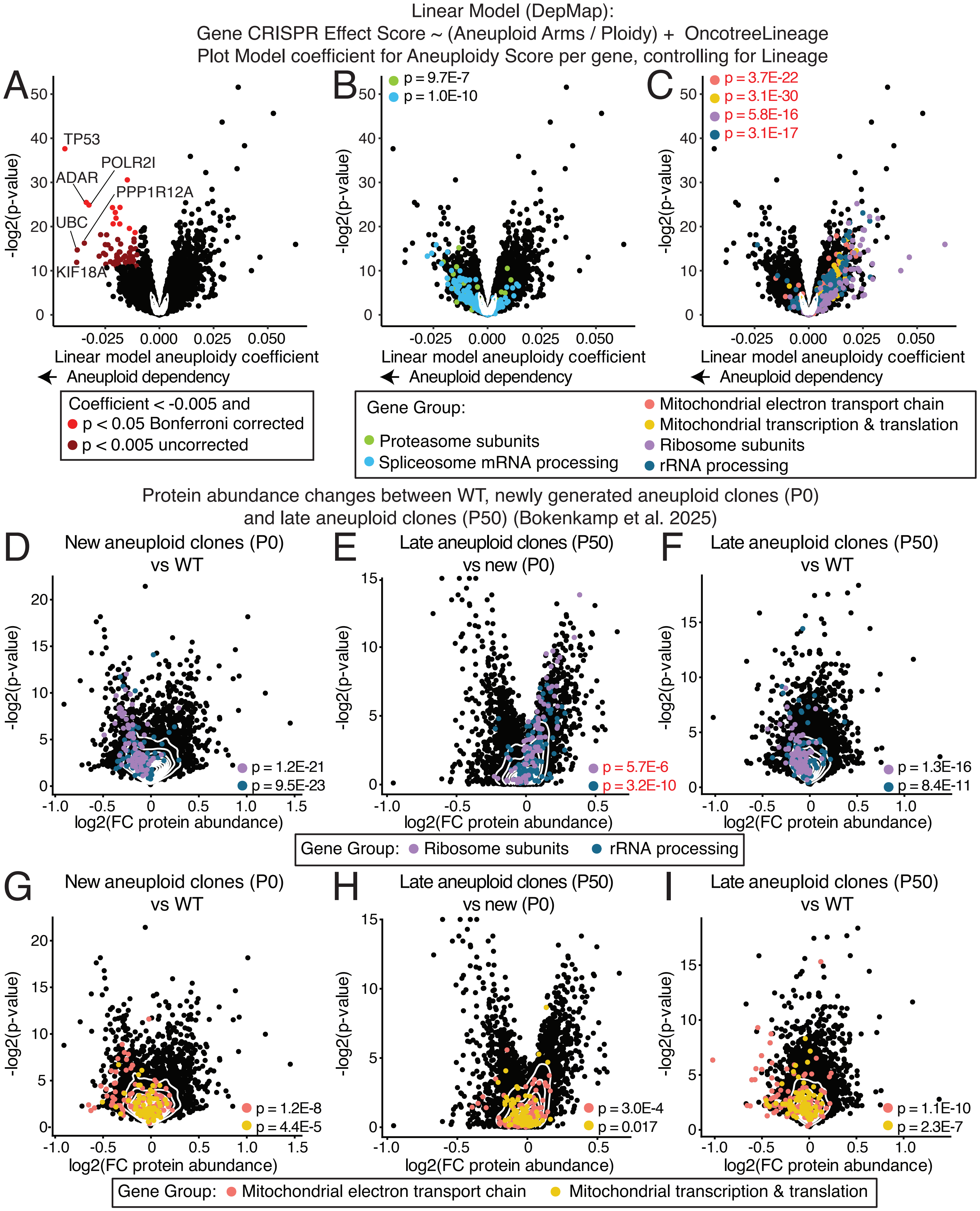


Figure S3: Ribosomal and mitochondrial dependencies in adapted aneuploid cell lines
A–C) Multivariate linear regression analysis of DepMap data showing the effect of aneuploidy score on CRISPR Effect Score while controlling for tumor lineage. Aneuploidy score defined as number of aneuploid arms corrected for ploidy. Negative coefficients indicate increased dependency in aneuploid cell lines. A) Significant genes (Bonferroni-adjusted $p < 0.05$) are shown in red; trends (uncorrected $p < 0.005$) in dark red. B–C) Genes belonging to aneuploid dependency gene groups identified in paired screens are highlighted. D–I) $Log_2$ fold changes in protein abundance from Bökenkamp et al. (2025) comparing newly generated aneuploid subclones (passage 0) with wild-type cells (D, G), late passage (P50) versus passage 0 (E, H), and late passage versus wild-type (F, I). Samples highlighted by gene groups. Two-sided t-test p-values compare each gene group to all other proteins. Black text indicates negative fold changes or coefficients; red text indicates positive fold change or coefficients.
