## Supplementary Figure 4 for "Paired CRISPR screens identify mitochondrial metabolism and UBE2H as aneuploid-specific dependencies in human cancer cell lines"

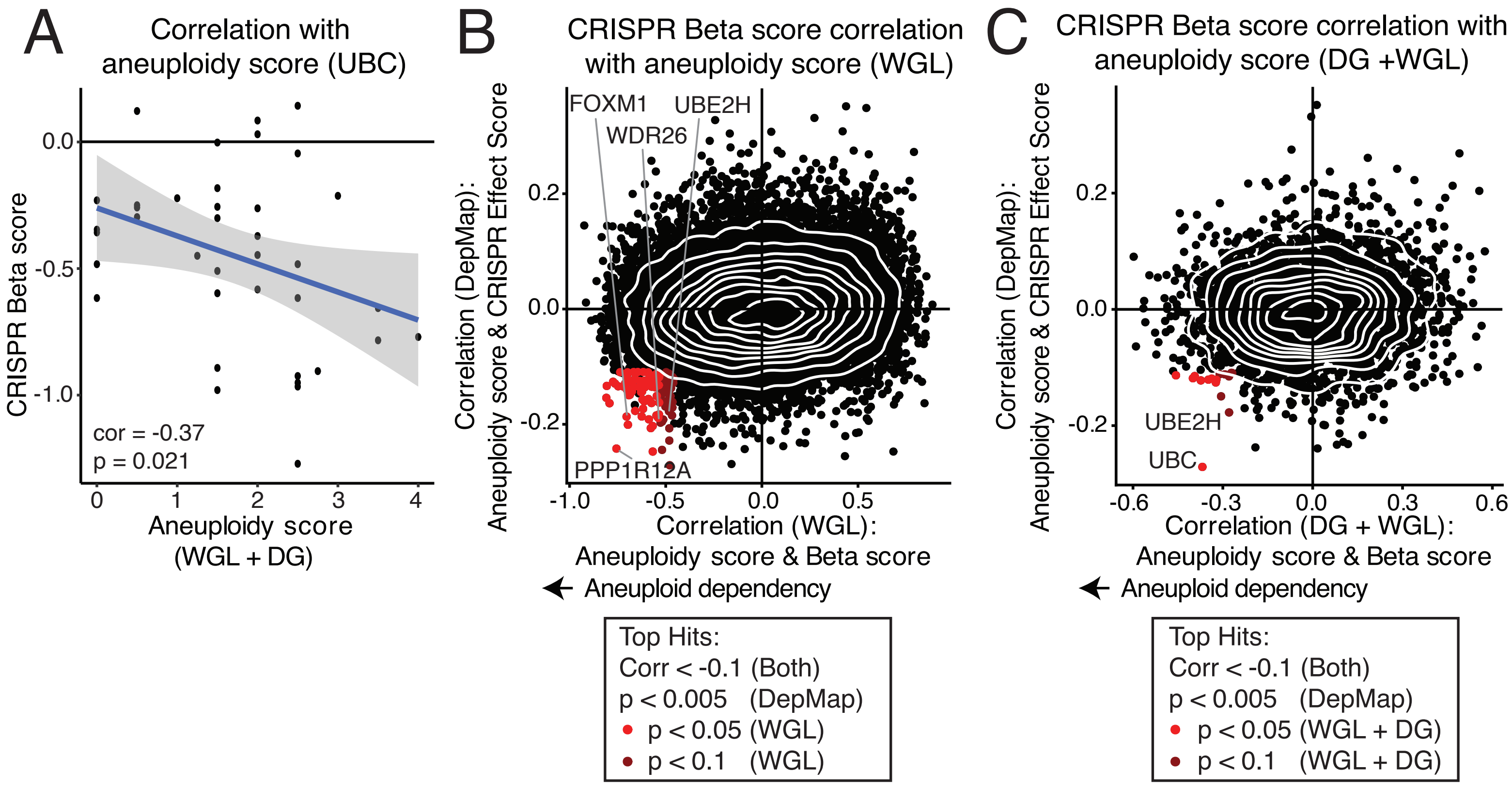


Figure S4: Aneuploidy burden correlation analysis

A) Pearson correlation between aneuploidy score and CRISPR beta score for UBC in paired WGL and DG screens. Aneuploidy score defined as number of aneuploid arms corrected for ploidy. B–C) Pearson correlations between aneuploidy score and gene dependency in B) DepMap versus WGL screens and C) DepMap versus merged WGL and DG screens. Genes with correlation < –0.1 in both datasets and meeting indicated p-value thresholds are highlighted.
