## Supplementary Figure 5 for "Paired CRISPR screens identify mitochondrial metabolism and UBE2H as aneuploid-specific dependencies in human cancer cell lines"

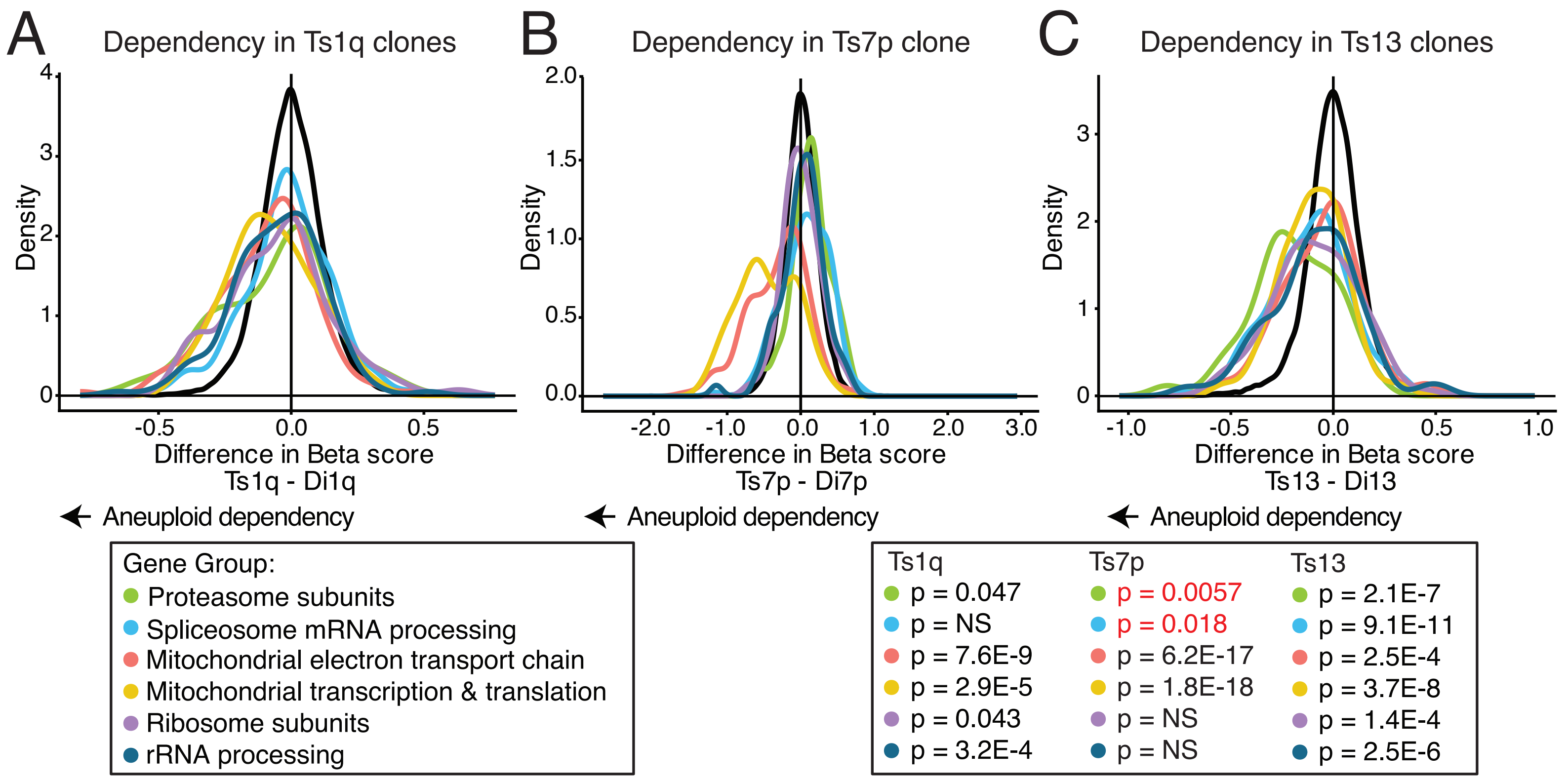


Figure S5: Gene group dependency per aneuploid arm of the WGL paired screen
A-C) Density plot of the paired difference in beta score per gene group in A) trisomy 1q subclones B) the trisomy 7p subclone and C) trisomy 13 subclones. P-values from two-sided t-tests of gene group beta scores versus non gene group scores. Black text denotes aneuploid dependency; red text denotes aneuploid resistance.
