## Supplementary Figure 6 for "Paired CRISPR screens identify mitochondrial metabolism and UBE2H as aneuploid-specific dependencies in human cancer cell lines"

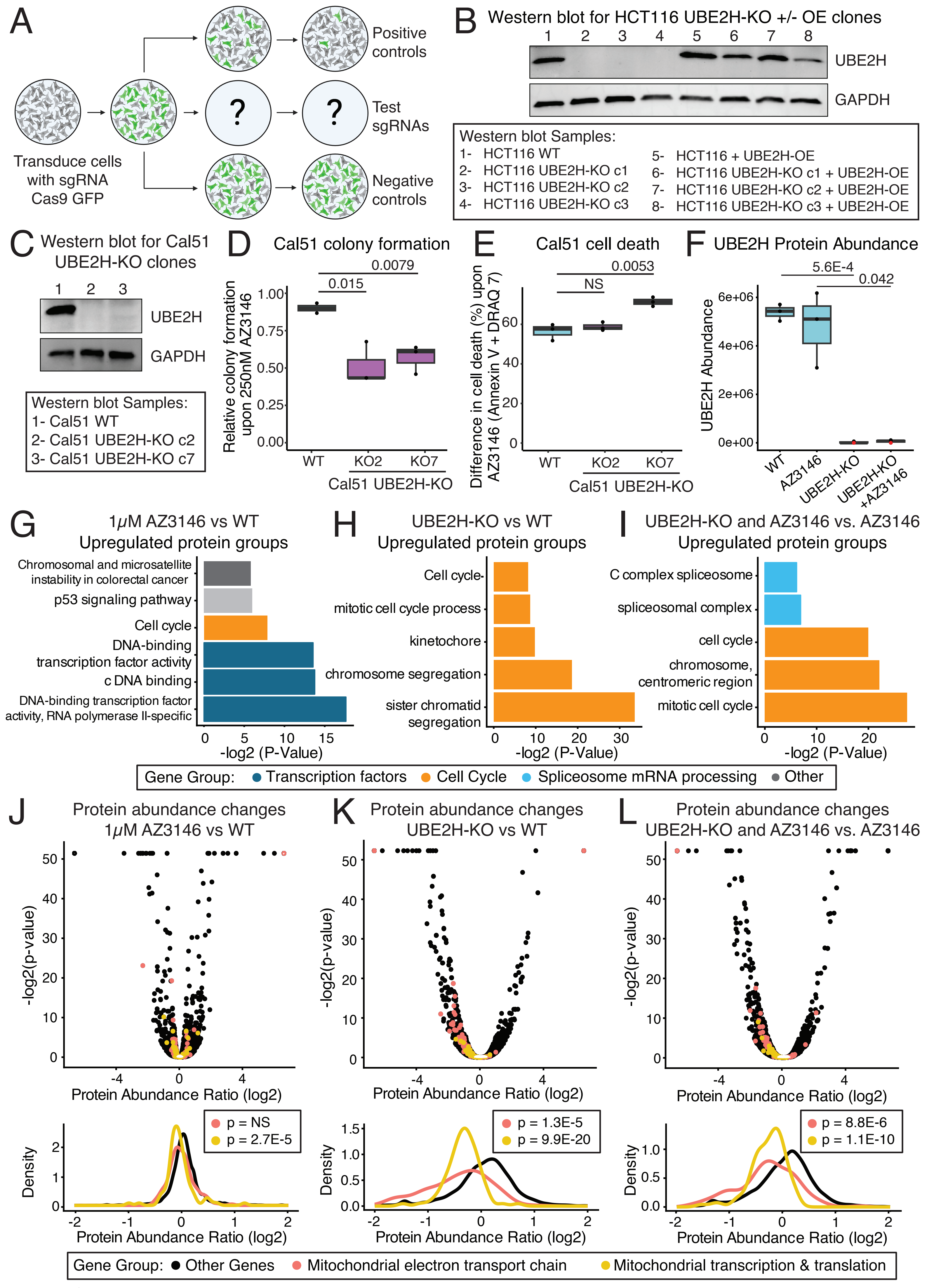


Figure S6: UBE2H functional validation and proteomics

Figure S6: UBE2H functional validation and proteomics

A) Schematic of competition assay used to measure sgRNA dropout relative to positive and negative controls over time. B) Western blot for HCT116 WT, UBE2H knockout clones and UBE2H cDNA overexpression clones. C) Western blot for Cal51 WT and UBE2H knockout clones. D) Colony formation assay in Cal51 WT and UBE2H knockout clones following 250nM AZ3146 treatment. Two-sided t-tests. E) Change in percentage of Annexin V and/or DRAQ7-positive Cal51 cells following 96 h treatment with AZ3146 (1 $\mu$M), relative to ethanol-treated controls; two-sided t-test. F) UBE2H protein abundance across conditions; samples with undetectable protein are indicated in red. Benjamini-Hochberg corrected background-based p-values. All samples were run in triplicate. G-I) g:Profiler gene group enrichment analysis of mass spectrometry data. Gene groups of upregulated proteins in HCT116 for G) AZ3146 treated versus wild-type, H) UBE2H-KO versus wild-type and I) AZ3146 treated UBE2H-KO versus AZ3146 treated wild-type cells. J-L) Label-free quantitative mass spectrometry analysis of protein abundance. Scatter (top) and density (bottom) plots show $\log_2$ protein abundance ratios for J) AZ3146 treated versus untreated HCT116 cells, K) UBE2H knockout versus wild-type HCT116 cells, and L) AZ3146 treated UBE2H knockout versus AZ3146 treated wild-type HCT116 cells. P-values from two-sided t-test of gene abundance ratios of genes in a gene group relative to genes not in the group.
