## Supplementary Figure 7 for "Paired CRISPR screens identify mitochondrial metabolism and UBE2H as aneuploid-specific dependencies in human cancer cell lines"

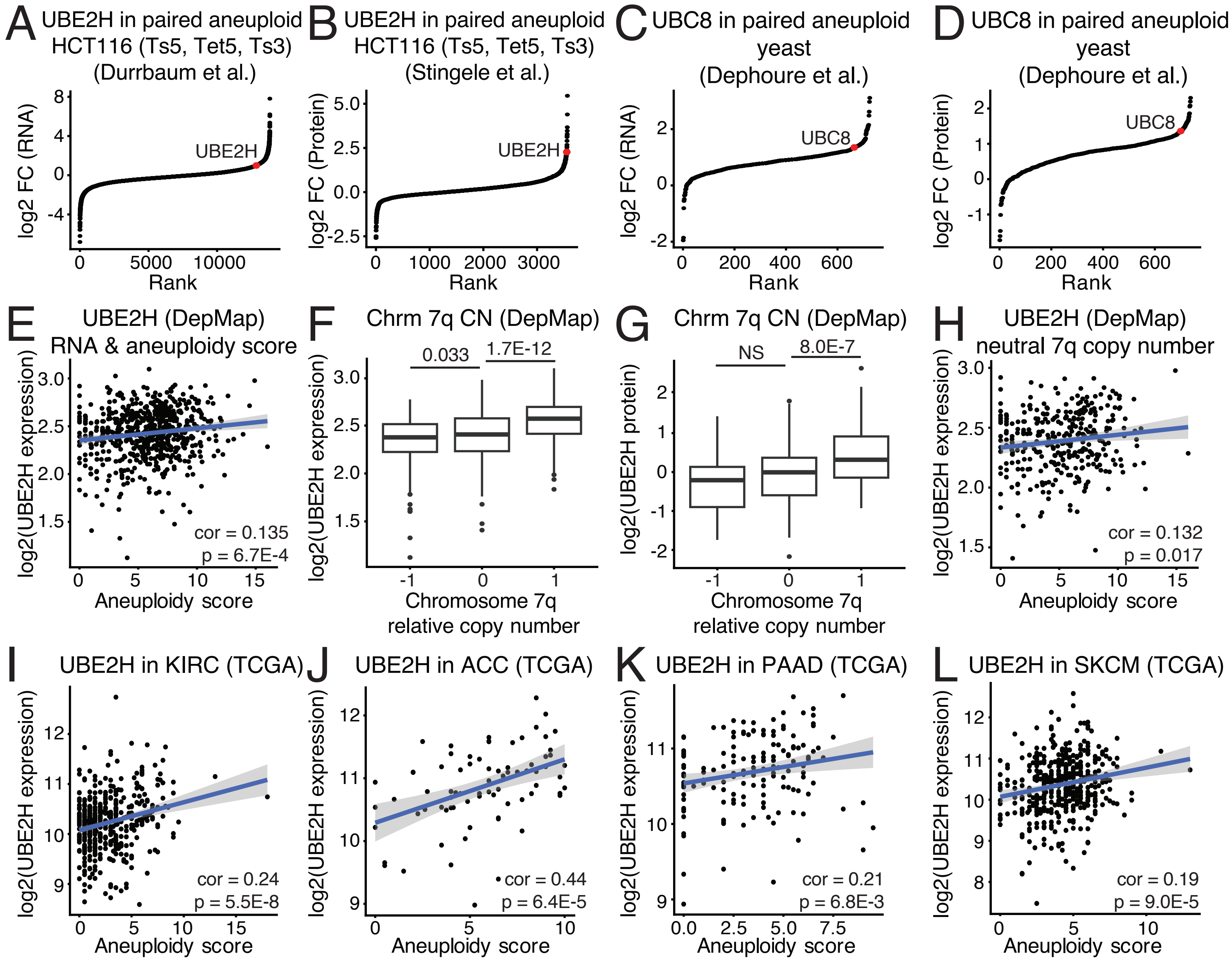


Figure S7: UBE2H expression correlates with aneuploidy in yeast, cell lines, and tumors

A-B) Genes ranked by mean log2 fold change of A) RNA expression or B) Protein abundance in aneuploid HCT116 clones relative to wild-type cells. Aneuploid clones were Trisomy 5, Tetrasomy 5, and Trisomy 3. C-D) Genes ranked by mean log2 fold change in C) RNA expression and D) protein abundance in disomic yeast relative to wild-type. E) UBE2H RNA expression correlated to aneuploidy score in DepMap cancer cell line data. Pearson correlation is shown. F-G) UBE2H RNA expression or protein abundance stratified by chromosome arm 7q copy number status; with 7q arm loss (-1), neutral (0), and gain (1) in DepMap data. H) UBE2H RNA expression correlated to aneuploidy score for samples with a neutral copy of chromosome arm 7q in DepMap data. I-L) UBE2H RNA expression Pearson correlated with aneuploidy score in TCGA data, subset by cancer type: I) kidney renal clear cell carcinoma (KIRC), J) adrenocortical carcinoma (ACC), K) pancreatic adenocarcinoma (PAAD), and L) skin cutaneous melanoma (SKCM).
