## Supplementary Data 2 for "Paired CRISPR screens identify mitochondrial metabolism and UBE2H as aneuploid-specific dependencies in human cancer cell lines"

### DG: HCT116 WT

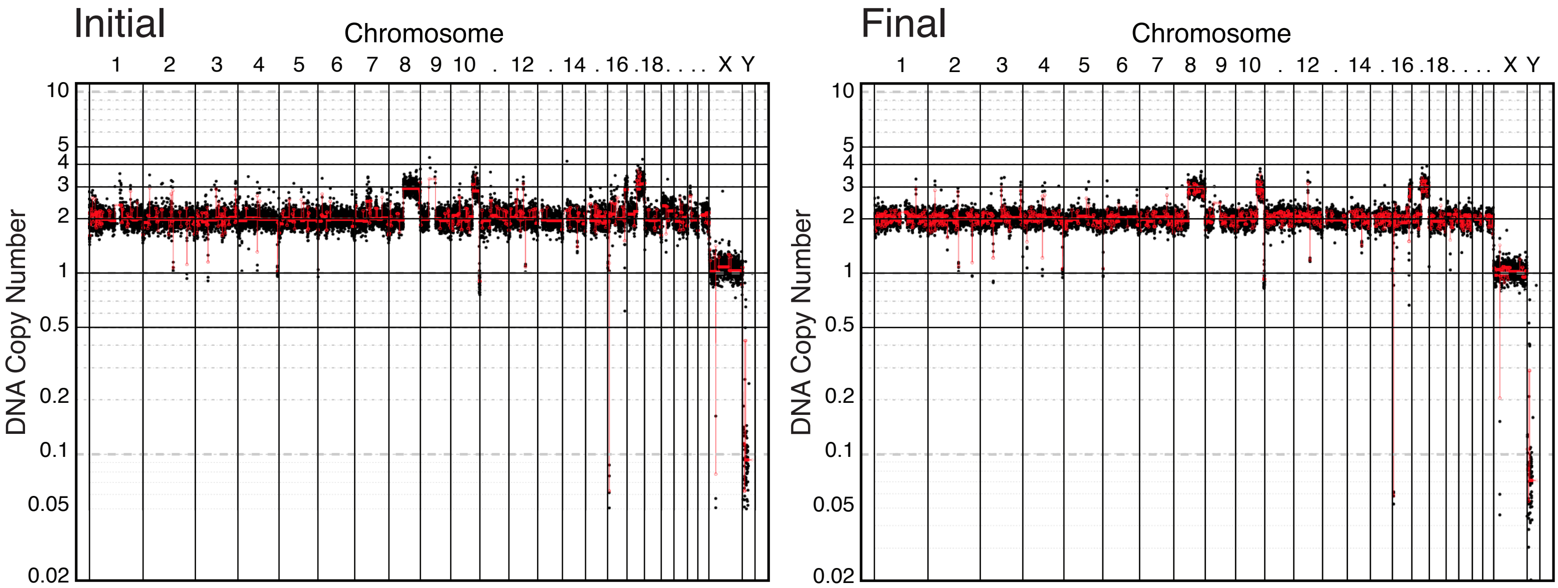


### DG: HCT116 Ts8

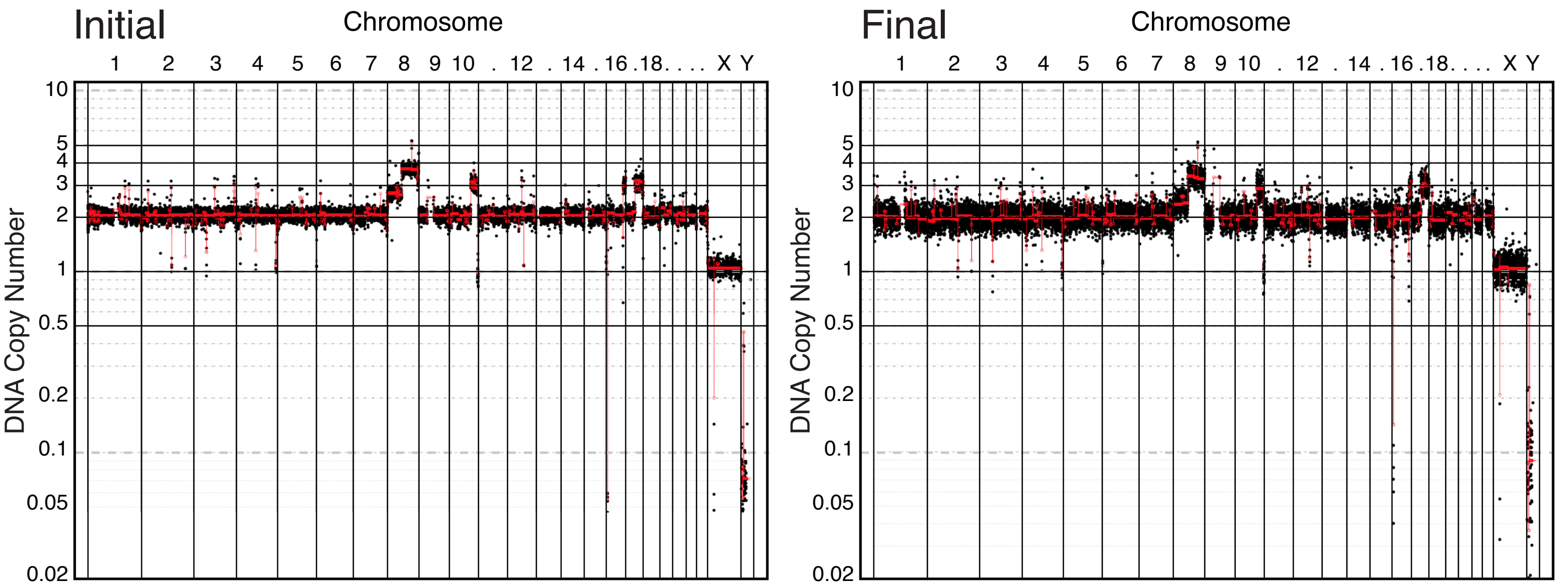

### DG: HCT116 Control 2

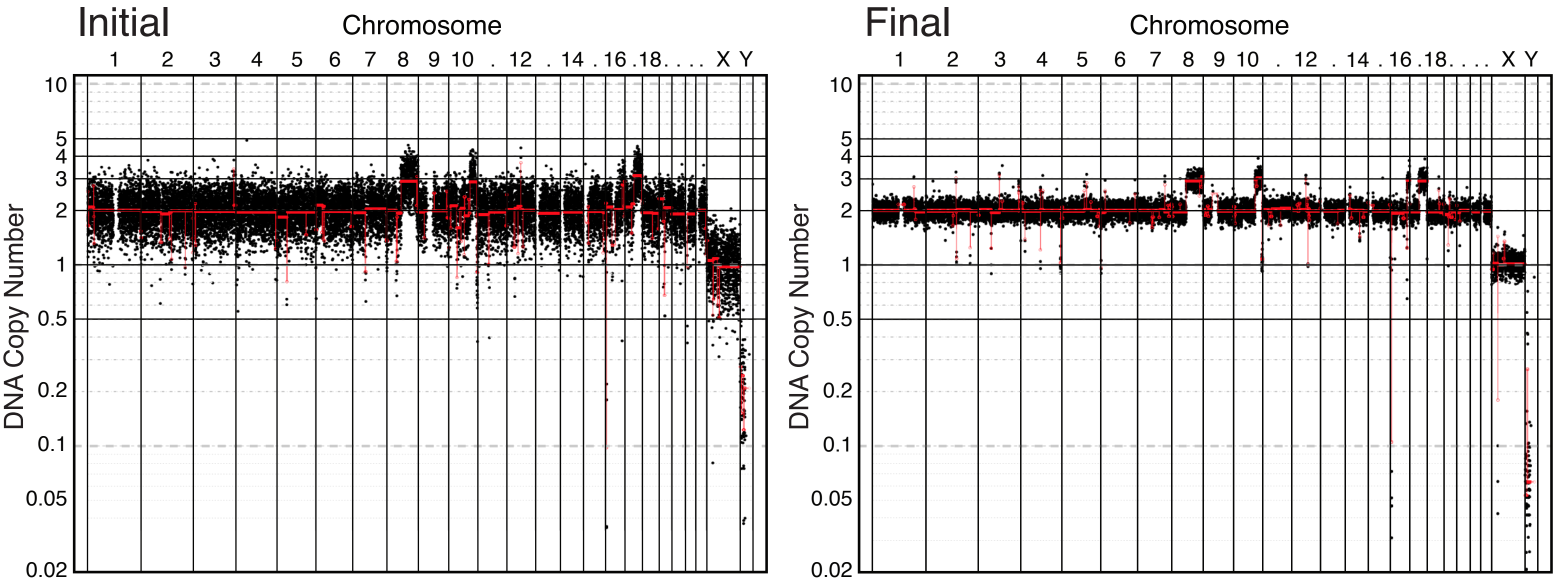


### DG: HCT116 Tet5p replicate 1

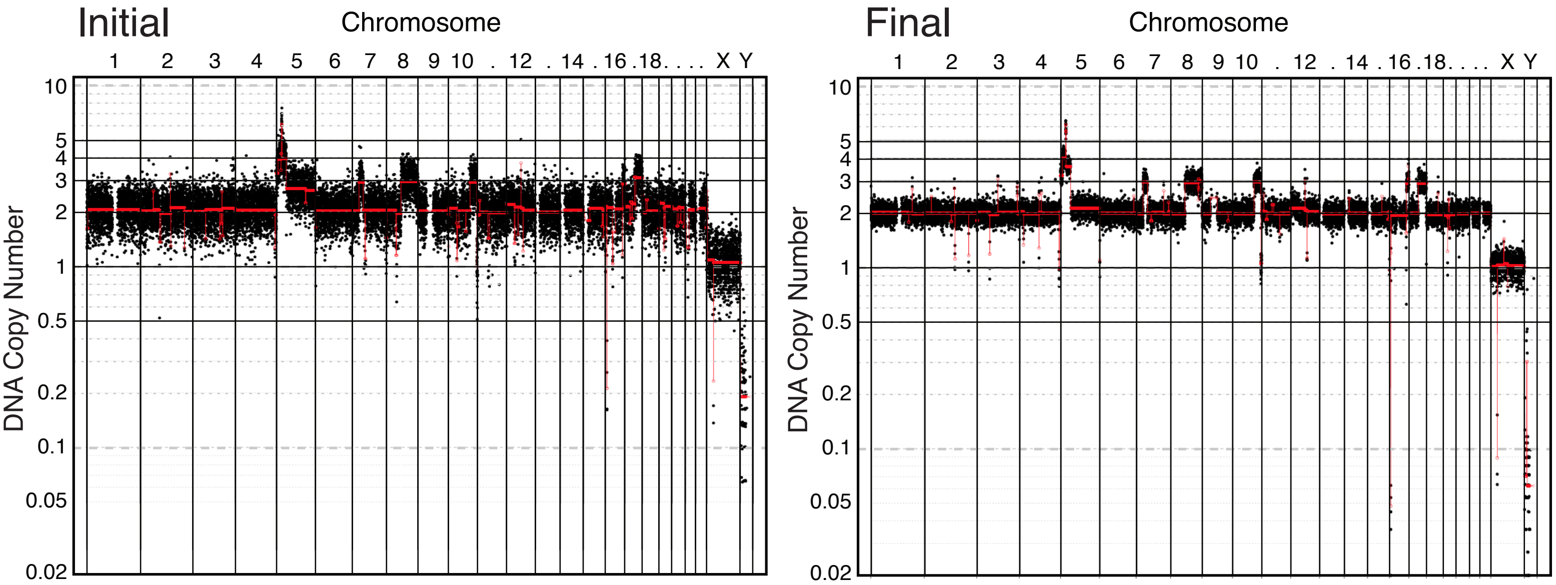


### DG: HCT116 Tet5p replicate 2

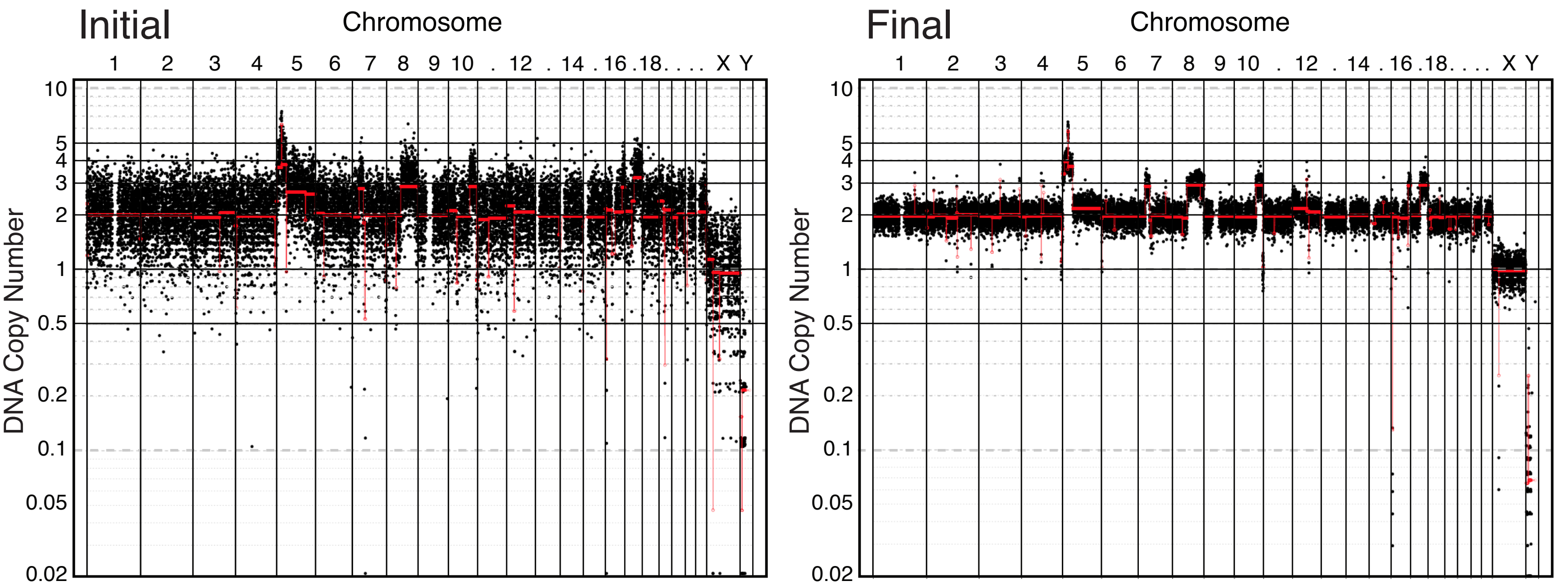

### DG: DLD1 Control 1

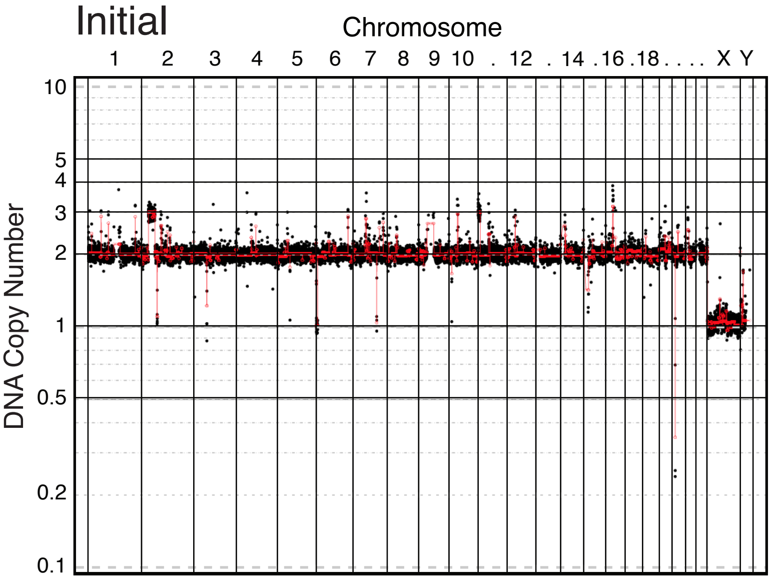


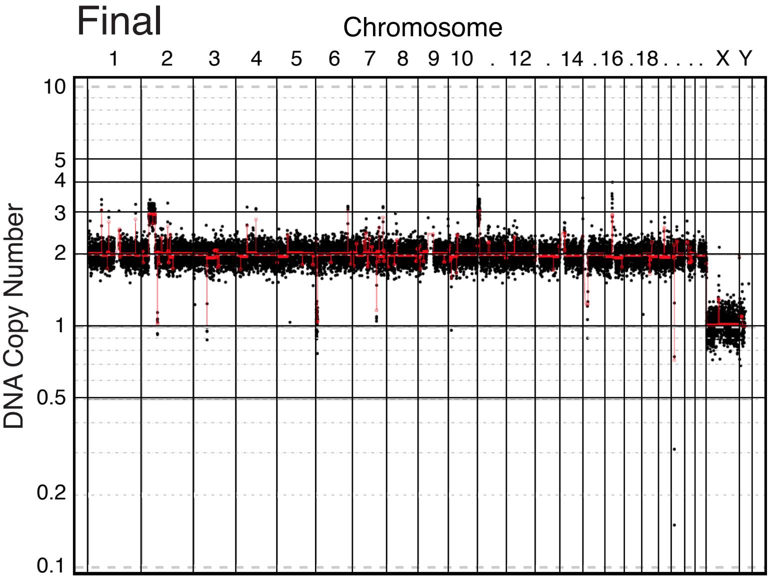


### DG: DLD1 Ts13

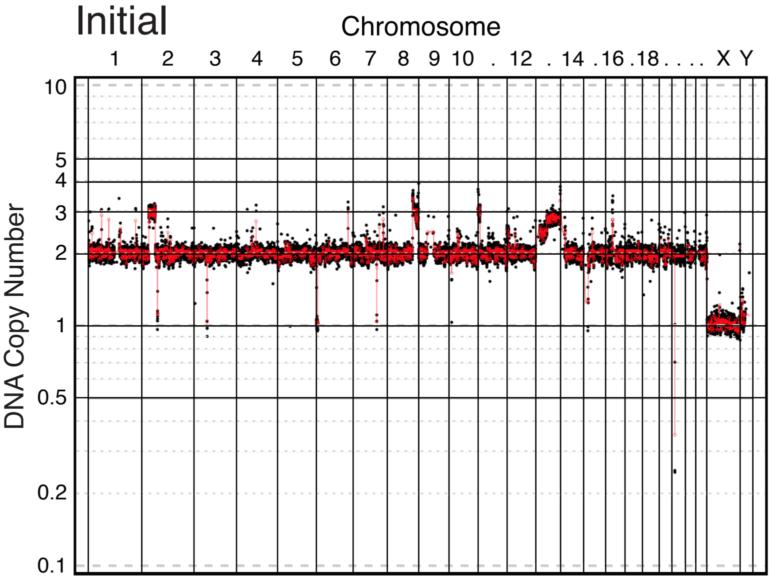


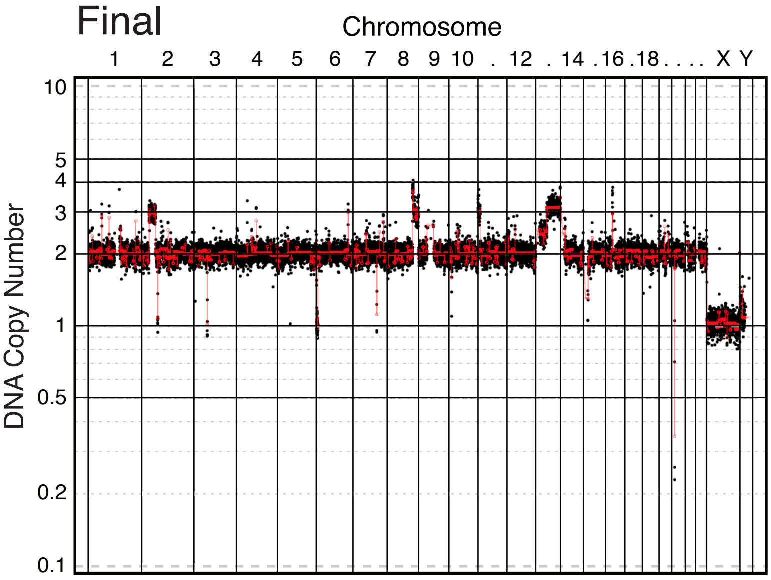


### DG: DLD1 Ts8.10

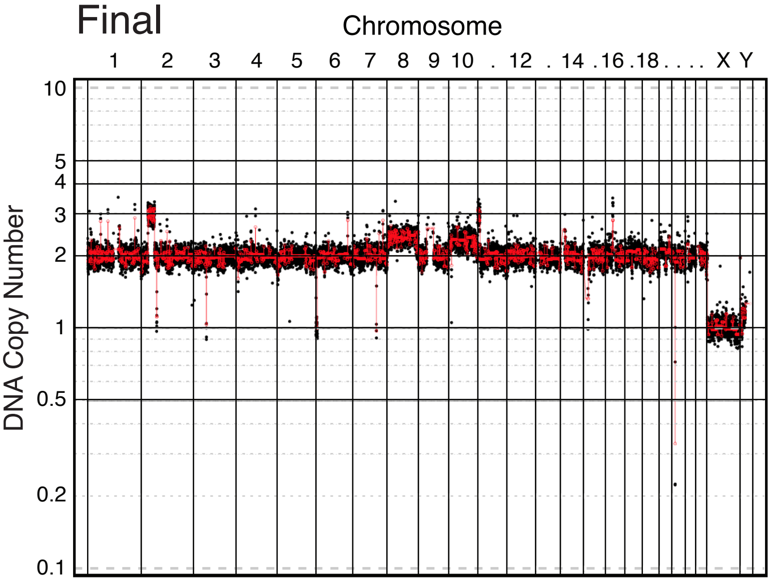

### DG: DLD1 Ts2.18 replicate 1

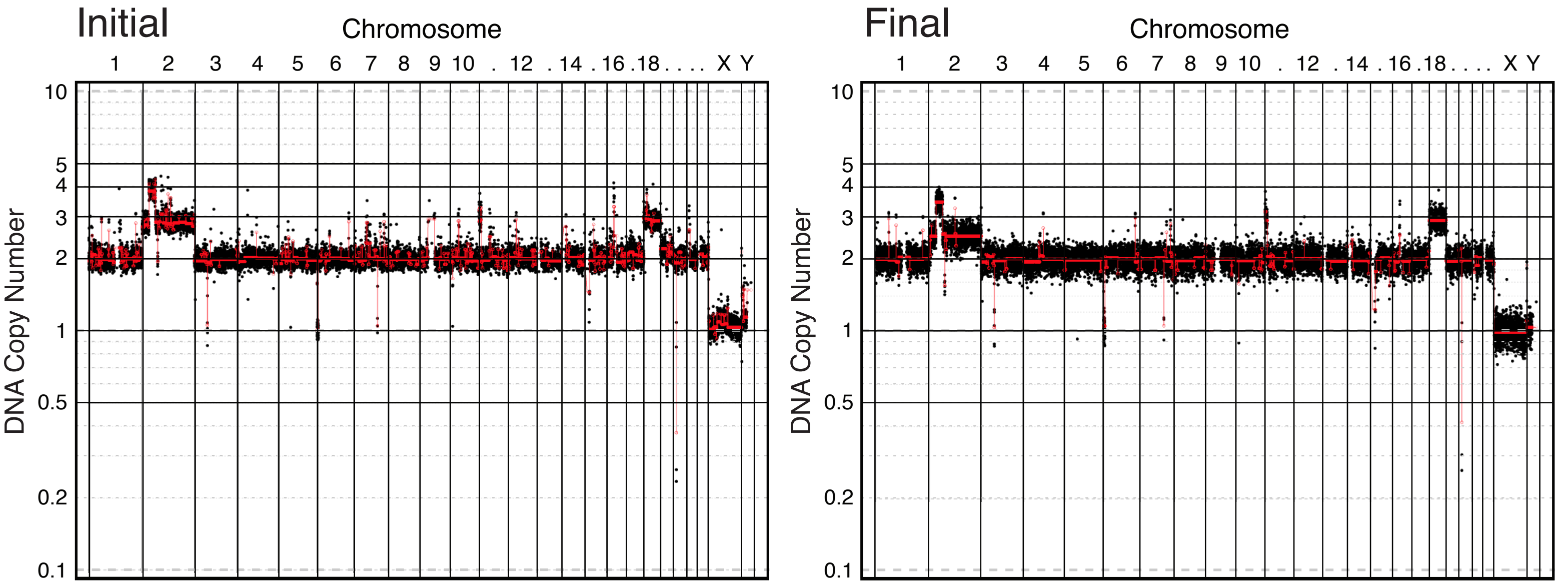

### DG: DLD1 Control 3

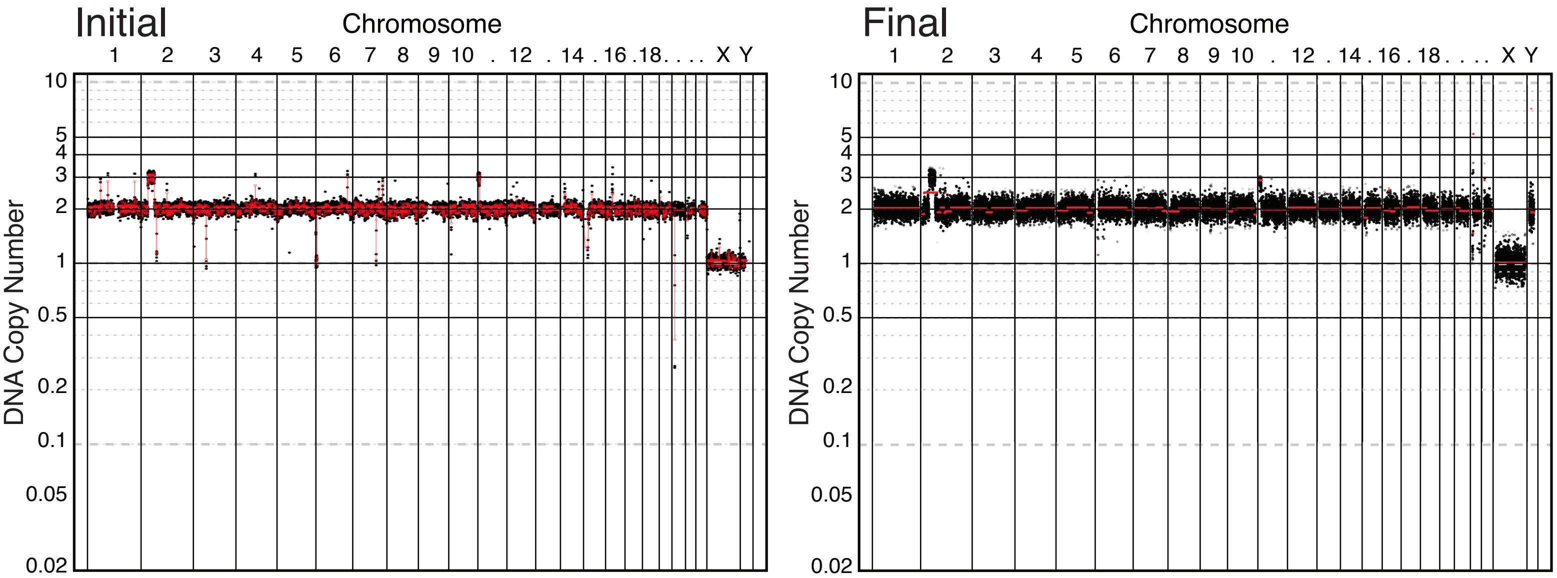


### DG: DLD1 Ts10.21

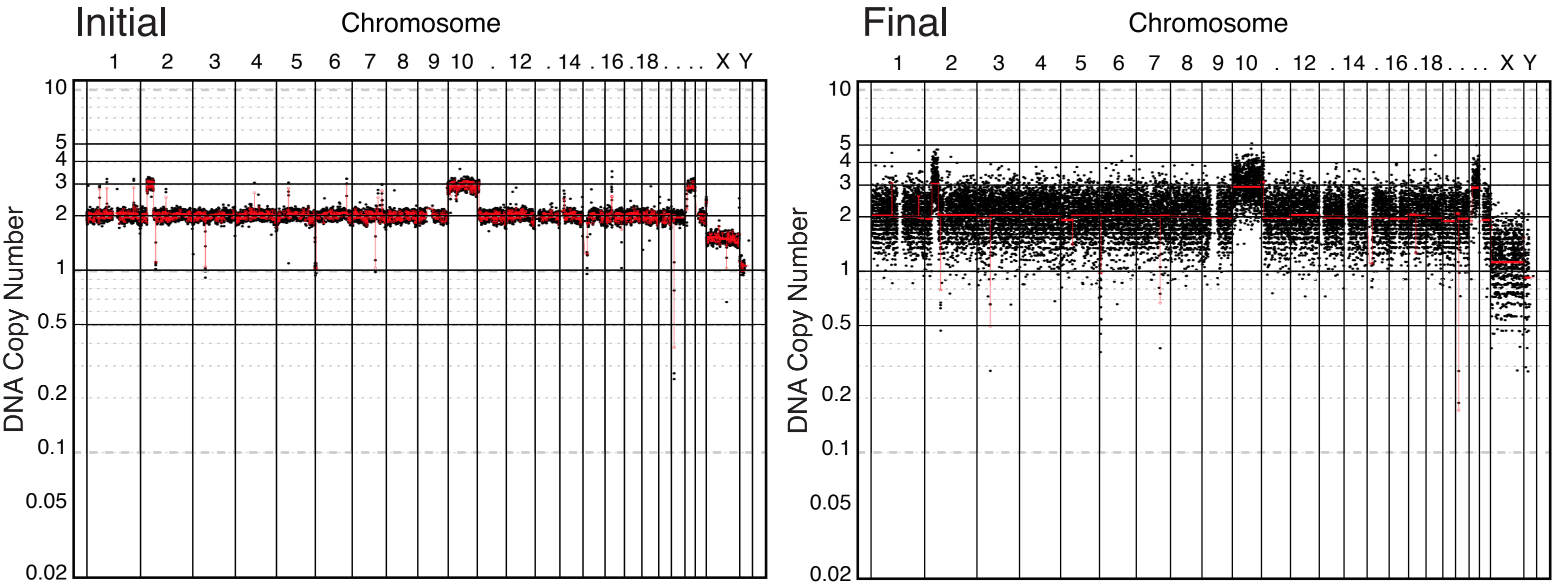


### DG: DLD1 Ts5.15

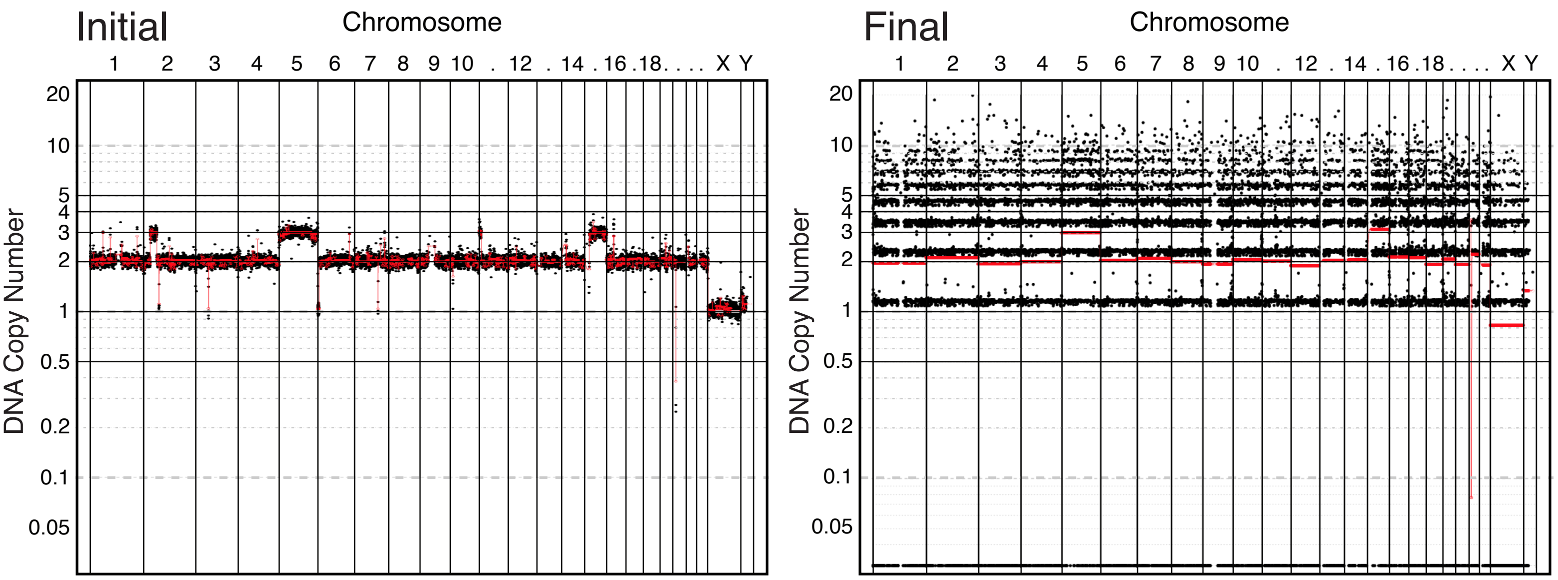

### DG: DLD1 Control 4

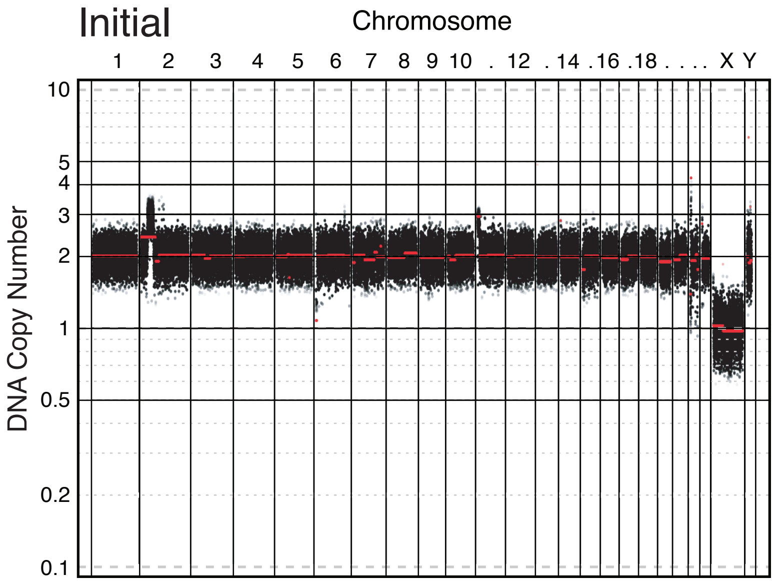


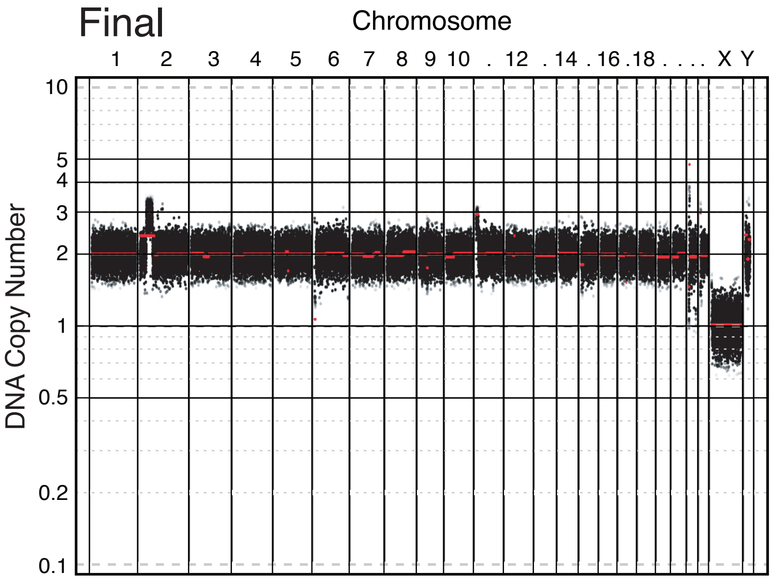


### DG: DLD1 Ts2.18 replicate 2

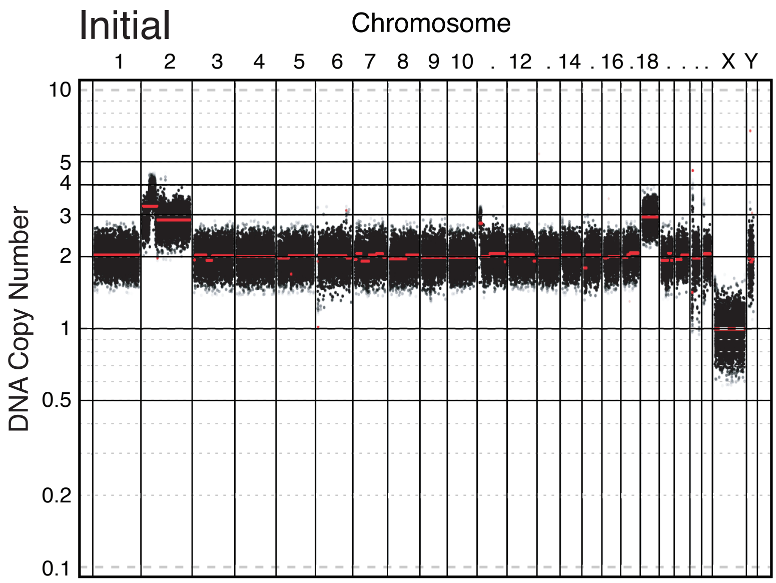


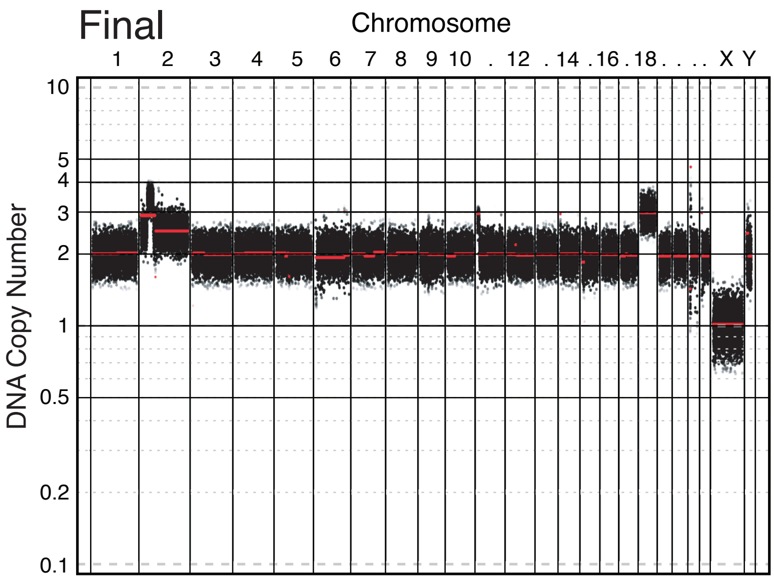


### DG: DLD1 Ts12.17

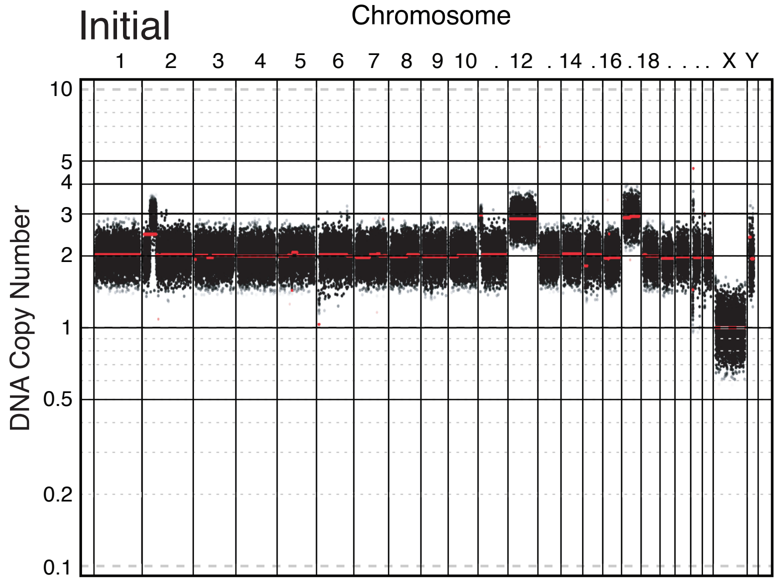


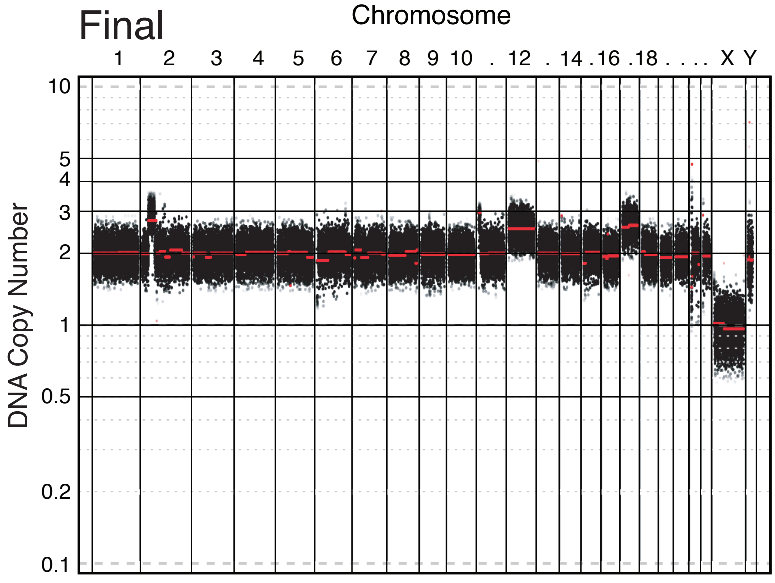

### DG: A2780 Control 1

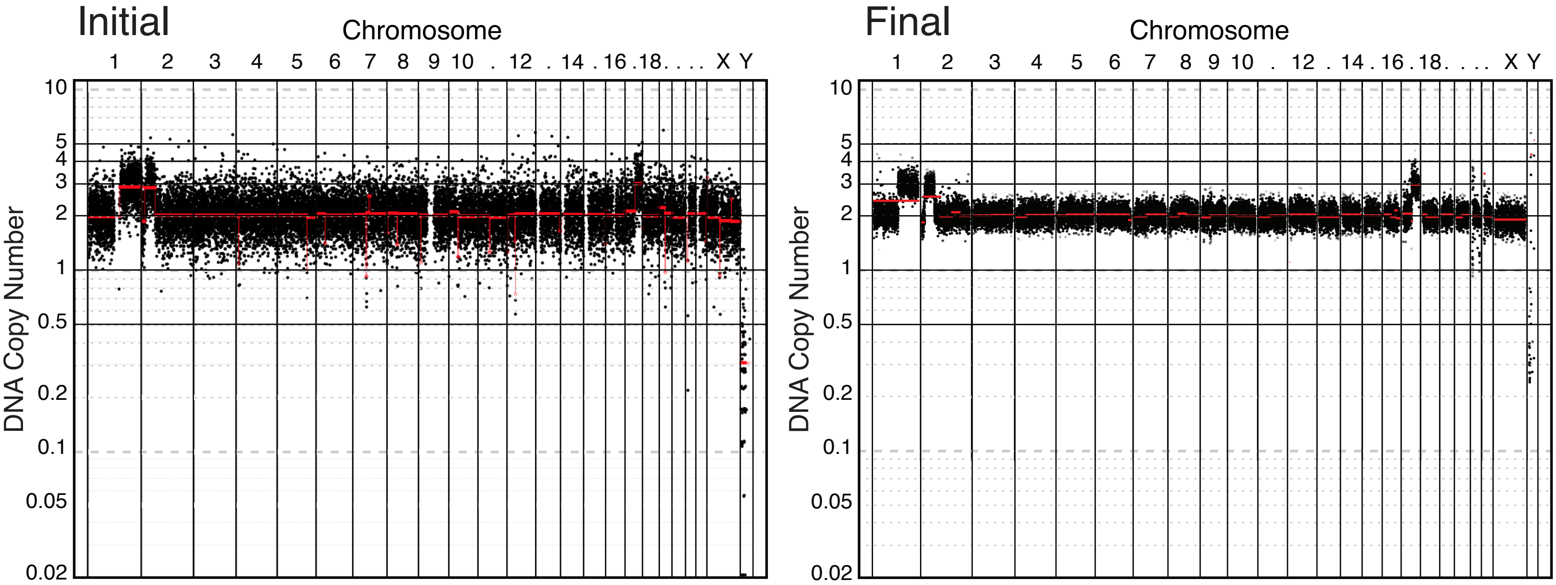


# DG: A2780 Ts2

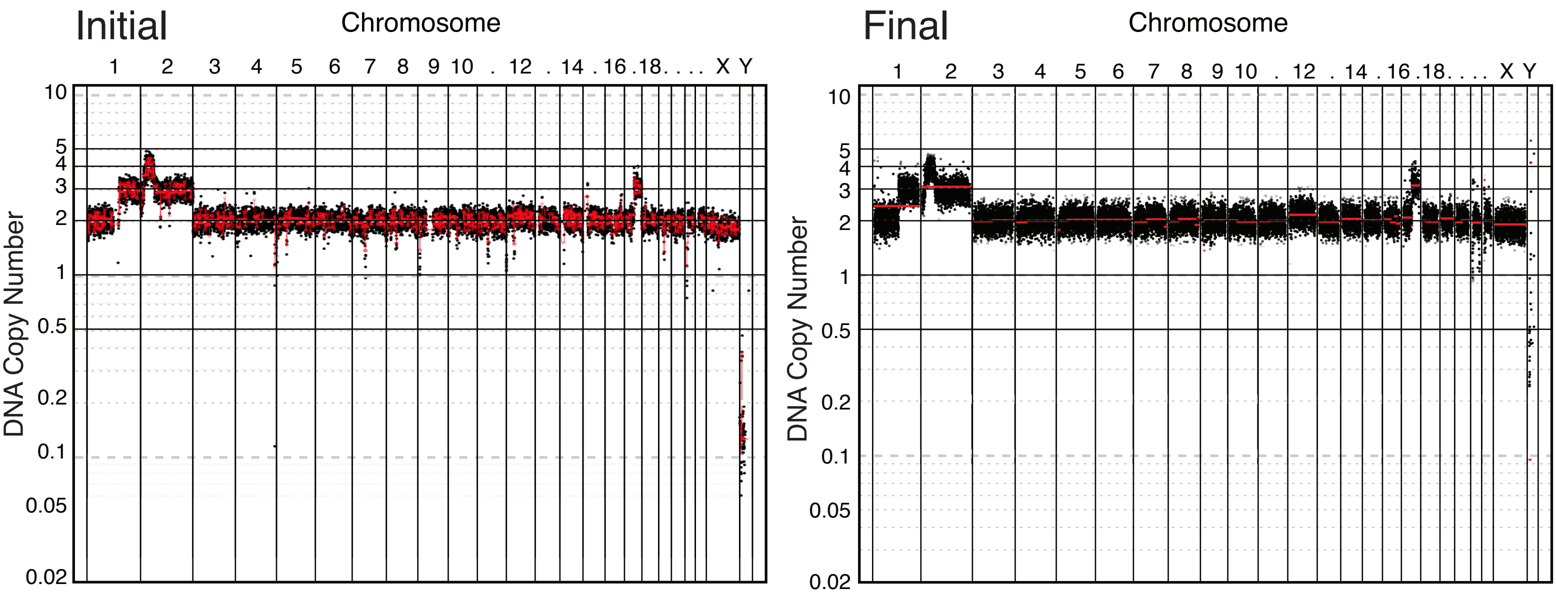


# DG: A2780 Ts10

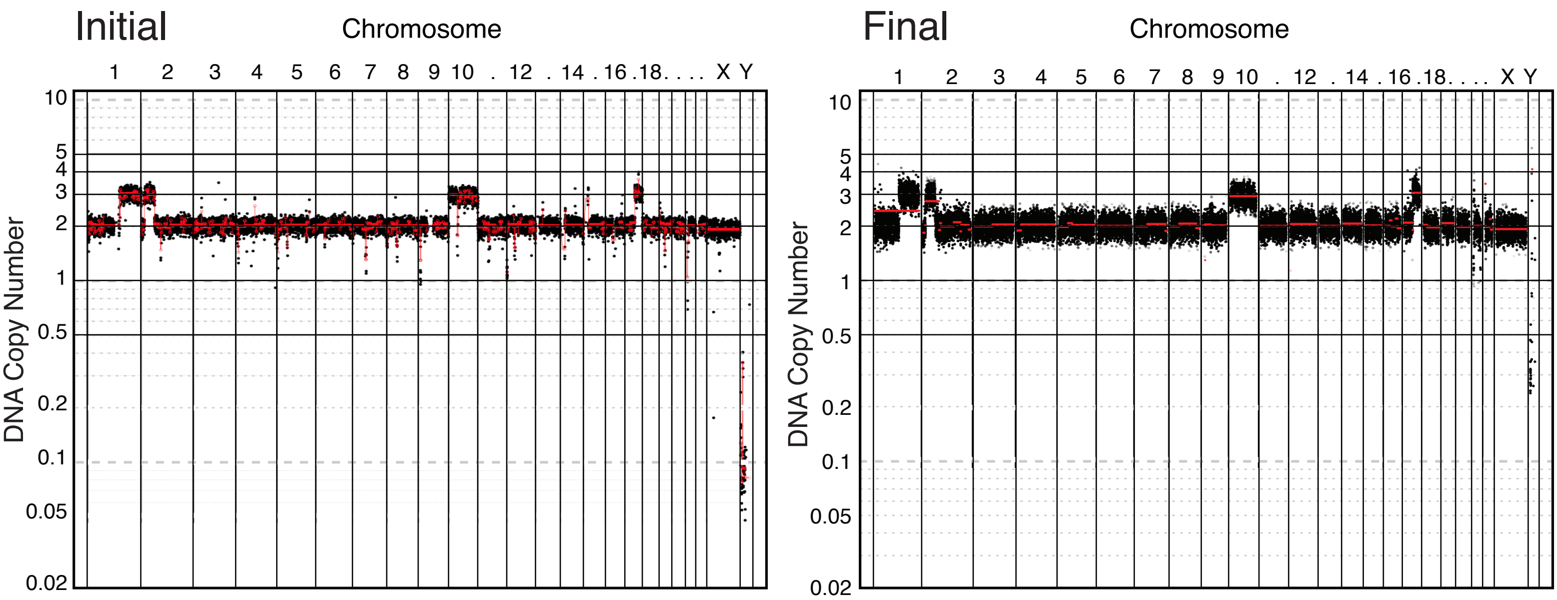

### DG: A2780 Control 2

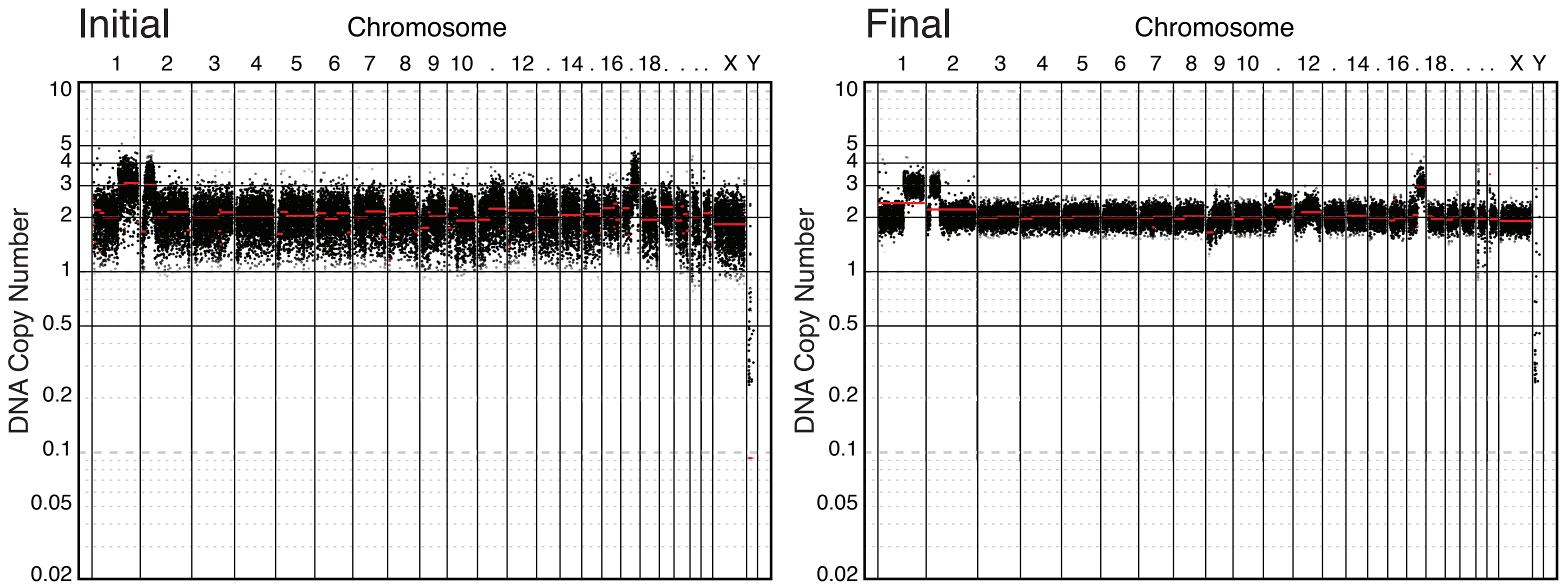


# DG: A2780 Ts8

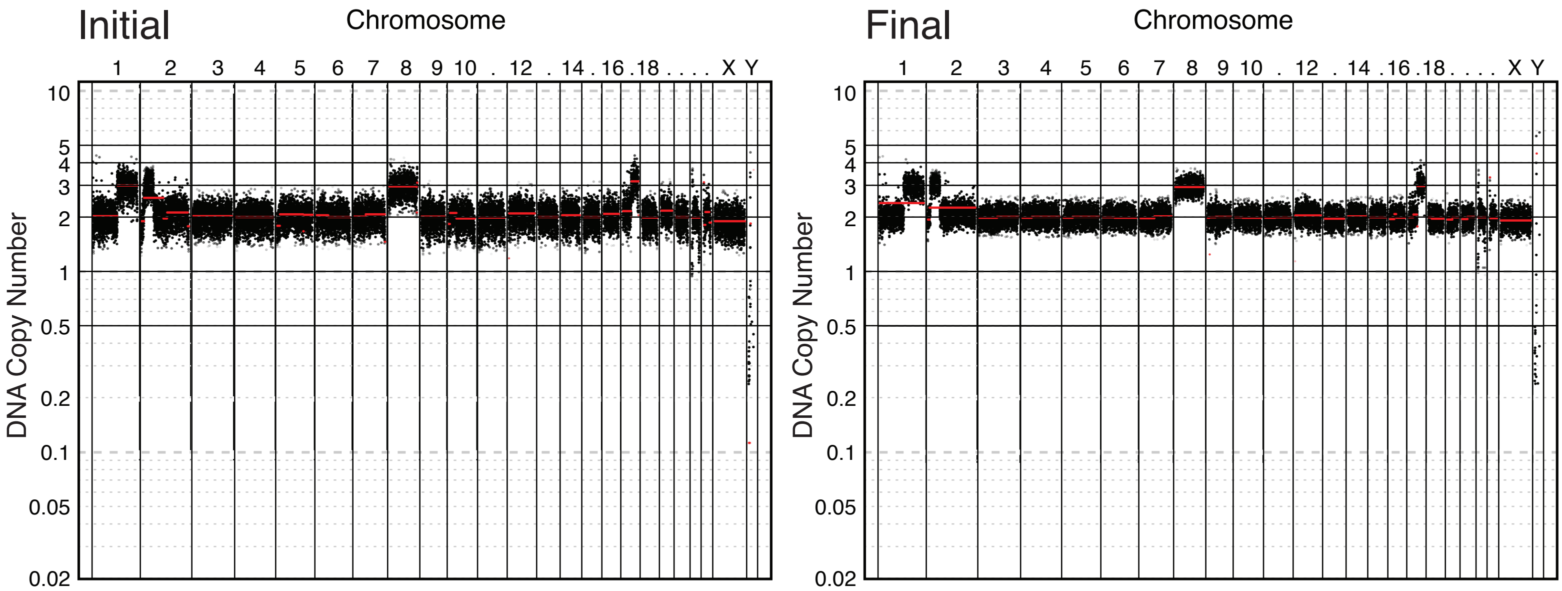


# DG: A2780 Ts18

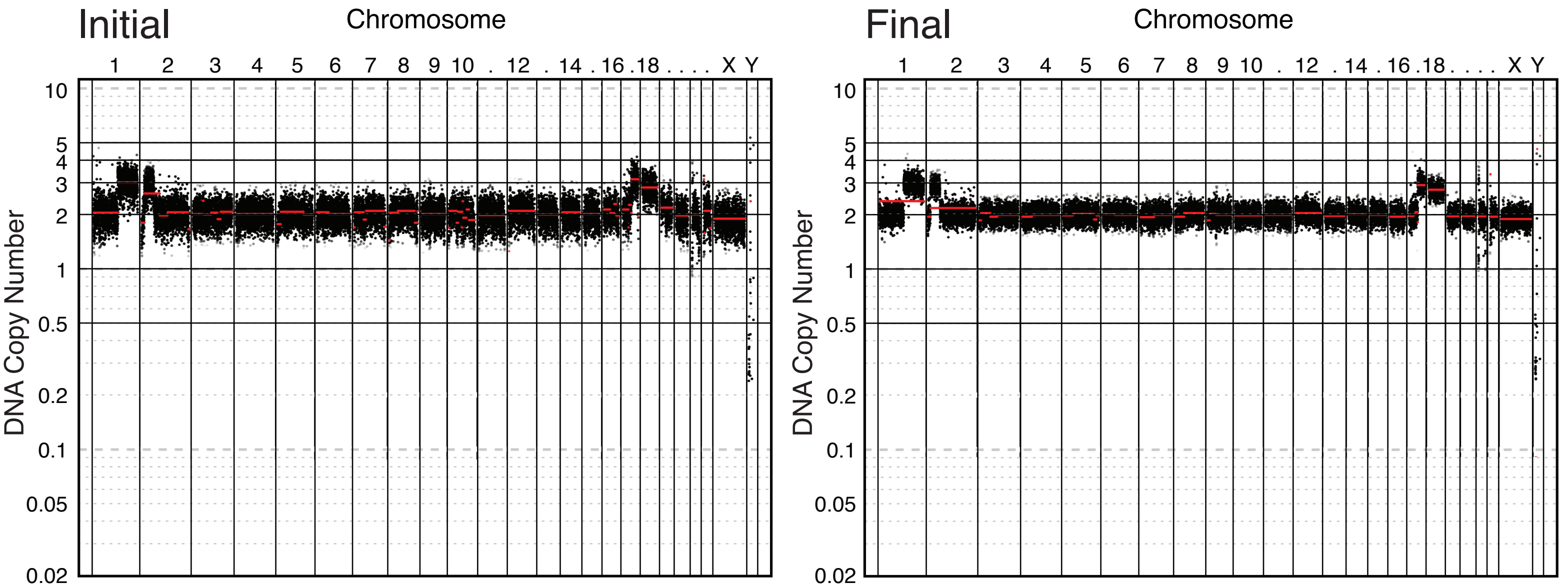

### DG: SNU1 control (c13)

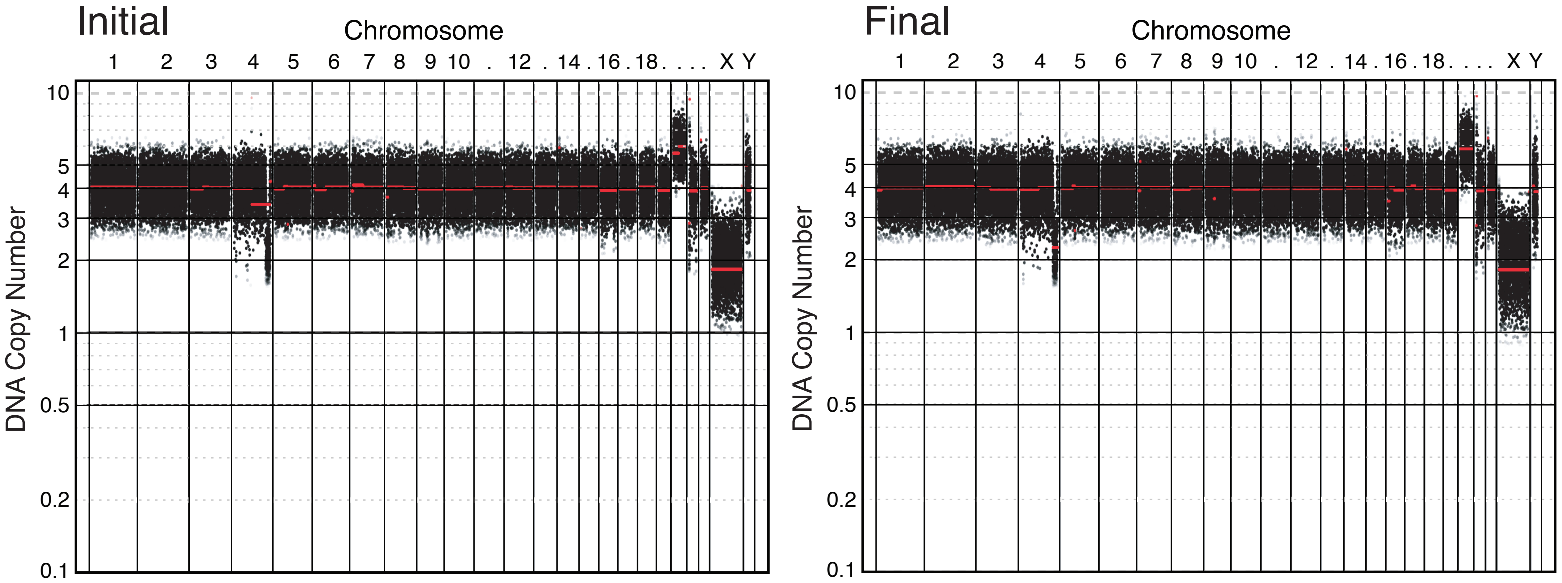


### DG: SNU1 Ts10.5.17_Ps2 (c12)

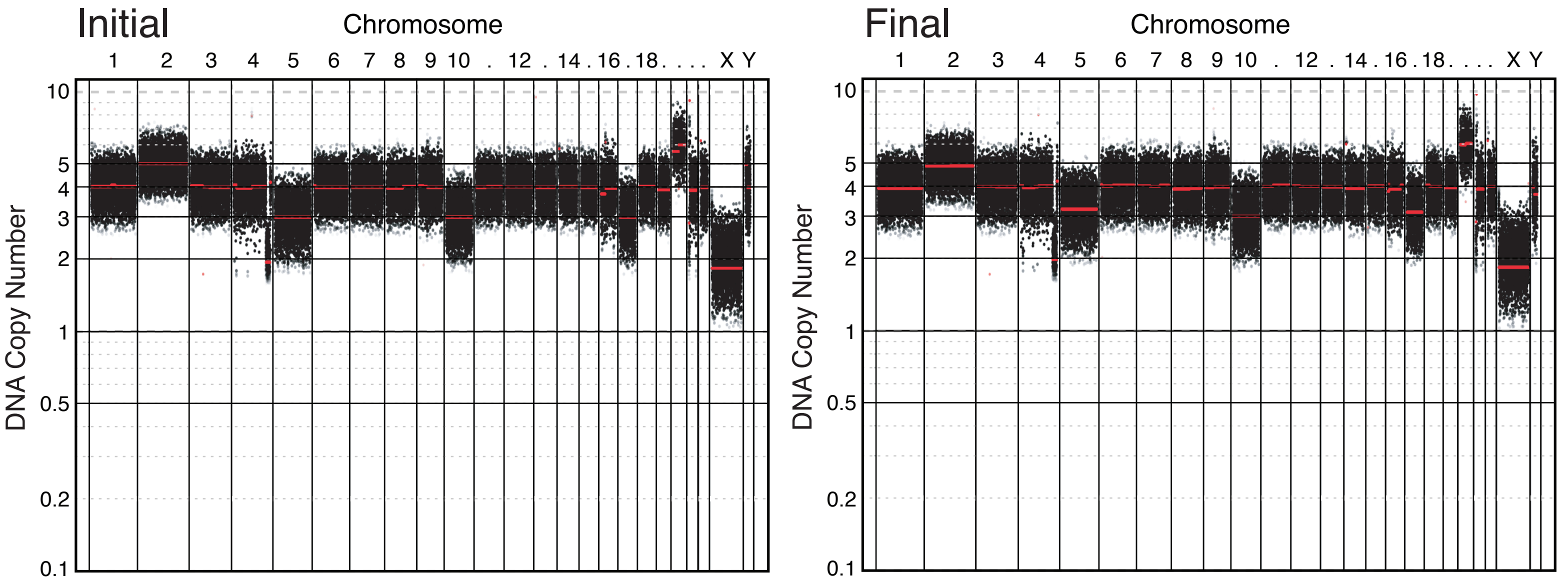


### DG: SNU1 DiY_Ts10.1.8.14_Ps5 (c24)

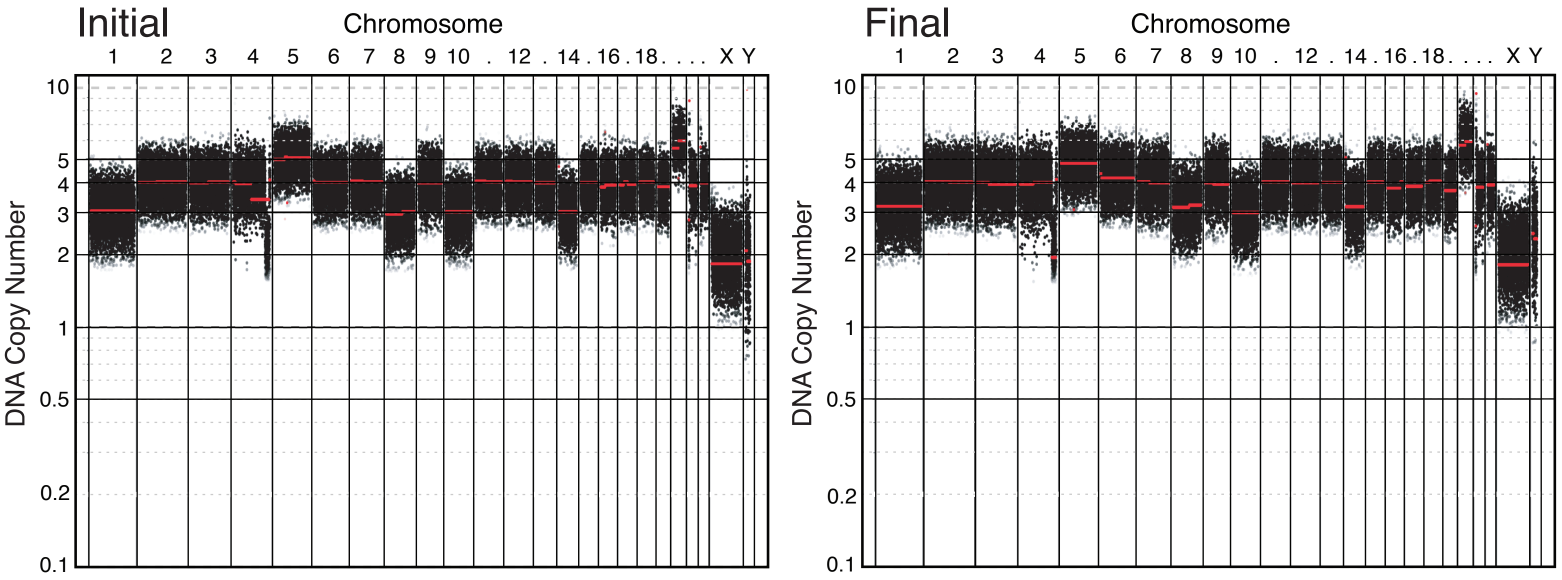

### DG: SNU1 Ts10.6q_Ps8.12 (c111)

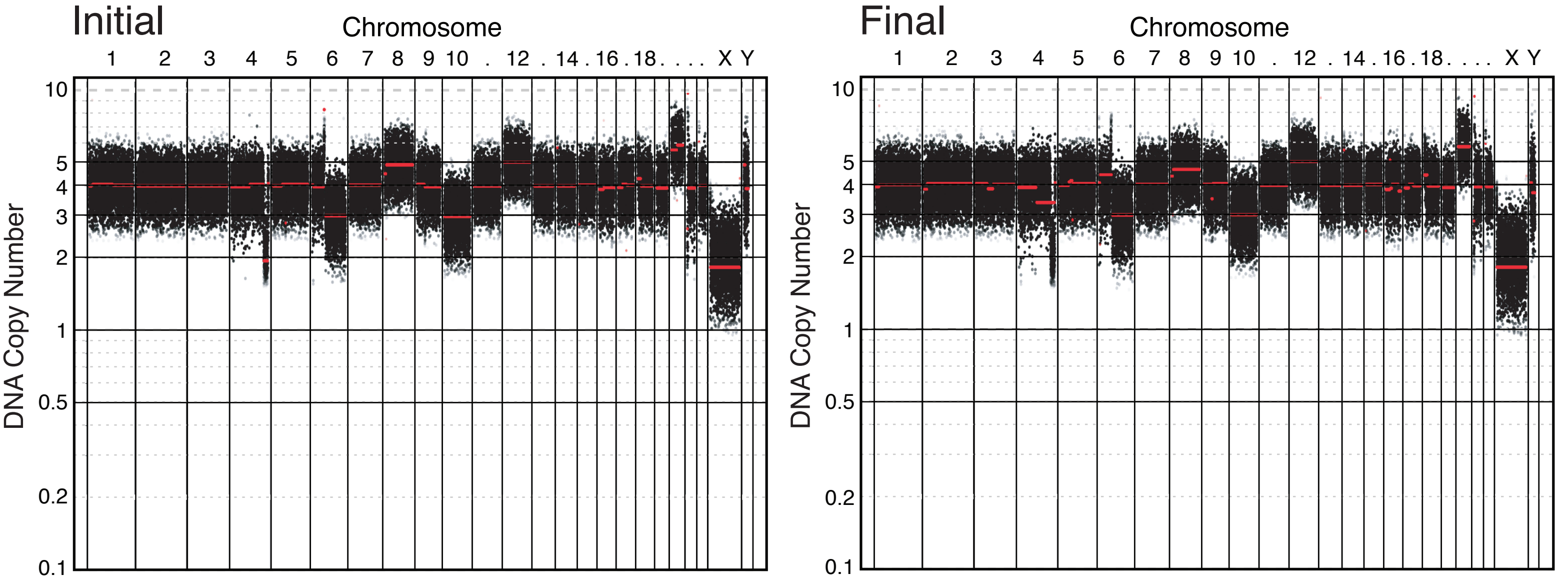


### DG: SNU1 Ts9.19 (c114)

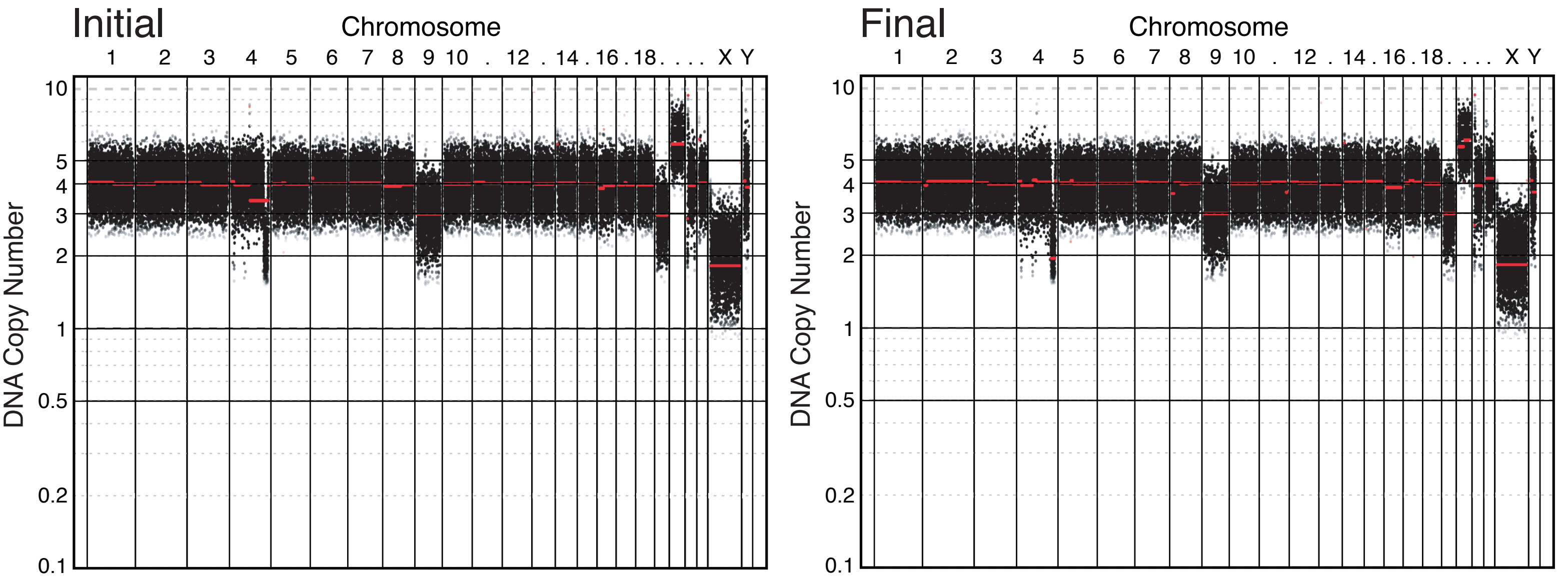

### WGL: DLD1 WT

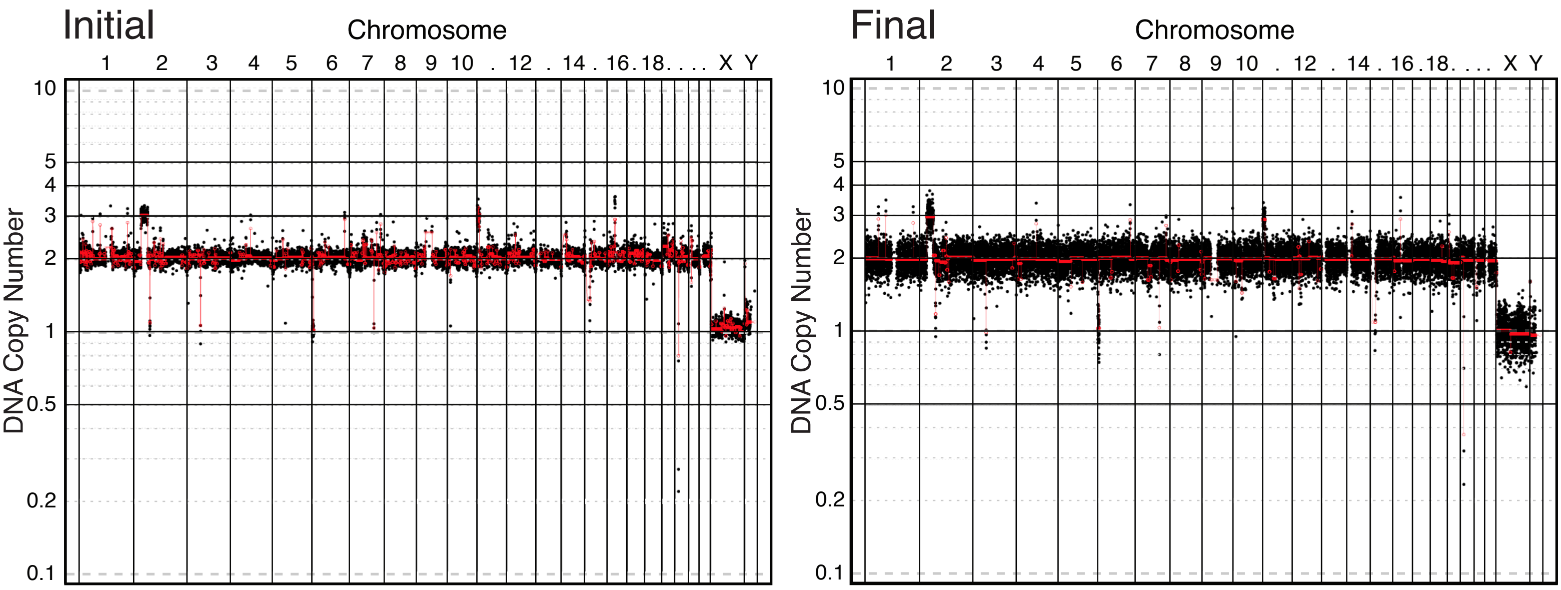


### WGL: DLD1 Ts13

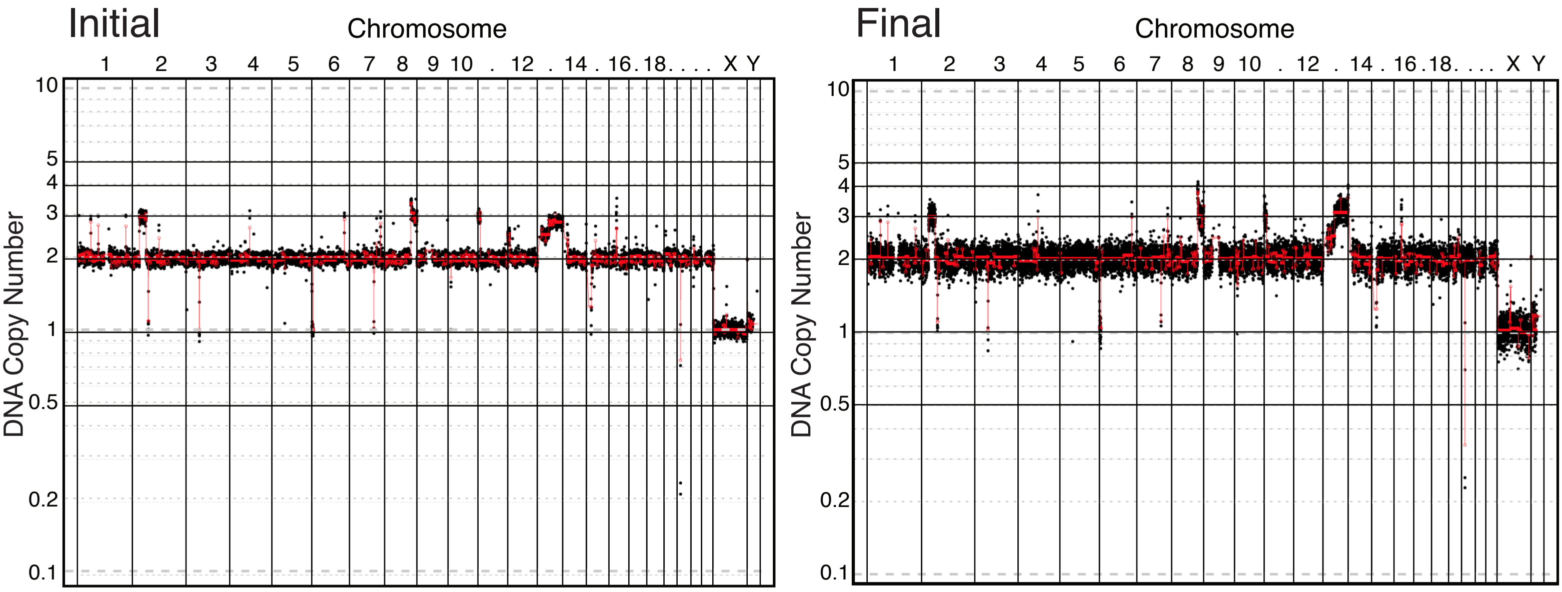

### WGL: Vaco432 WT

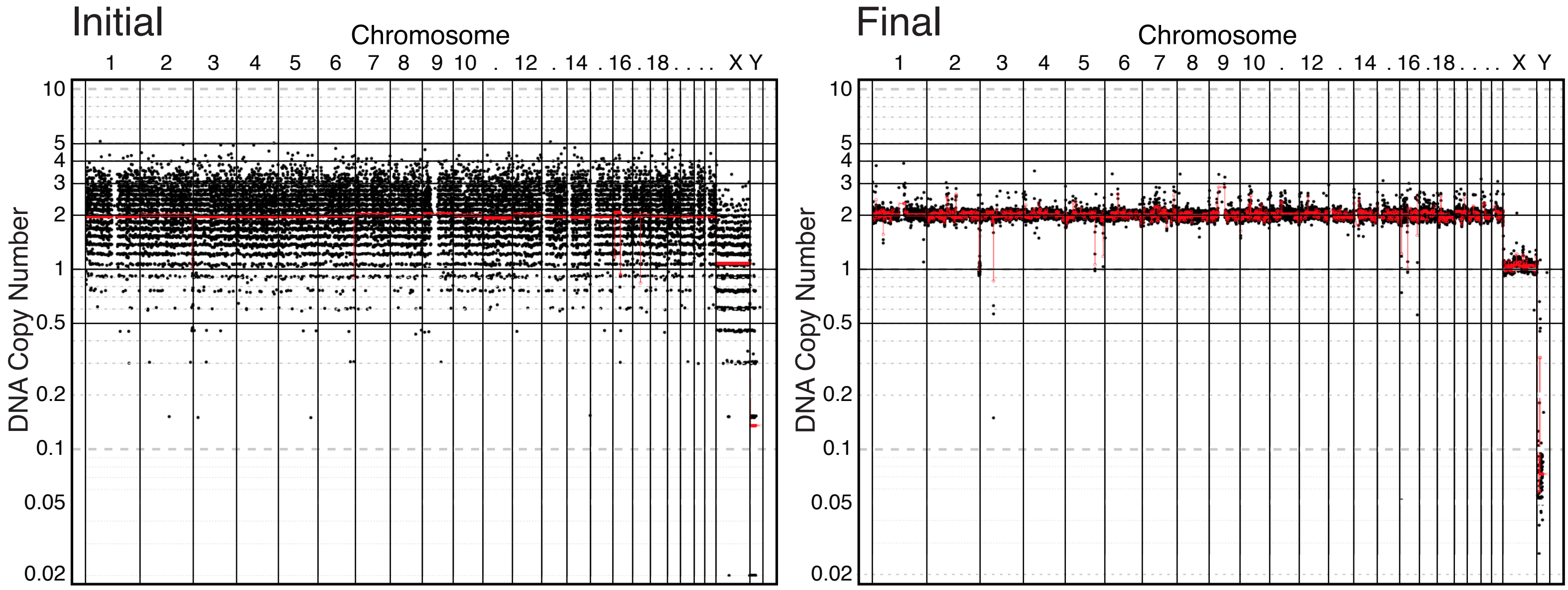


### WGL: Vaco432 Ts13

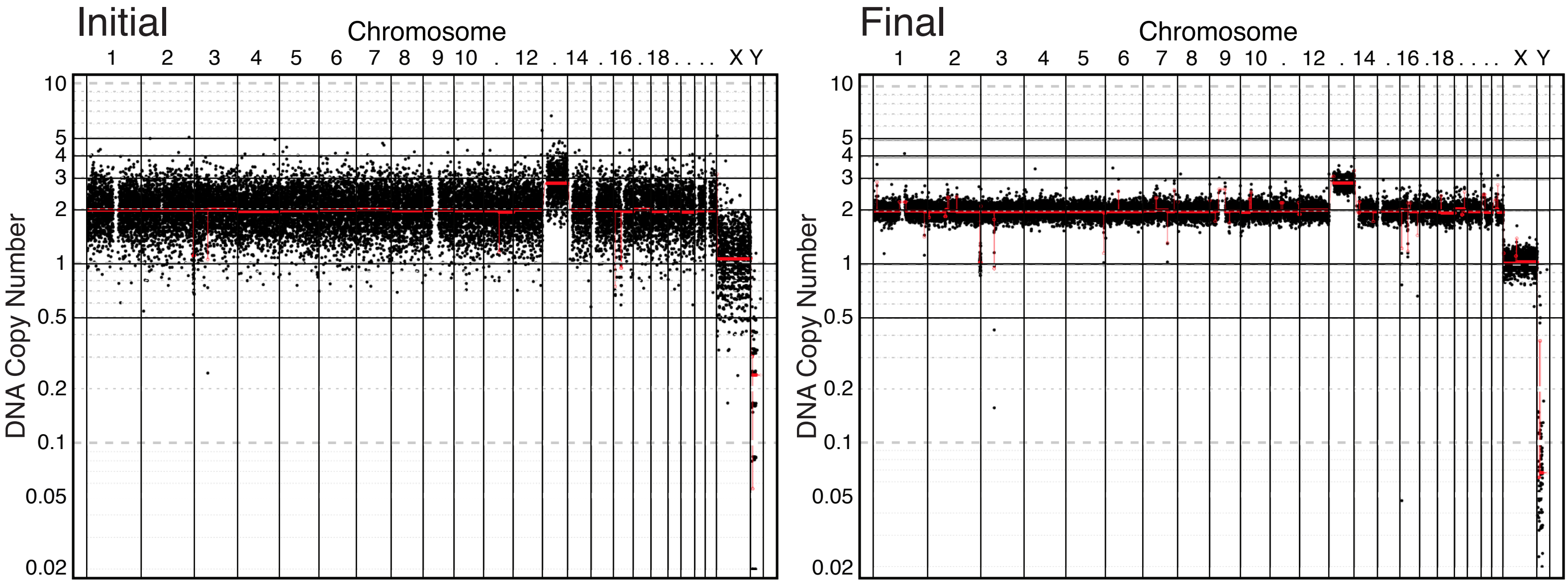

### WGL: A2780 WT

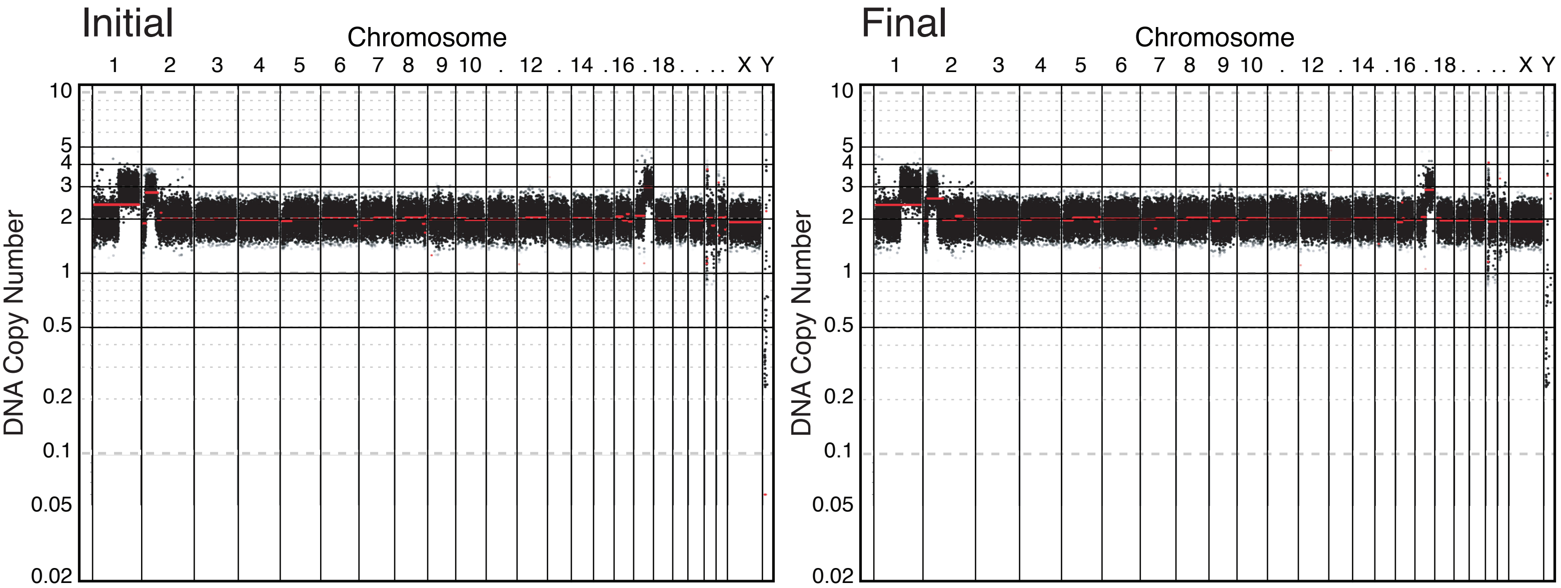


### WGL: A2780 Disomy 1q

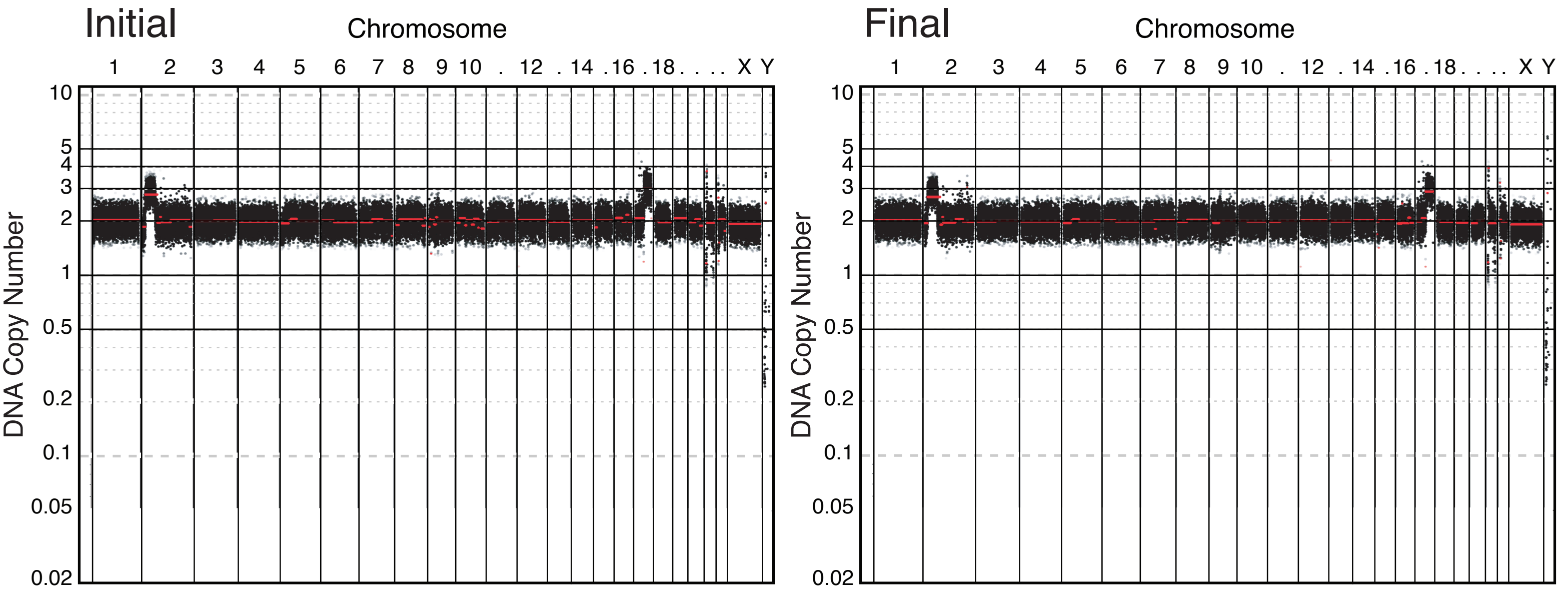

### WGL: A2058 WT

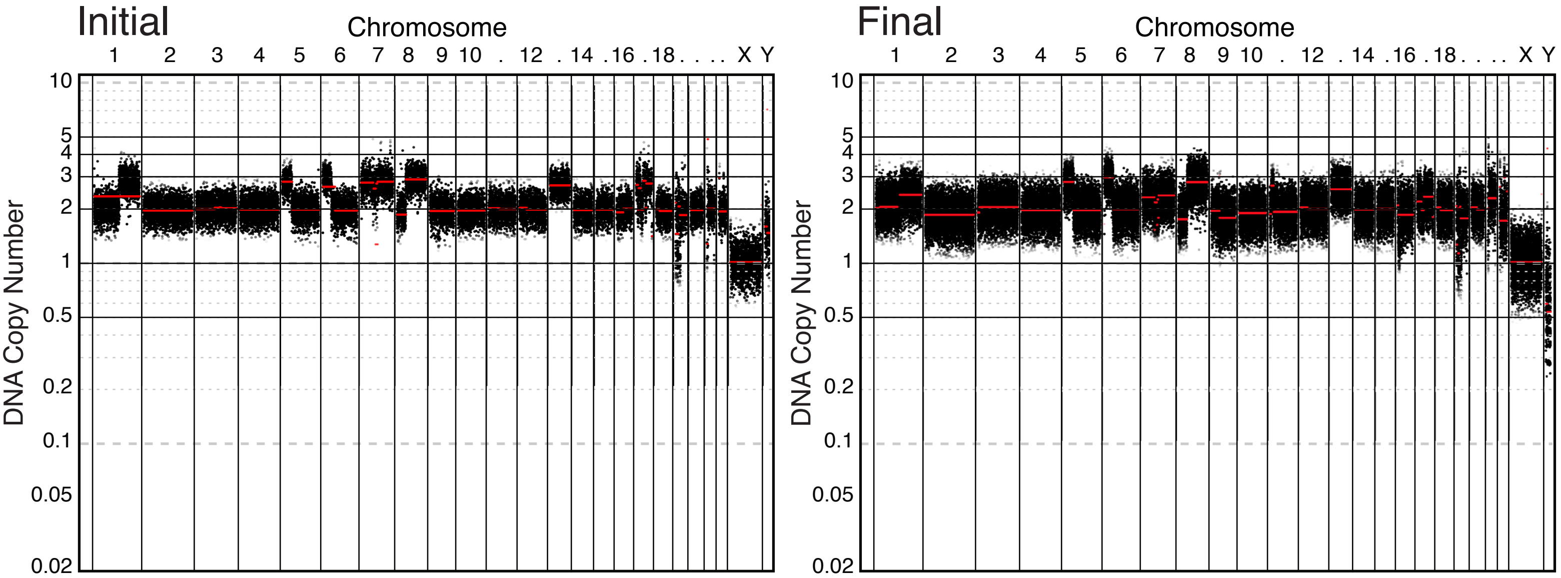


### WGL: A2058 Disomy 1q

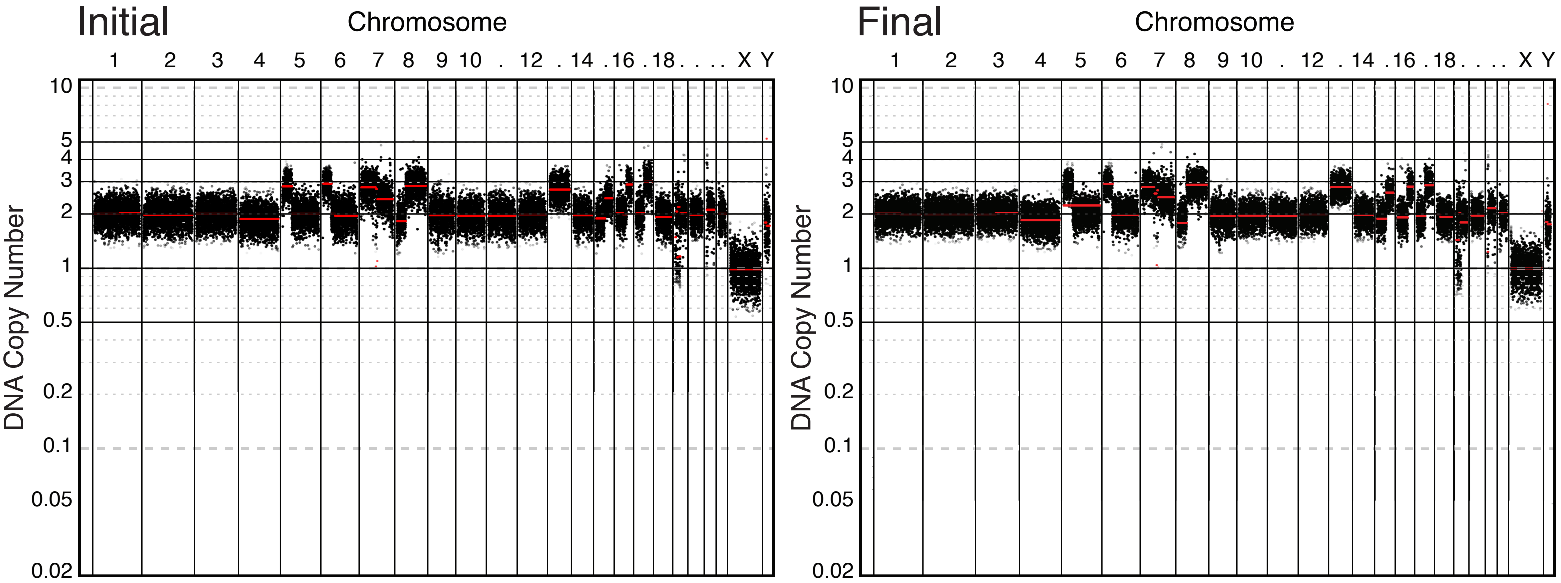


### WGL: A2058 Disomy 7p

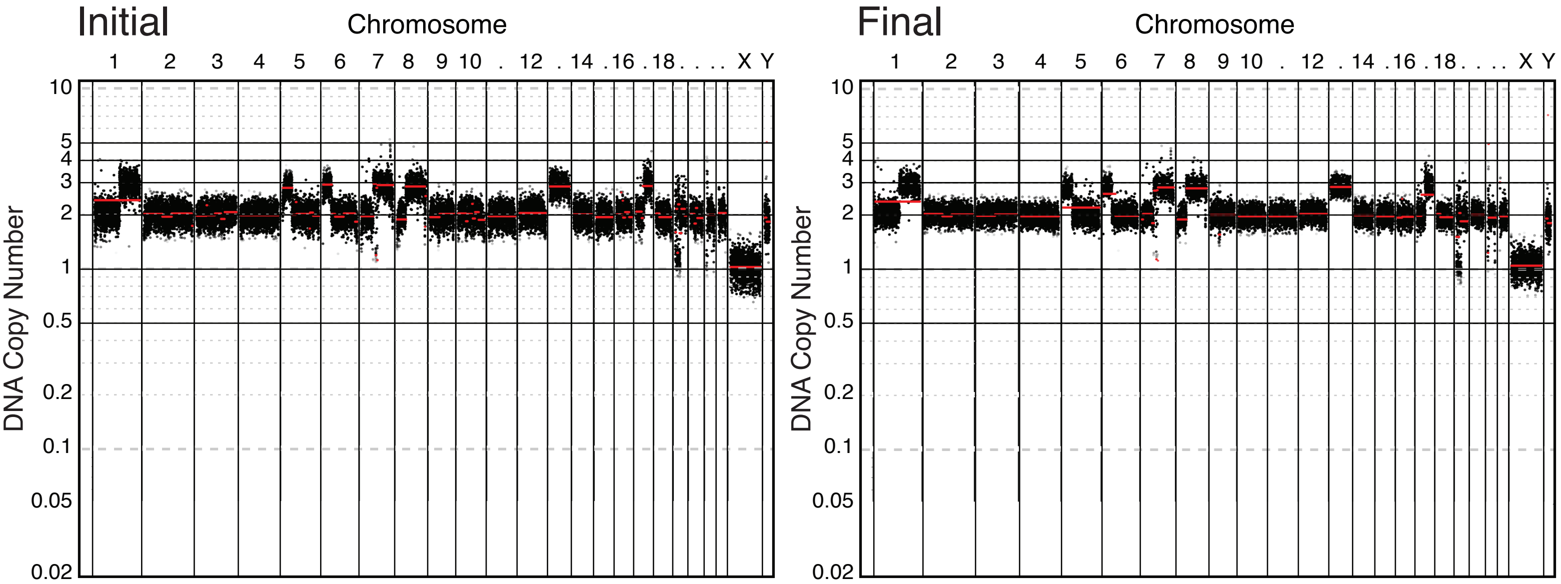

### WGL: AGS WT

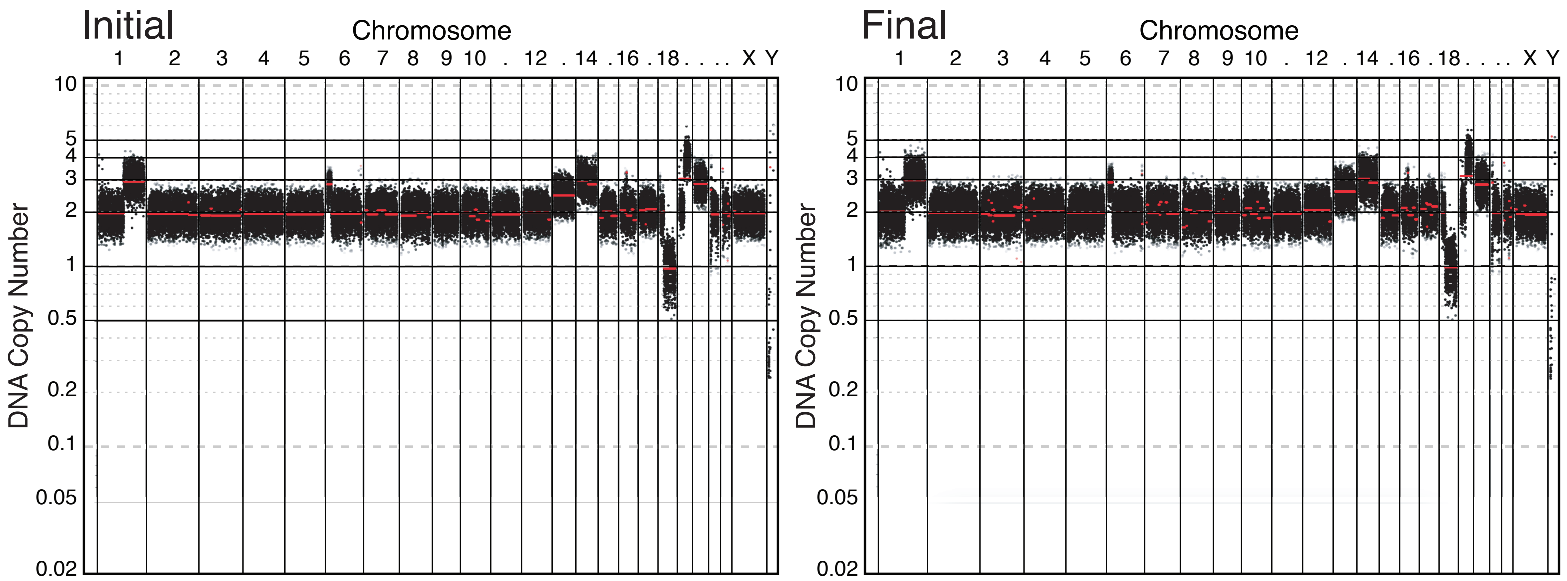


### WGL: AGS Disomy 1q

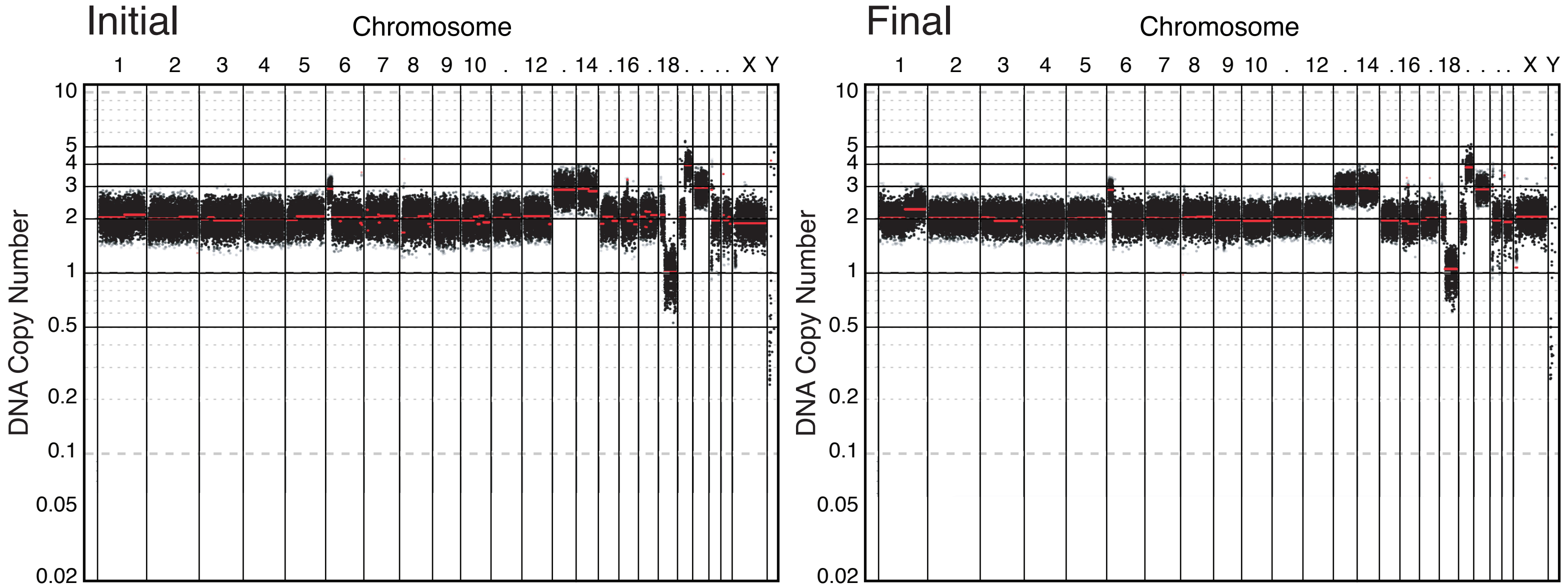

### WGL: MCF10A WT

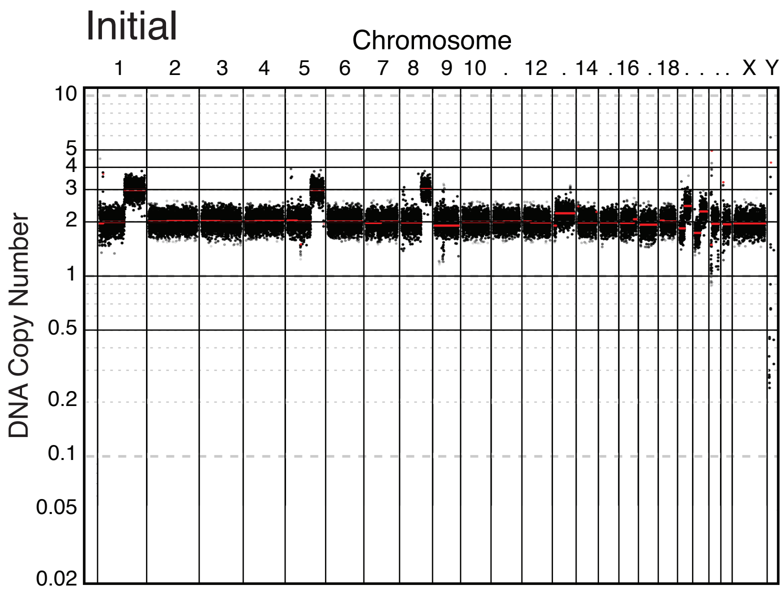


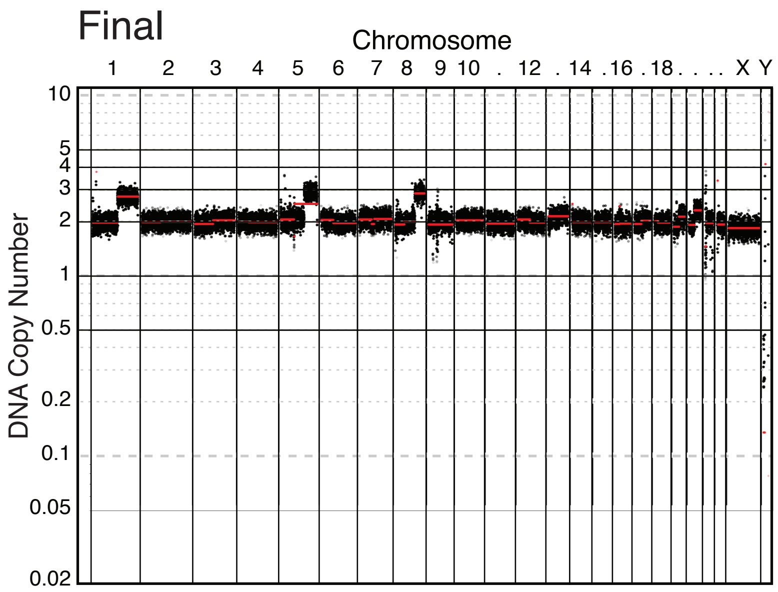


### WGL: MCF10A Disomy 1q

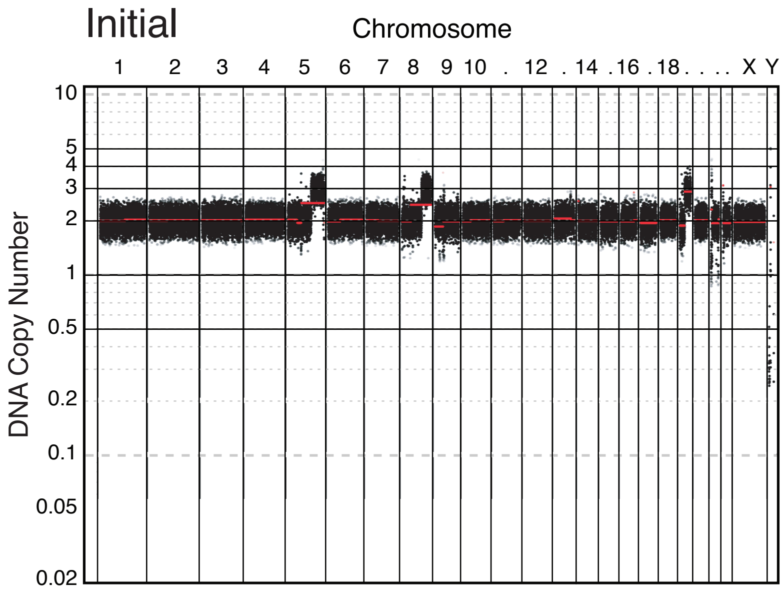


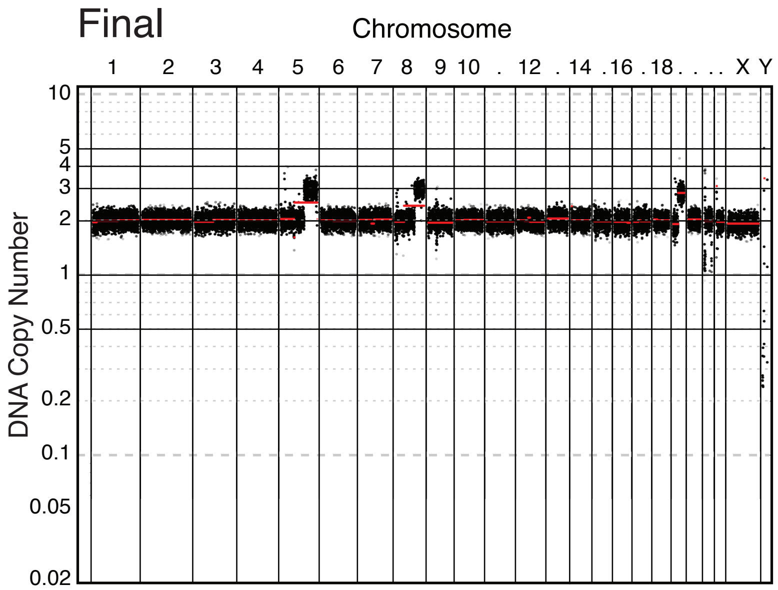
